# FlowMap: Geometry–Dynamics Consistent Embedding of RNA Velocity for Interpretable Cellular Trajectories

**DOI:** 10.64898/2026.09.21.753192

**Authors:** Jingyuan Hu, Harinder Singh, Jishnu Das, Luca Pinello, Rong Ma

## Abstract

Single-cell RNA sequencing reveals how cells vary across states, but these measurements capture only static snapshots of dynamic biological processes. RNA velocity addresses this limitation by estimating how gene expression is changing over time, providing directional information about cell state transitions. However, current approaches treat cellular state and dynamics separately, leading to representations that violate fundamental geometric consistency and obscure biological interpretation. Here, we present FlowMap, a framework that jointly models cellular states and their dynamics within a unified geometric representation. FlowMap simultaneously reconstructs a smooth low-dimensional manifold of gene expression and constrains RNA velocity to follow its local geometry, producing coherent and denoised representations of cellular trajectories. Across simulated and real datasets, FlowMap recovers interpretable dynamical patterns, including continuous progressions, branching events, cyclic behaviors, and stable-like states. It highlights key genes and gene programs involved in development that play distinct roles in these dynamics, and extends naturally to spatial transcriptomics, where it captures spatially organized developmental processes. Together, FlowMap establishes a principled geometric framework for modeling cellular dynamics, bridging representation learning and dynamical inference in single-cell analysis.

---

Single-cell RNA sequencing enables the characterization of cellular heterogeneity across diverse biological systems, including development, tissue homeostasis, immune responses, and the cell cycle. By profiling large populations of cells, it reveals structured variation in gene expression associated with continuous changes in cellular state. However, because these measurements capture only static snapshots, inferring the directionality of cell-state transitions and the dynamics of cellular lineage and fate decisions remains a fundamental challenge [1, 2].

RNA velocity, introduced by La Manno et al. [3], addresses this challenge by providing directed, local information about how cellular gene expression is changing over time. Rather than treating each cell as a static snapshot, RNA velocity estimates how a cell’s gene expression profile is shifting toward its immediate future state. Although originally derived from differences between unspliced and spliced mRNA abundances, the concept more broadly encompasses approaches that leverage transcriptional kinetics, metabolic labeling, or related measurements to capture short-term transcriptional change [4, 5]. When applied across many cells, these estimates reveal coordinated patterns of progression that reflect underlying cellular dynamics and enable the identification of continuous transitions, branching processes, and cyclic behaviors [6, 7]. In addition to single-cell sequencing, RNA velocity complements trajectory-based analyses by adding directionality and temporal ordering to static single-cell measurements.

A common observation underlying many single-cell analyses is that cellular states occupy a low-dimensional structure within the high-dimensional space of gene expression, making low-dimensional embeddings central to interpretation and visualization [8, 9]. At the same time, cellular transitions are inherently continuous processes, whereas experimental measurements sample cells as discrete points along these transitions [10, 11]. Bridging this gap naturally calls for a representation that captures continuous change across state space rather than isolated transitions between individual cells.

A convenient way to describe continuous change is through a *vector field*, which assigns to each cellular state a direction and magnitude of change in gene expression. In this setting, individual-cell RNA velocity estimates provide local directions of change and may exhibit measurement noise and cell-specific variation. When cells are modeled collectively, these estimates give rise to a continuous vector field that captures smooth changes across neighboring states, with the direction at each point reflecting the average behavior of nearby cells [5] (Fig. 1A). Such a vector field offers a principled way to describe how cells move through state space, filling in the regions between observed cells and providing a coherent representation of cellular dynamics.

**Figure 1:**
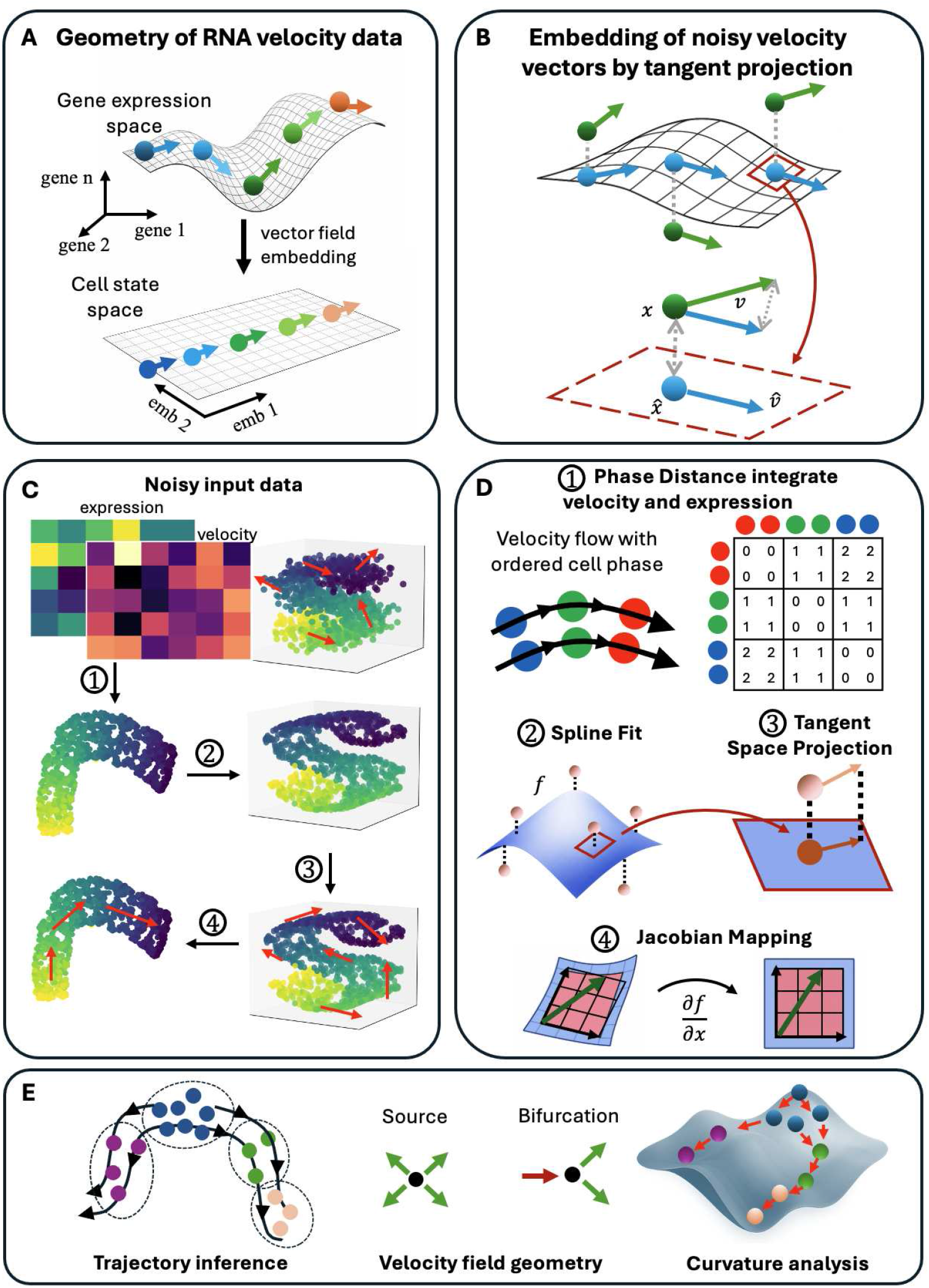
Overview of the FlowMap framework for geometry-preserving RNA velocity embedding. **(A)** RNA velocity defines local directions of change on a low-dimensional cell-state manifold embedded in gene-expression space. **(B)** FlowMap projects noisy velocity vectors onto the local tangent space of the learned manifold, producing geometry-consistent embedded dynamics. **(C)** FlowMap constructs a velocity-aware embedding and refines it into a smooth manifold representation with coherent vector-field structure. **(D)** The method proceeds through four steps: phase-distance construction, spline-based manifold fitting, tangent-space velocity projection, and Jacobian-based velocity mapping. **(E)** The resulting vector field supports trajectory inference, geometric characterization of flow structure, and curvature-based analysis of cellular dynamics.

For this description to be meaningful, the representation of cellular states and their short-term changes must be consistent: changes in gene expression should align with the structure of the state space itself, rather than pointing off the manifold into directions unsupported by the data [5]. In practice, however, RNA velocity vectors are estimated from sparse and noisy molecular measurements, and therefore contain components arising from sampling noise, model misspecification, or high-dimensional variation that is not supported by the underlying cellular manifold. Using these raw vectors directly can therefore introduce directions that are inconsistent with the geometry of the inferred cellular manifold. FlowMap addresses this by constraining observed velocities to the tangent space of the learned manifold, removing off-manifold components while jointly modeling gene expression and RNA velocity profiles at single-cell resolution within a vector-field-preserving manifold learning framework. Our approach integrates both modalities to generate low-dimensional cell embeddings together with coherent, cell-specific developmental directions, representing transcriptional dynamics as smooth vector fields defined on the inferred manifold (Fig. 1B-D).

Despite this intuitive geometric relationship between cellular states and their short-term changes, most existing RNA velocity analysis pipelines do not explicitly enforce geometric consistency between gene expression embeddings and inferred dynamics [3–5, 12]. In practice, low-dimensional representations are typically constructed from gene expression alone, while directional information is incorporated post hoc through local transition probabilities or neighborhood-based smoothing. Although these two-step approaches are effective for visualization, they treat cellular states and dynamics as separate objects. Without explicitly accounting for the intrinsic interactions between these two modalities, the reliability of such approaches critically depends on the initial expression-based cell embedding, potentially propagating errors and obscuring true developmental dynamics, particularly when reconstructing continuous flows or comparing global dynamical structures [5, 10]. Methods such as PAGA [13] partially address this limitation by preserving coarse-grained topological connectivity between cellular populations, while more recent approaches such as GraphVelo [14] improve local manifold consistency by projecting velocity vectors onto neighborhood-derived tangent spaces, highlighting the growing importance of geometry-aware velocity analysis. Deep learning frameworks such as deepVelo [15] and VeloAE [16] further learn latent representations of cellular dynamics, but these mappings are implicit and therefore do not provide an explicit coordinate description of the underlying manifold or vector field for direct geometric analysis. As such, existing methods generally lack an explicit, unified geometric representation of the developmental manifold that enables differential-geometric characterization of both cellular states and their dynamics. This mismatch highlights a gap between the conceptual interpretation of RNA velocity—as continuous change along a state manifold—and its practical representation in low-dimensional analyses.

To address this limitation, we introduce FlowMap, a framework that explicitly models the geometric relationship between cellular states and their short-term transcriptional changes. FlowMap takes gene expression and RNA velocity as joint inputs and is agnostic to the specific method used to estimate RNA velocity, allowing velocity estimates from any existing or future algorithm to be incorporated into the analysis. At its core, FlowMap learns a unified geometric representation by defining a smooth, differentiable mapping between a low-dimensional latent state space and the original gene expression space, while constraining the inferred velocity field to lie locally tangent to the learned cell-state manifold. In this way, variation in cellular state and local dynamical information are coupled within a common geometric framework rather than treated as separate representations. The resulting model provides a geometrically coherent view of cellular states and their dynamics, enabling RNA velocity to be interpreted consistently with the geometry of the underlying state manifold and supporting the characterization of developmental processes through quantities such as vector-field topology, acceleration, curvature, and spatial gradients. Importantly, the assumption that cellular states lie on a smooth manifold and that their dynamics are governed by a smooth vector field is already adopted, either explicitly or implicitly, by most existing approaches [4, 5, 14]. A key advantage of FlowMap is that this assumption is not left unchecked: the framework enables a goodness-of-fit analysis of the inferred velocity field to assess its adequacy for a given dataset, thereby allowing users to evaluate model compatibility and obtain more reliable inferential results with associated uncertainty quantification.

Through this geometric reconstruction of the vector field, FlowMap provides a unified framework for studying how cellular trajectories evolve across the state landscape. It enables the identification of dynamical features such as developmental origins, branching events, and regions where trajectories change direction. By linking these geometric properties to transcriptional programs, FlowMap helps reveal the molecular mechanisms associated with cell-state progression and fate commitment. The same geometric framework naturally extends to spatial transcriptomics, allowing transcriptional dynamics to be interpreted in the context of tissue organization.

## Results

### The role and limitations of low-dimensional embeddings for cellular dynamics

Low-dimensional embeddings provide the primary interface for interpreting single-cell data, enabling visualization of cellular organization, trajectories, and population structure [6, 7]. In practice, analyses of RNA velocity are almost always performed in this space, where both cellular states and their inferred dynamics are visualized and interpreted. Any approach to modeling RNA velocity must therefore operate within this representation while accounting for its geometric properties.

Classical manifold learning methods such as Isomap [17] and locally linear embedding (LLE) [18], together with more recent approaches including t-SNE [19] and UMAP [20], aim to represent high-dimensional cellular states within a low-dimensional space while preserving aspects of local or global geometric structure. However, these methods distort geometric relationships in the original gene-expression space, including distances, angles, and global metric structure [21] (Fig. S1). As a result, relative positions in the embedding are not intended to be interpreted quantitatively, and different choices of algorithms, parameters, or random initializations can yield qualitatively different layouts of the same data.

These limitations become especially pronounced when directional information such as RNA velocity is visualized on low-dimensional embeddings. RNA velocity is defined in the original gene-expression space, but interpreted in the embedding, creating a mismatch between where dynamics are computed and where they are visualized. As a result, the resulting velocity fields can exhibit distortions such as inconsistent local directions or apparent reversals of global trends (Fig. 1A). These artifacts arise in part because velocity is typically incorporated after the embedding has been constructed, using local smoothing, transition probabilities, or interpolation on k-nearest-neighbor graphs to convey directional trends (Fig. S2).

Because commonly used embeddings do not define an explicit or differentiable mapping back to gene-expression space, they do not provide a well-defined tangent space in which local velocity vectors can be interpreted as directions of change. Consequently, cellular states and their dynamics are treated as separate representations. This separation leads to embedding-dependent flow patterns, increased uncertainty in velocity directions, and inconsistencies across embeddings due to local neighborhood noise (Fig. S3). Although low-dimensional embeddings remain invaluable for visualization and intuition, these observations highlight the need for additional geometric structure to obtain a consistent representation of cellular dynamics.

### The need for geometric consistency between cellular states and dynamics

A consistent representation of cellular dynamics requires that cellular states and their short-term changes be modeled within a unified geometric framework. RNA velocity provides local estimates of how gene expression changes and can be interpreted as defining a vector field over the cell-state space [3, 5]. For this interpretation to be meaningful, the directions of change must be compatible with the geometry of the underlying state space. Geometrically, this requires that velocity vectors lie in the local tangent directions of the manifold defined by gene expression, ensuring that local changes are aligned with the global structure of cellular variation.

To achieve this, the embedding must be linked to the original gene-expression space through an explicit and differentiable mapping. Such a mapping defines a local tangent space at each point, allowing velocity vectors to be projected consistently between the two spaces. This provides a principled way to interpret RNA velocity in the embedding while preserving its geometric meaning in the original space. In this formulation, the embedding is no longer treated as an independent representation, but as a coordinate system for a smooth manifold that captures both cellular states and their dynamics.

Under this framework, RNA velocity can be incorporated directly into the geometry rather than added post hoc. Local directions of change are constrained by the manifold structure, and global flow patterns emerge from the integration of these local dynamics. This leads to a representation in which cellular trajectories are smooth, coherent, and stable, and in which both state and dynamics are described consistently within the same geometric model (Fig. 1B).

### Design principles of FlowMap for geometry-consistent velocity embedding

FlowMap embeds cellular states and RNA velocity within a single, coherent representation rather than treating them as separate objects (Methods, “Overview of the FlowMap framework”). It takes gene expression and the corresponding RNA velocity vectors as inputs, with kinetic parameter estimation handled by the upstream framework, and focuses on how these local transcriptional changes are organized across the cell-state space. By reconstructing the embedding as a smooth manifold with an explicit mapping to gene expression, FlowMap ensures that inferred velocity vectors follow the geometry of the underlying state space (Fig. 1C). In this way, the embedding is not only a visualization of cell states, but also a coordinate system in which their dynamics can be interpreted consistently. Since FlowMap operates directly on gene expression and the corresponding RNA velocity vectors, it is independent of the upstream velocity estimation procedure and can be applied uniformly to velocity fields inferred by existing methods.

First, FlowMap incorporates directional information directly at the embedding stage by introducing a phase distance that captures the relative, local progression of cells along the RNA velocity flow (Fig. 1D; Methods, “Velocity-aware distance metric”). This distance is computed by locally aligning neighboring cells along their velocity directions using a least-squares formulation, providing an estimate of separation along the underlying dynamical trajectory. This phase represents a cell’s relative position along the local RNA velocity flow. Cells with similar phase occupy nearby positions along the same trajectory, whereas larger phase differences indicate greater separation in their relative ordering. Because phase is estimated only from local neighborhoods, it provides a local measure of ordering along the flow. Unlike standard embeddings constructed solely from expression similarity, this velocity-aware distance distinguishes cells that are transcriptionally similar but positioned at different stages of a transition. As a result, the embedding more clearly reflects progression along continuous processes, producing layouts in which trajectories aligned with the vector field are emphasized and directionality is directly visible (Fig. S4, Fig. S5). Across synthetic dynamical systems, incorporating phase distance improved neighborhood preservation, vector field smoothness, and directional consistency relative to embeddings constructed without phase weighting (Fig. S6; Table. S2).

Second, FlowMap reconstructs the embedding as a smooth, parametric surface using a spline-based model [22], related to classical approaches for smooth manifold reconstruction such as principal curves and nonlinear manifold learning [17, 18, 23]. This yields an explicit and differentiable mapping between embedded coordinates and gene expression space (Fig. 1D; Methods, “Manifold reconstruction”). The reconstruction acts as a principled denoising step: by enforcing smoothness across the embedding, the model reduces cell-to-cell noise while preserving coherent large-scale structure in the data (Fig. S7). At the same time, it provides a global description of the data that remains consistent across the entire embedding, rather than relying only on local neighborhood relationships (Fig. S1).

Crucially, the explicit spline mapping defines a smooth manifold together with a well-behaved tangent space at every embedded point, supplying the geometric structure needed to embed RNA velocity consistently. RNA velocity vectors are projected onto the tangent space of the reconstructed manifold (Methods, “Tangent space and velocity projection”), ensuring that inferred transcriptional changes follow directions supported by the embedded state space rather than cutting across it (Fig. 1D). This projection enforces consistency between cellular states and their instantaneous dynamics by construction. The resulting representation allows RNA velocity to be interpreted as a vector field defined on a smooth low-dimensional state space, enabling downstream geometric analyses such as trajectory inference, identifying the geometric structure of the velocity field, and curvature-based characterization of cellular dynamics (Fig. 1E).

### Benchmarking FlowMap on simulated and real single-cell data

To evaluate the ability of FlowMap to recover underlying dynamical structure, we performed benchmarking on both synthetic vector fields with known ground-truth dynamics and real single-cell datasets (Methods, “Benchmark”). We benchmarked FlowMap against a diverse set of existing embedding and trajectory visualization methods commonly used for dynamical single-cell analysis [4, 5, 14, 24]. In simulated systems spanning linear, branching, and cyclic flows, FlowMap produces embeddings that preserve the geometry of trajectories and maintain alignment between local velocity directions and global progression (Fig. S8). These qualitative observations are supported by quantitative metrics, where FlowMap consistently achieves higher directional cosine similarity and smoothness, along with improved trustworthiness across a range of dynamical regimes (Table. S3). In particular, gains in magnitude correlation and cosine similarity are most pronounced in nonlinear and branching systems, indicating improved recovery of complex dynamical structure.

We further assessed performance on representative single-cell datasets exhibiting diverse dynamical regimes through two complementary comparisons. In the first benchmark, we fixed the cell-state embedding and compared only the resulting velocity embeddings, where FlowMap yielded more coherent and stable projected vector fields than alternative velocity visualization strategies (Fig. S9). In the second benchmark, we compared the full analysis procedure, including both cell-state embedding and velocity embedding, and found that FlowMap better preserved the coupling between manifold geometry and directional dynamics across datasets (Fig. S10).

Importantly, FlowMap is agnostic to the choice of upstream velocity estimation method and can be applied uniformly to velocity fields derived from different models, including stochastic, dynamical, and dynamo-based approaches. By projecting the learned low-dimensional vector field back into gene-expression space, FlowMap quantifies velocity-field consistency using the cell-level relative error between predicted and observed gene velocities (Fig. S11). This reconstruction residual measures how well a given RNA velocity estimate can be represented by a smooth vector field tangent to the learned cell-state manifold. It therefore serves two complementary purposes: it provides a common geometric criterion for systematically comparing upstream RNA velocity estimation methods, and it acts as a goodness-of-fit diagnostic for the underlying modeling assumption that cellular dynamics are well described by a smooth manifold and an associated tangent vector field. The residual profiles were dataset dependent: Dynamo showed the lowest median relative error in the Dentate Gyrus dataset, whereas scVelo stochastic achieved the lowest median relative error in the Larry dataset (Fig. S11C). These differences demonstrate that no single velocity estimation method is uniformly most compatible with the geometry of all datasets and illustrate how FlowMap can identify, on a dataset-specific basis, which velocity estimates are most consistent with the inferred state-space structure. More broadly, this analysis reveals systematic differences across methods in both fidelity and stability, providing a principled quantitative framework for evaluating RNA velocity estimates and assessing model adequacy beyond qualitative visualization.

Together, these results demonstrate that enforcing geometric consistency between cellular states and their associated dynamics not only improves embedding quality, but also provides a unified framework for evaluating and comparing velocity estimation methods.

### FlowMap recovers RNA velocity field geometry and gene-level consistency in cell-cycle data

To evaluate whether FlowMap preserves the geometry of RNA velocity–defined vector fields and enables identification of genes whose dynamics are consistent with this geometry, we analyzed a cell-cycle dataset with experimentally defined temporal structure. Specifically, we used the scEU-seq dataset of RPE1-FUCCI cells [25], in which cells are fluorescently labeled by cell-cycle phase and follow a well-characterized sequential progression, providing an external reference for ordering and directionality.

FlowMap recovers a cyclic embedding that aligns with the annotated cell-cycle phases using a curated set of 33 known cell-cycle genes [26] (Fig. 2A). All three methods recover the overall cyclic trajectory. FlowMap additionally produces a smoother projected velocity field that is more consistent with the reconstructed manifold. The fitted manifold provides a smooth low-dimensional representation of the data that captures the global structure of the cell-cycle trajectory (Fig. S12A), with embedding-derived angular coordinates showing strong agreement with ground-truth cell-cycle position (circular correlation = 0.60; Fig. S12B).

**Figure 2:**
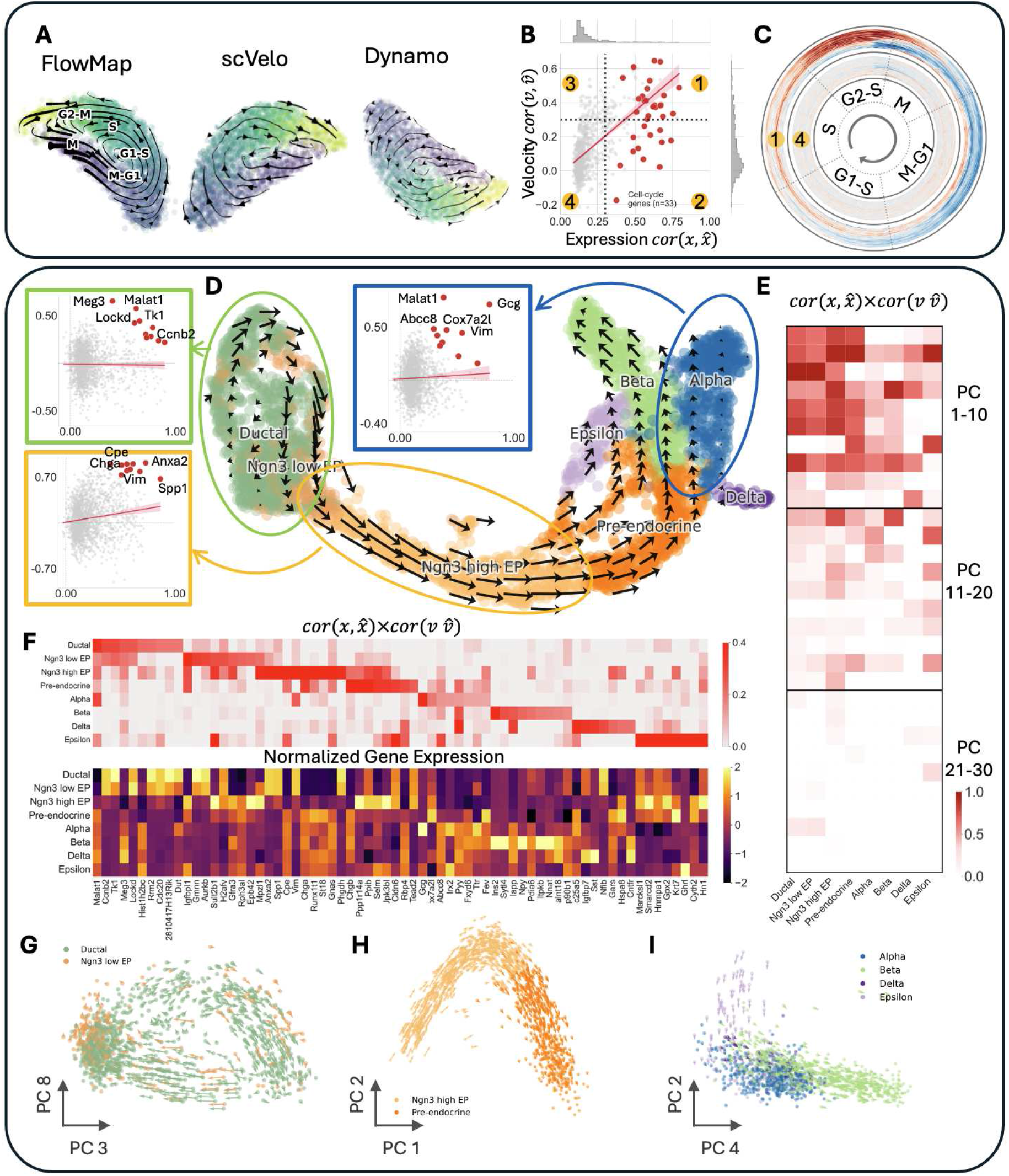
FlowMap embedding and reconstruction of gene expression and RNA velocity in cell-cycle and pancreatic datasets. **(A)** Velocity field embeddings of cell-cycle data using FlowMap, scVelo, and Dynamo. **(B)** Gene-level reconstruction consistency for expression and velocity. **(C)** Expression patterns of genes with high and low reconstruction consistency across cell-cycle phases. **(D)** FlowMap embedding of pancreatic development data with highlighted regions and representative genes. **(E)** Heatmap of reconstruction consistency across PC1–30 for each pancreatic cell type. **(F)** Reconstruction consistency and normalized expression of FlowMap selected top 10 consistency genes across pancreatic cell populations. **(G–I)** Velocity fields projected onto selected principal component pairs for ductal, endocrine progenitor, and endocrine subtype populations.

To quantify how well the reconstructed manifold preserves gene-level structure, we assessed consistency between observed and reconstructed values for both gene expression and RNA velocity. For the curated set of 33 known cell-cycle genes, the reconstruction achieves global *R*^2^ = 0.37 for expression and *R*^2^ = 0.24 for velocity when evaluated across cells (Fig. S12C; Methods, “Gene-level consistency of the reconstructed manifold and velocity field”). In contrast, across all 816 highly variable genes, reconstruction performance is substantially lower (*R*^2^ = 0.07 for expression and *R*^2^ = −0.10 for velocity), indicating that only a subset of genes exhibit dynamics consistent with the inferred manifold.

We further evaluated gene-level consistency by computing Pearson correlations between observed and reconstructed expression and velocity values (Fig. 2B), with the 33 curated genes highlighted. Based on these correlations, genes were stratified into consistent (expression *r >* 0.3 and velocity *r >* 0.3) and inconsistent groups (expression *r <* 0.3 and velocity *r <* 0.3). Consistent genes exhibit coherent, phase-aligned patterns along the cell-cycle trajectory, whereas inconsistent genes show no clear structure (Fig. 2C). These results indicate that FlowMap selectively captures genes whose dynamics are aligned with the underlying cellular trajectory.

### FlowMap identifies dynamical gene programs that preserve local velocity structure during pancreatic differentiation

To assess FlowMap in a heterogeneous developmental system without explicit temporal labels, we applied it to the pancreatic endocrinogenesis dataset [27]. In contrast to the relatively homogeneous cell-cycle trajectory, pancreatic differentiation contains multiple branching endocrine lineages and diverse cell states (Fig. 2D), suggesting that distinct transcriptional programs may govern local dynamics in different regions of the developmental manifold. The reconstructed velocity field is qualitatively similar to the original scVelo representation, while exhibiting a modest improvement in local smoothness after geometry-constrained projection (Fig. S9).

We first examined reconstruction consistency at the gene level and observed strong heterogeneity across the embedding (Fig. 2D,F). Different cell populations exhibited distinct subsets of genes whose expression and velocity were jointly reconstructed with high consistency, indicating that different transcriptional programs dominate in different developmental regions. In particular, ductal cells, endocrine progenitors, and mature endocrine subtypes each displayed locally enriched sets of dynamically consistent genes.

Importantly, the genes prioritized by FlowMap were substantially different from genes identified by conventional differential expression or RNA velocity kinetics analyses (Fig. 2F, Fig. S13). Differential expression methods prioritize genes that separate cell populations, while kinetic approaches identify genes with strong transcriptional dynamics. In contrast, FlowMap prioritizes genes whose low-dimensional representations preserve coherent local velocity structure (Fig. S14). Gene ontology analysis showed that these genes are enriched for diverse biological processes associated with lineage specification, differentiation, metabolism, and cellular organization across pancreatic cell populations (Fig. S17). As a result, FlowMap preserved nuanced local flow patterns that were weakened or obscured in embeddings constructed from conventional gene selections. These results suggest that embedding-consistent genes capture aspects of developmental dynamics that are not recovered by differential expression or kinetic significance alone.

To further characterize these dynamical programs, we next analyzed principal components (PCs) as proxies for coordinated transcriptional subspaces (Fig. S16). Reconstruction consistency varied substantially across PCs and cell states (Fig. 2E, S15), with different populations exhibiting high consistency in distinct PC subspaces. For example, ductal and Ngn3-low endocrine progenitor cells showed the strongest consistency in PC3 and PC8 (Fig. 2G, S18A), whereas later endocrine states were more coherent in PC1, PC2, and PC4 (Fig. 2H,I). These observations suggest that pancreatic differentiation is organized by multiple locally dominant dynamical programs rather than a single global mode of variation. Importantly, local vector fields can contain multiple partially competing or orthogonal dynamical programs (Fig. S18B), highlighting that any single low-dimensional embedding may capture only one dominant aspect of the underlying dynamics rather than fully representing the complete dynamical landscape.

### Interpreting gene expression gradients along developmental flows

Having established a geometry-consistent embedding of cellular states and velocities, we next used this representation to analyze how gene expression varies along inferred developmental flows. We applied this framework to the Larry dataset [28], which profiles 49,302 hematopoietic progenitor cells under-going continuous differentiation across a broad hematopoietic manifold (Fig. 3A). The smooth embedding reconstructed by FlowMap enables both cellular trajectories and gene expression to be represented directly on a continuous state space (Fig. 3B; Fig. S19).

**Figure 3:**
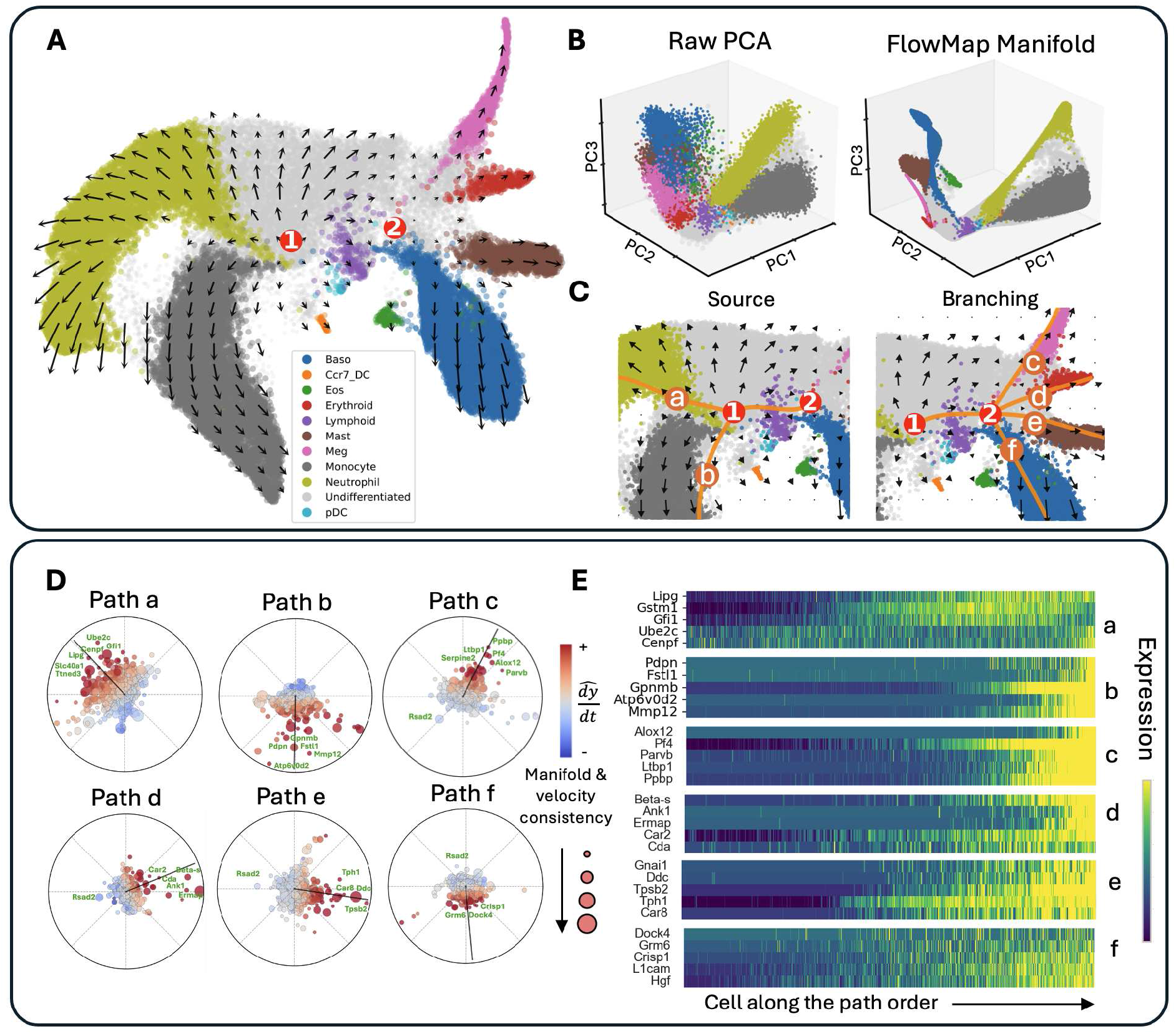
Gene-expression gradient alignment with RNA velocity. **(A)** FlowMap embedding with the reconstructed velocity vector field overlaid. **(B)** Comparison of the raw PCA embedding and the FlowMap-reconstructed manifold visualized in PC1–PC3 space. **(C)** Fixed points identified in the FlowMap embedding, illustrating the source and branching progenitor states together with least-action trajectories connecting progenitor and lineage-specific terminal states. **(D)** Gene markers identified by alignment between gene-expression gradients and the velocity vector field for each inferred path; the black arrow indicates the mean velocity direction, and points show the relative angle between each gene’s gradient and the velocity. **(E)** Expression of top-ranked genes along each path, ordered by cells along the inferred trajectory.

The geometric structure of the embedded velocity field is directly interpretable. We first identify fixed points as locations where the local velocity magnitude approaches zero (Fig. 3C). Combined with principal component consistency analysis at these locations (Fig. S20), we identify two major fixed points (Methods, “Vector field geometry”): a source corresponding to the undifferentiated root population (Fixed Point 1) and a branching progenitor state giving rise to megakaryocyte, erythroid, mast, and basophil lineages (Fixed Point 2). Starting from these fixed points, we infer developmental trajectories by computing optimal paths through the embedded velocity field using a least-action principle (Methods, “Least-action path formulation for trajectory inference”; Fig. 3C). This produces continuous developmental flows connecting progenitor and lineage-specific terminal states directly within the geometry of the embedding. Unlike pseudotime approaches, which impose a one-dimensional global ordering of cells, FlowMap defines differentiation trajectories locally through the vector field itself. As a result, trajectories naturally follow branch structure and local flow geometry, allowing distinct developmental programs to coexist within the same manifold without requiring a single shared progression axis.

Within this framework, gene expression is modeled as a smooth scalar function over the embedded state space (Fig. S19F). The gradient of this function specifies the local direction of maximal increase in expression, providing a geometric description of how transcriptional programs vary across cellular states (Methods, “Gene-expression gradients and velocity alignment”). In contrast to pseudotime-based analyses, which measure expression changes only along a predefined one-dimensional ordering, gradient fields are evaluated locally relative to the inferred developmental flow and can therefore capture branch-specific and path-dependent behavior.

When evaluated relative to the velocity field, the orientation of a gene’s gradient field characterizes its relationship to differentiation. Gradients may align with the developmental flow, oppose it, or lie largely orthogonal to it. Aligned gradients correspond to genes whose expression increases along differentiation trajectories, whereas anti-aligned gradients correspond to genes whose expression decreases during progression (Fig. 3D,E). Genes with primarily orthogonal gradients vary largely independently of the dominant differentiation axis, reflecting transcriptional variation that is locally structured but not strongly coupled to progression itself, and may distinguish neighboring subpopulations occupying similar positions along the same developmental trajectory (Fig. S21A,B).

Projecting gene expression gradients onto the inferred developmental flows further revealed distinct geometric patterns across trajectories. Genes with strongly negative projections exhibited substantial overlap across developmental paths, suggesting broadly shared transcriptional programs that decrease during differentiation (Fig. S21C). In contrast, genes with strongly positive projections showed substantially lower overlap between trajectories and were more frequently path-specific, consistent with lineage-associated programs that emerge preferentially along individual developmental branches (Fig. S21C). Representative examples are reflected directly in the expression patterns on the embedding (Fig. 3E; Fig. S21D), indicating that the geometry of the embedded flow captures both globally shared differentiation dynamics and branch-specific transcriptional programs.

### Curvature as a second-order descriptor of cellular state transitions

While RNA velocity provides a local estimate of the direction and magnitude of state change, it does not directly characterize how trajectories evolve along the cell-state space. In particular, cells undergoing similar instantaneous motion may nonetheless differ in whether their trajectories continue smoothly or undergo directional changes. To capture this aspect of dynamical structure, we analyzed the curvature of RNA velocity trajectories on the reconstructed FlowMap embedding. Curvature provides a second-order description of developmental flow, quantifying changes in trajectory direction rather than velocity magnitude alone.

This can be understood by analogy to motion constrained on a curved surface (Fig. 4A): even when the instantaneous velocity is fixed, trajectories may bend either because of the geometry of the underlying surface or because the direction of motion itself changes. In this setting, curvature captures how motion deviates from a straight trajectory, distinguishing between trajectories that passively follow the geometry of the state space and trajectories that actively redirect toward alternative developmental directions.

**Figure 4:**
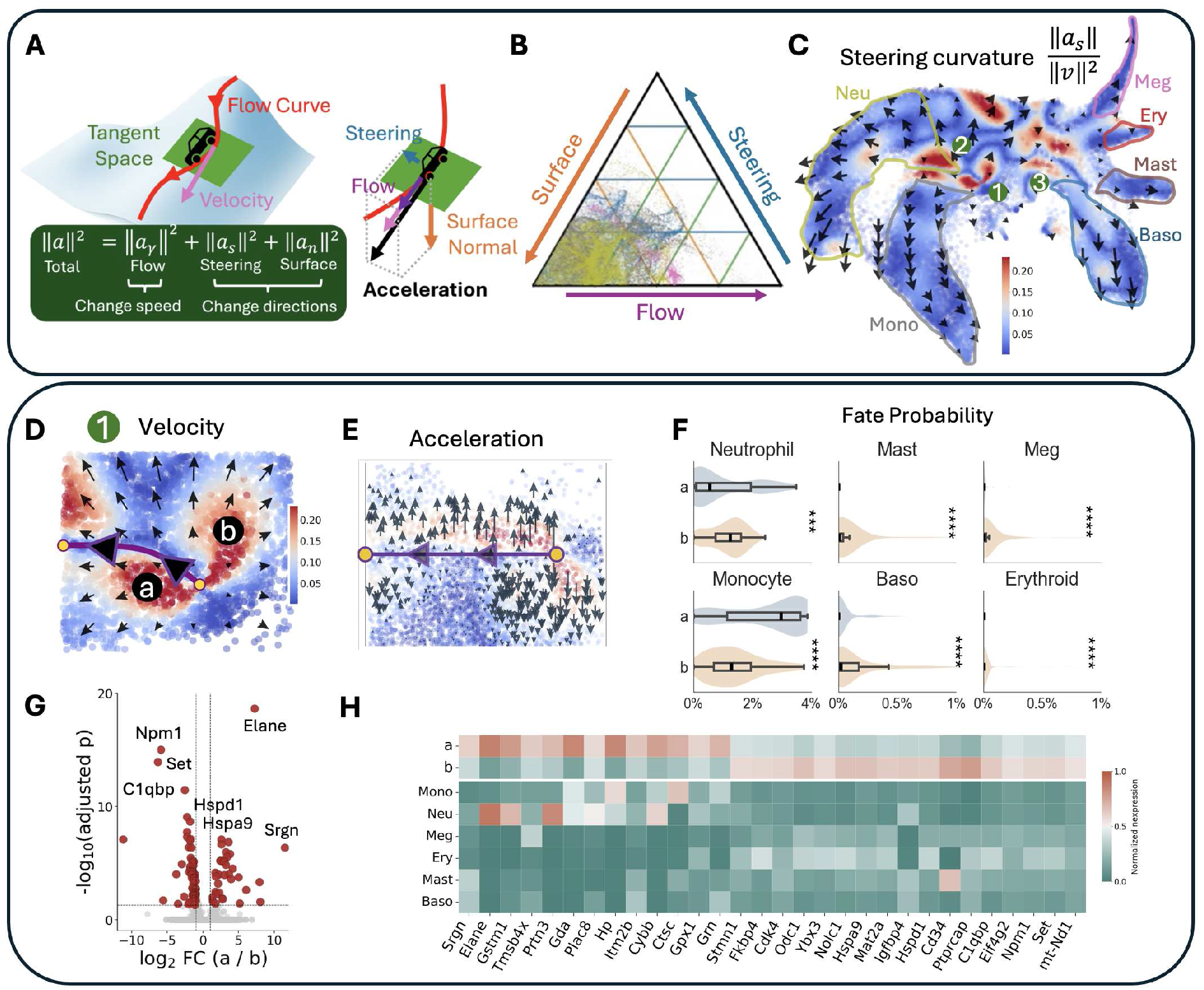
Curvature-based analysis of RNA velocity embeddings. **(A)** Schematic illustration of curvature decomposition along trajectories, showing flow, surface, and steering components of acceleration. Surface and steering components together define the curvature of the trajectory. **(B)** Ternary plot showing the distribution of curvature components across all cells. **(C)** FlowMap embedding colored by steering curvature, with three high-curvature regions highlighted. **(D)** Zoomed-in view of region 1, corresponding to an undifferentiated progenitor population exhibiting elevated steering curvature. Two subregions (a and b) with distinct directional tendencies are labeled. **(E)** Local velocity field and steering vectors in region 1, showing progenitor cells steering toward two distinct developmental directions. **(F)** Comparison of clonotype-derived and inferred fate probabilities between subregions a and b across annotated cell types. **(G)** Differential gene expression between subregions a and b, shown as a volcano plot of log_2_ fold change versus adjusted *p*-value. **(H)** Heatmap of top differentially expressed genes between subregions a and b across clusters and cell types.

Motivated by this intuition, several previous studies have explored curvature-related quantities to describe differentiation trajectories and developmental progression, often drawing connections to changes in cell fate or lineage commitment [5, 29, 30]. However, these approaches typically rely on scalar curvature measures or graph-based proxies that summarize trajectory bending as a single quantity and therefore do not distinguish between different sources of directional change. As a result, distinct dynamical behaviors—such as acceleration along a trajectory, bending imposed by the structure of the cell-state space, or active steering toward alternative states—are conflated.

FlowMap resolves this limitation by treating RNA velocity as a continuous description of cellular motion constrained to an embedded state space, allowing trajectory changes to be decomposed into three distinct and interpretable components (Fig. 4A), analogous to classical decompositions of motion on embedded manifolds in differential geometry [31]. First, the *flow* component captures acceleration along the trajectory itself, indicating whether cells are accelerating or decelerating during progression. Second, the *surface* component reflects bending imposed by the geometry of the embedded state space, capturing how trajectories follow the global organization of accessible cell states. Third, the *steering* component captures directional changes that occur within the manifold of valid states, highlighting local redirection toward alternative trajectories or developmental branches. Together, the surface and steering components define the curvature of the developmental flow (Fig. S22).

Across the full cell population, the surface curvature component accounts for the largest fraction of the total curvature signal (Fig. 4B), indicating that much of the observed trajectory bending reflects the geometry of the reconstructed cell-state manifold. As a result, scalar curvature measures alone may obscure more localized directional changes in developmental flow. To isolate these directional effects, we examined steering curvature across the cell-state manifold. Steering curvature was highly localized within gene expression (Fig. 4C), with elevated values concentrated in early progenitor populations and substantially weaker signals in terminally differentiated states. We identified three major steering-curvature regions (Fig. 4C; Fig. S23), all associated with transitional progenitor states.

Region 1 corresponds to an undifferentiated progenitor population located upstream of the neutrophil/monocyte (Neu-Mono) and megakaryocyte/mast cell/erythroid/basophil (Meg-Mast-Ery-Baso) lineages. Within this region, steering curvature reveals two subpopulations exhibiting distinct developmental directions (Fig. 4D,E), despite remaining closely intermixed in the low-dimensional embedding. These subpopulations show differences in both clonotype-derived and inferred fate probabilities (Fig. 4F), with one branch preferentially biased toward neutrophil and monocyte differentiation and the other toward megakaryocytic, erythroid, mast-cell, and basophil fates. Differential expression analysis identified transcriptional programs that distinguish these progenitor states (Fig. 4G,H). The Neu-Mono-biased branch showed elevated expression of genes associated with early myeloid differentiation, including *Elane, Prtn3*, and *Gstm1*, whereas the Meg-Mast-Ery-Baso-biased branch exhibited increased expression of lineage-associated genes such as *Pf4, Hba-a2*, and *Car1*. These results suggest that steering curvature captures lineage-specific molecular biases before the emergence of clearly separated terminal populations. Notably, the transcriptional differences observed within the progenitor compartment were more pronounced than those observed between downstream mature states, indicating that early developmental commitment may be encoded through subtle directional changes in cellular trajectories.

Similar patterns were observed in Regions 2 and 3 (Fig. S23), where localized steering curvature identified additional progenitor populations undergoing lineage diversification. In each case, steering curvature highlighted transitional states associated with distinct fate preferences and transcriptional programs, supporting a broader role for directional geometry in revealing early developmental decision points that are not readily apparent from cell-state position alone.

### Curvature analysis reveals pausing-associated transition states in B-cell differentiation

FlowMap embeddings additionally enable differential-geometric analysis of the reconstructed gene-expression landscape. Steering curvature and Ricci curvature characterize complementary aspects of this landscape. Steering curvature measures how RNA-velocity vectors change direction along cellular trajectories, highlighting local turning points and bifurcations. In contrast, Ricci curvature describes the geometry of the gene-expression manifold itself, independent of the inferred velocity vectors. Intuitively, Ricci curvature measures whether nearby geodesics on the manifold tend to converge (positive curvature) or diverge (negative curvature), thereby providing a geometric characterization of how the underlying state space is organized (Fig. 5A). Ricci curvature is closely related to the surface component of the curvature of velocity-flow trajectories, as both reflect how the induced manifold geometry shapes local flow structure (Fig. 4A; Supplementary Note 3.2), and these quantities exhibit similar spatial patterns in the B-cell embedding (Fig. S24A). Compared with surface curvature, Ricci curvature provides a more intrinsic description of the geometry because it depends only on the reconstructed manifold rather than the local velocity vectors, making it useful for characterizing the organization of cellular state space. Recent studies have likewise shown that curvature-based geometric analyses can reveal developmental organization and trajectory rewiring in biological systems [32].

**Figure 5:**
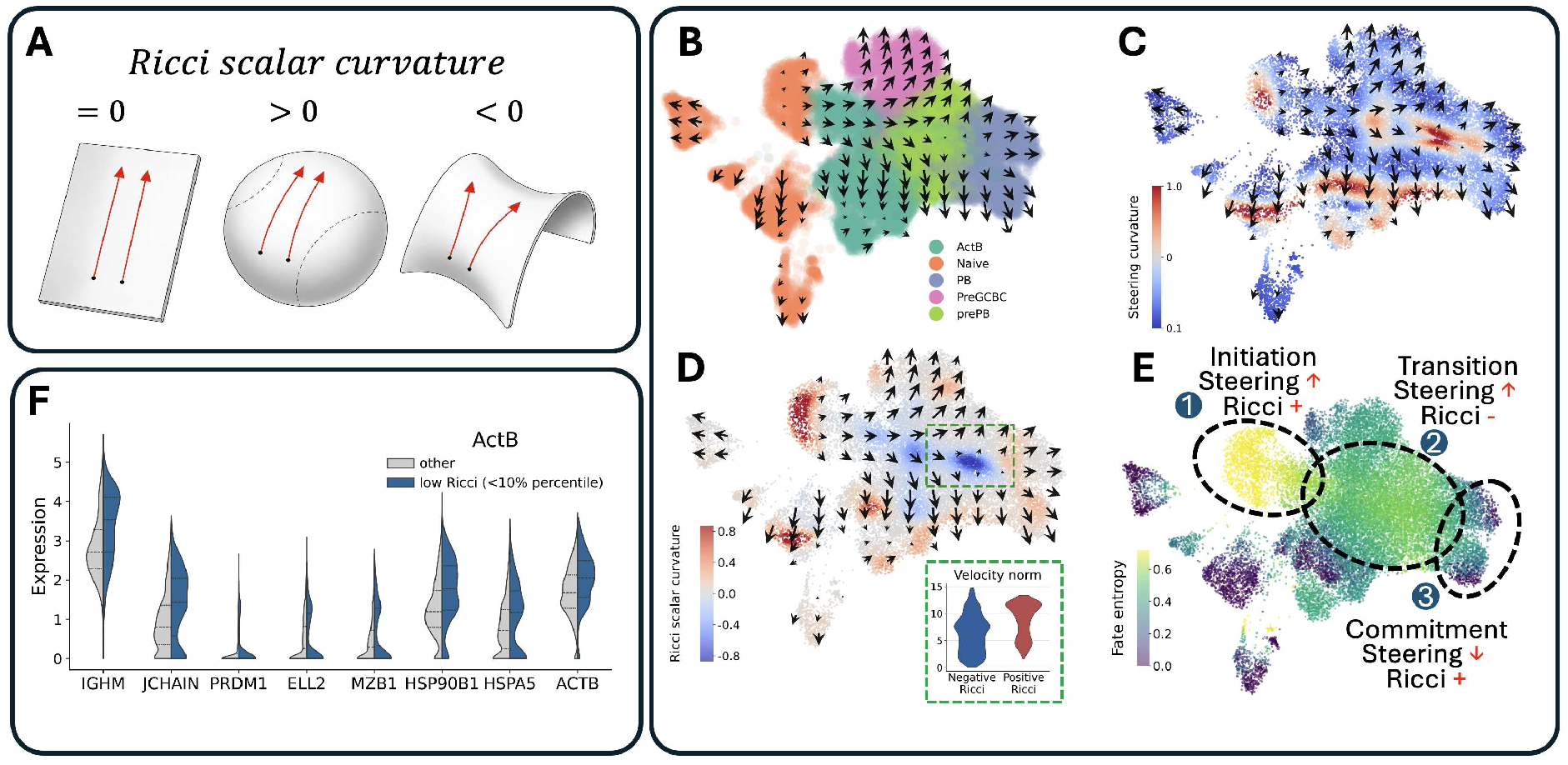
Differential-geometric characterization of B-cell differentiation dynamics using FlowMap. **(A)** Schematic illustration of Ricci scalar curvature on a manifold. Positive curvature corresponds to converging geodesics, zero curvature to parallel geodesics, and negative curvature to diverging geodesics. **(B)** FlowMap embedding of B-cell differentiation trajectories with the inferred RNA velocity vector field overlaid. **(C)** Embedding colored by steering curvature. **(D)** Ricci scalar curvature computed from the induced embedding geometry, with the highlighted transition region used to compare velocity norm between low- and higher-Ricci cells. **(E)** CellRank fate entropy projected onto the embedding. Higher entropy reflects increased uncertainty in downstream lineage commitment. **(F)** Expression distributions of representative genes comparing activated B cells (ActB) with low Ricci curvature, defined as the bottom 10th percentile of Ricci scalar curvature, against the remaining ActB population.

To demonstrate this capability, we applied FlowMap to a human B-cell differentiation dataset in which RNA velocity reveals bifurcating trajectories from activated B cells toward plasmablast (PB) and germinal center precursor (preGC) fates [33] (Fig. 5B). Steering curvature identifies regions where developmental trajectories undergo pronounced directional changes and branching events, providing a geometric measure of local trajectory dynamics (Fig. 5C). In contrast, Ricci scalar curvature characterizes the intrinsic geometry of the reconstructed gene-expression landscape, independent of the local velocity direction (Fig. 5D). Positive Ricci curvature was observed in the initial activated B-cell population and terminal differentiated states, whereas negative Ricci curvature emerged in intermediate transition regions connecting alternative developmental fates. Notably, these negative-curvature regions closely overlapped with previously reported low-velocity pausing states during fate specification, suggesting that developmental pausing occurs in geometrically divergent regions of the underlying state space (Fig. 5D).

We next compared Ricci curvature with fate entropy, which quantifies the uncertainty of a cell’s future developmental fate (Fig. 5E). Together, steering curvature, Ricci curvature, and fate entropy capture complementary geometric aspects of differentiation. Steering curvature describes how developmental trajectories locally change direction, Ricci curvature characterizes whether the underlying landscape is locally convergent or divergent, and fate entropy indicates a cell’s position along the progression from multipotency toward terminal commitment. Despite arising from these distinct principles, all three measures consistently identified the intermediate transition region separating the initial and terminal cell states (Fig. 5E; Fig. S24B). This concordance suggests that the geometry of the reconstructed gene-expression landscape captures biologically meaningful features of developmental organization beyond trajectory direction alone. Differential expression analysis further identified genes enriched within negative-curvature regions, including plasmablast-associated programs such as JCHAIN, PRDM1, and ELL2 (Fig. 5F), indicating that these geometrically divergent transition regions are accompanied by transcriptional programs associated with emerging fate commitment.

### Identifying spatially localized gene effects during mouse organogenesis

Spatial gradients of gene expression play a central role in morphogenesis, where coordinated changes in gene activity establish tissue patterning, developmental boundaries, and spatial axes through morphogen-mediated signaling programs [34]. Recent work has shown that gene expression gradients can be treated as geometric objects over spatial transcriptomic coordinates [35]. Here, we extend this perspective by analyzing spatial gradients relative to transcriptional dynamics inferred from RNA velocity.

We analyzed a spatial mouse organogenesis dataset [36] by first estimating RNA velocity with SIRV [37] and embedding the resulting cellular states and dynamics using FlowMap. The inferred RNA velocity vectors represent local derivatives of the transcriptomic state, visualized at the spatial coordinates of the corresponding cells. This representation relates transcriptional dynamics to tissue organization while preserving the anatomical context of the tissue (Fig. 6A). On the resulting developmental landscape, we estimated for each gene the direction of increasing expression, thereby defining a spatial gradient field over the tissue embedding.

**Figure 6:**
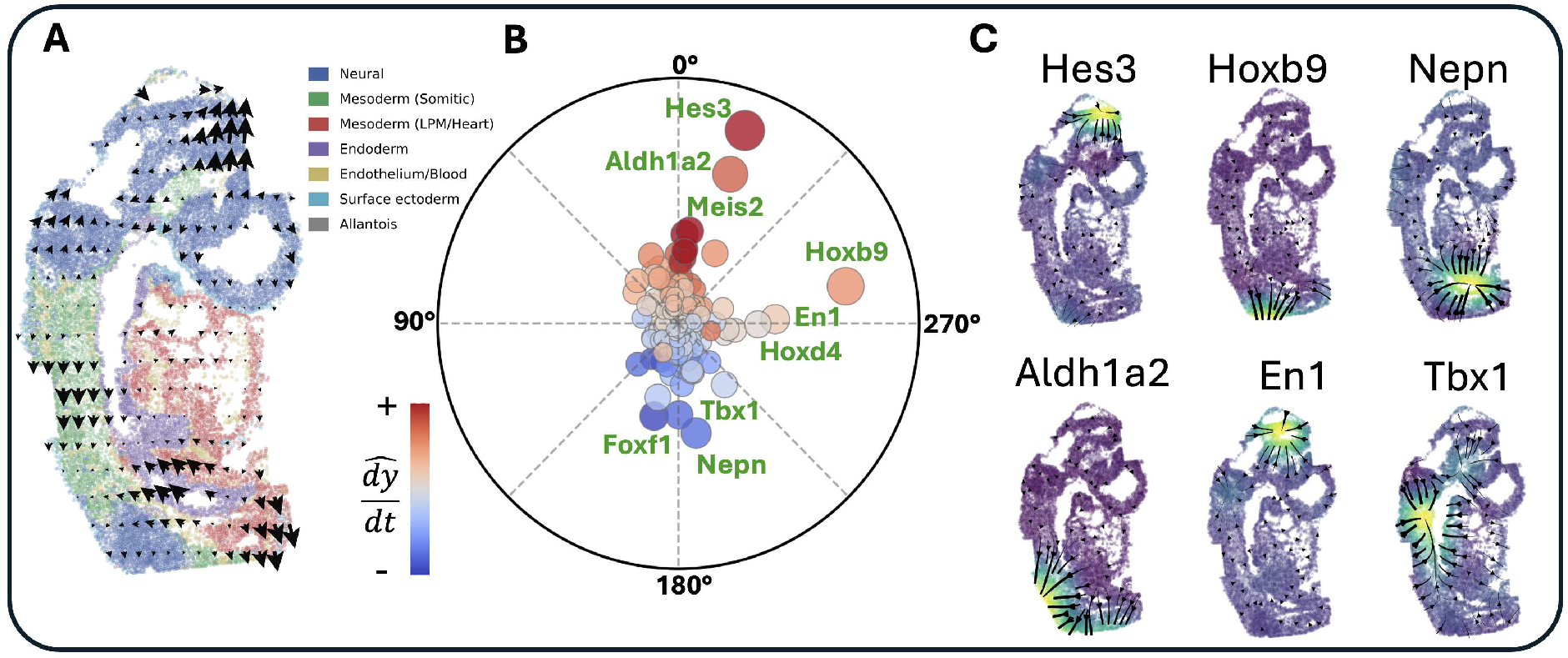
Gene gradient geometry and RNA velocity in spatial mouse organogenesis. **(A)** Spatial RNA velocity field overlaid on tissue spatial coordinates. **(B)** Polar plot of gene expression gradients: angle denotes gradient direction relative to the velocity, radius denotes magnitude, and color indicates alignment with velocity. **(C)** Spatial gradient fields for representative genes, showing expression patterns and local gradient directions.

Comparing gene expression gradients with the local velocity field allows us to characterize how spatial patterning relates to developmental progression. When gradients align with the inferred flow, gene expression increases along the direction of transcriptional change (Fig. 6B; Fig. S25). Consistent with known developmental biology, several aligned genes correspond to active signaling and patterning programs (Fig. 6C). For example, *Aldh1a2*, a key enzyme in retinoic acid synthesis involved in anterior–posterior patterning during embryogenesis [38], exhibits strong alignment across neural and mesodermal tissues, suggesting increasing activity along the inferred developmental trajectory.

In contrast, genes with gradients largely orthogonal to the velocity field appear less coupled to the dominant direction of developmental progression and instead reflect positional or regional patterning cues (Fig. 6C). For instance, *Hoxb9*, a posterior Hox gene associated with axial identity specification, exhibits a gradient largely decoupled from the local transcriptional flow. Similar relationships were observed for *Hes3* and *En1* in neural regions, and *Nepn* and *Tbx1* in endodermal and mesodermal tissues, highlighting diverse interactions between spatial organization and transcriptional dynamics across developmental lineages.

## Discussion

We introduce a geometric framework for RNA velocity that explicitly couples the representation of cellular states with their associated dynamics. Rather than treating low-dimensional embeddings and velocity estimates as separate objects, FlowMap models a smooth manifold together with its local tangent structure, ensuring that inferred dynamics are consistent with the geometry of the state space. This provides a principled foundation for interpreting RNA velocity as a vector field over cellular states.

Existing RNA velocity pipelines typically construct embeddings based on gene expression and incorporate directional information post hoc through smoothing or graph-based methods. While effective for visualization, these approaches do not enforce geometric consistency between cellular states and their inferred dynamics. FlowMap addresses this gap by reconstructing a differentiable mapping between embedding and gene-expression space, enabling velocity to be interpreted within a well-defined geometric framework.

Our results show that cellular dynamics in heterogeneous systems are not governed by a single global program, but instead arise from multiple locally structured dynamical processes. These local dynamics correspond to distinct biological programs that dominate in different regions of the state space and can be resolved through subspace-specific representations. Within this framework, gene-level gradients provide a natural way to identify trajectory-specific markers aligned with local directions of change, linking dynamical structure to underlying regulatory programs. In addition, analysis of steering curvature highlights regions where trajectories diverge, enabling the identification of putative fate decision points. Together, these results demonstrate that modeling RNA velocity within a geometric framework enables both a more faithful representation of dynamics and richer biological interpretation.

FlowMap relies on the quality of the input RNA velocity estimates, which can vary depending on modeling assumptions and data characteristics. The geometric framework provides a way to assess these estimates: discrepancies between predicted and reconstructed dynamics can be used to evaluate and compare velocity methods, offering a principled diagnostic for model selection. In addition, as with any low-dimensional representation, the embedding inevitably compresses high-dimensional gene-expression data and cannot capture all aspects of the underlying dynamics. Nevertheless, the spline-based parametric reconstruction enables the identification of dominant dynamical modes and their local structure, allowing key patterns of cellular progression to be recovered despite this compression. Cell density also varies across the state space, and some states may therefore be represented by few or no observed cells. The continuous spline representation can interpolate across sparsely sampled regions and, in principle, extrapolate into nearby unobserved regions. Such estimates are expected to be most reliable when supported by nearby observed cells.

More broadly, FlowMap’s input-based design is compatible in principle with velocity fields inferred from spliced and unspliced RNA [3, 4], derived from time-resolved RNA measurements using metabolic labeling [39], or estimated from multimodal data using methods such as MultiVelo [40]. In particular, MultiVelo extends RNA velocity estimation by incorporating chromatin accessibility into a dynamical model of gene expression; from the perspective of FlowMap, its output is an alternative upstream velocity estimate that can be analyzed under the same geometric constraints. This compatibility offers a potential route toward interpreting velocity estimates from different upstream methods and measurement modalities within a shared geometry-constrained framework.

In heterogeneous systems, these dynamics are inherently multi-modal, and a single embedding may capture only one dominant mode at a time. A more complete understanding would require decomposing the system into multiple coupled dynamical components, which could be further resolved within the same geometric framework. Recent approaches have suggested that such heterogeneity may arise from evolving gene regulatory programs that can be jointly inferred together with RNA velocity dynamics [41–43]. Within this view, changes in gene regulatory structure may govern the large-scale organization of the underlying dynamical landscape, while local geometric representations capture its observable trajectories. More broadly, this perspective suggests that cellular dynamics are best understood as compositions of local geometric processes rather than a single global flow.

## Methods

### Interpretation of RNA Velocity

RNA velocity provides a quantitative description of how a cell’s molecular profile changes over time, inferred from the coupling between unspliced (*U*) and spliced (*S*) mRNA abundances. The kinetic model for a single gene is:

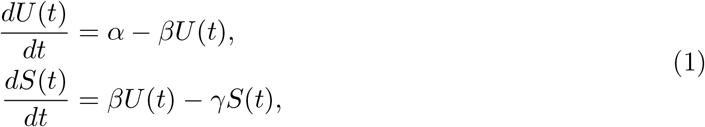

where *α, β*, and *γ* are gene-specific kinetic parameters.

For a given cell, the RNA velocity vector

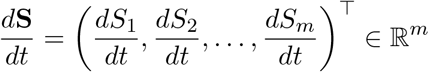

represents the instantaneous change in expression across all *p* genes.

#### Manifold view and role of the Jacobian

Intuitively, a cell’s state can be described by a small set of intrinsic coordinates y ∈ ℝ^*d*^ that capture its biological identity and position along developmental or functional trajectories. Even though gene expression lives in a high-dimensional space ℝ^*m*^, not every combination of expression levels is biologically valid — cells occupy only a smooth, low-dimensional surface (manifold) within this space. The mapping *ψ*: ℝ^*d*^ → ℝ^*p*^ takes a point in this intrinsic *state space* to its corresponding gene expression profile.

The Jacobian J_*ψ*_(y) tells us how small changes in state coordinates translate into changes in gene expression. It defines the local *tangent space* — the set of all gene expression changes that are consistent with a valid movement along the manifold. When we take an observed RNA velocity vector 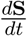 and project it onto this tangent space, we filter out components that would move the cell off the manifold (i.e., biologically implausible directions) and keep only those that reflect genuine state changes. This projection step both denoises the velocity vector and constrains it to represent an evolution of the cell’s underlying state.

#### Chain rule connection

By the multivariate chain rule, RNA velocity (change in expression per unit time) factors into:

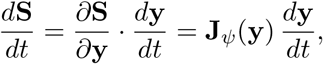

where J_*ψ*_(y) ∈ ℝ^*p*×*d*^ is the Jacobian of *ψ*, with entries *∂S*_*i*_*/∂y*_*j*_.

Expanded in matrix form:

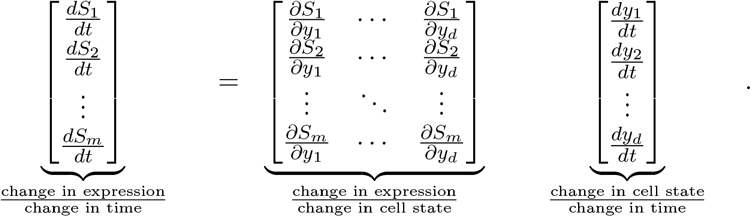

#### Recovering state-space velocity

Conceptually, we want to answer the question: *what change in cell state would best explain the observed change in gene expression?* The Jacobian J_*ψ*_(y) acts as a bridge between the two spaces: it tells us how a small movement in state space translates into changes in gene expression. By using this bridge in reverse, we can map the observed gene expression change back into a change in state coordinates, giving us the intrinsic velocity 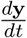 that best explains the data.

Formally, we solve:

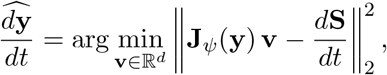

whose closed-form solution uses the Moore–Penrose pseudoinverse:

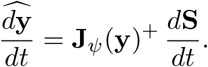

In a 2-D embedding, 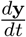 gives the best local linear approximation of how the cell’s state is evolving, yielding a denoised and interpretable trajectory in the intrinsic state space.

### FlowMap embedding method

#### Overview of the FlowMap framework

The core rationale behind FlowMap is to model the gene expression space as a *differentiable manifold*, where RNA velocity defines a continuous vector field over this manifold. By learning a smooth, parametric mapping between the low-dimensional embedding and the high-dimensional expression space, FlowMap preserves the manifold’s geometric structure and enables direct, quantitative analysis of cellular dynamics.

Common embedding methods such as t-SNE, UMAP, or PHATE are powerful for preserving local or global geometry, yet they do not retain an explicit mapping to the original space. Using these embeddings as a starting point, FlowMap reconstructs them as a differentiable manifold, providing a major advantage: the ability to map between embedding and gene space, recover tangent spaces, and quantify feature contributions in a consistent framework. This differentiable structure expands the interpretability of the embedding beyond visualization while preserving the strengths of the original method, including:

- **Parametric mapping:** reconstructing the manifold explicitly via a spline function, yielding a smooth, analytical representation of the data and enabling direct mapping between embedding coordinates and gene expression space.
- **Interpretability:** providing a transparent alternative to graph-based or heuristic embeddings, with well-defined derivatives that support velocity projection and gene-level attribution.
- **Geometric understanding:** enabling the analysis of local and global structure, including curvature, fixed points, and branching topology.

Guided by these goals, FlowMap introduces several key components:

- Incorporating directional information through a velocity-aware distance metric that captures both spatial proximity and local flow.
- Reconstructing the manifold with a spline mapping, enabling smooth back-projection and tangent space computation.
- Refining embeddings via joint optimization of positions and vector alignment to minimize reconstruction error and velocity mismatch.
- Analyzing manifold geometry to classify fixed-point dynamics (e.g., sources, sinks, saddles) and identify genes aligned with inferred trajectories.

FlowMap takes as input a gene expression matrix *X* ∈ ℝ^*N*×*p*^ and a corresponding RNA velocity matrix *V* ∈ ℝ^*N*×*p*^ for *N* cells and *p* genes. The goal is to construct a low-dimensional embedding that is geometrically consistent with the RNA velocity–defined dynamics.

The procedure consists of the following steps:

1. **Velocity-aware distance construction**. For each pair of cells, we compute a phase distance that quantifies their relative position along the RNA velocity flow (“ Velocity-aware distance metric“). This phase distance is combined with Euclidean distance in expression space using a weighting parameter *α*, yielding a velocity-aware distance matrix.
2. **Low-dimensional embedding**. A neighborhood graph is constructed from the combined distance matrix, and a low-dimensional embedding is obtained using a standard manifold learning algorithm such as UMAP or t-SNE. This produces an initial representation that reflects both expression similarity and directional progression.
3. **Smooth manifold reconstruction**. To obtain an explicit and differentiable mapping between the embedding and gene expression space, we fit a spline that maps embedding coordinates to the original expression space (or a reduced PCA representation; “ Manifold reconstruction“). This reconstruction defines a smooth parametric manifold underlying the embedding.
4. **Tangent-space velocity projection**. Using the spline mapping, we compute the tangent space of the reconstructed manifold at each embedded point (“ Tangent space and velocity projection“). RNA velocity vectors are projected onto these tangent spaces, ensuring that inferred transcriptional changes are consistent with the geometry of the embedded state space.
5. **Optional joint refinement**. FlowMap optionally supports a joint optimization procedure that adjusts embedding coordinates and projected velocities by minimizing expression reconstruction error and velocity mismatch (“Embedding optimization algorithm“). This step is not required for geometric consistency and is disabled by default.

Together, these steps produce a low-dimensional representation in which cellular states and RNA velocity are embedded in a geometrically coherent manner, with velocity vectors constrained to lie along the reconstructed manifold.

### Velocity-aware distance metric

In the context of a continuous vector field, the *phase* of a point can be understood as its position along a flow trajectory, analogous to a timestamp measured in the intrinsic coordinates of the field. Points with the same phase lie on level sets orthogonal to the direction of flow, and moving along the flow direction changes phase while preserving other coordinates on the manifold. For RNA velocity data, phase provides a natural way to quantify temporal progression between cells, separating differences aligned with the primary direction of flow from those arising in directions orthogonal to it.

To compute the phase distance between two nearby cells, FlowMap estimates their relative position along the local velocity flow by allowing each cell to move along its RNA velocity vector and measuring how much progression is required for the two states to align. Intuitively, if two cells lie at different phases along the same trajectory, they can be brought closer in expression space by sliding them forward or backward along the flow; the amount of sliding reflects their separation along the dynamical direction. Cells that are synchronized along the flow have similar optimal offsets and thus small phase distance, whereas cells that are ahead of or behind one another along a trajectory require different displacements to align and therefore have large phase distance. A formal definition of the resulting phase distance, along with its interpretation as a symmetric first-order time difference, is provided in Supplementary Notes 1.1–1.2.

Formally, let *x*_*i*_, *x*_*j*_ ∈ ℝ^*p*^ denote the gene expression profiles of cells *i* and *j*, with corresponding RNA velocity vectors *v*_*i*_, *v*_*j*_ ∈ ℝ^*p*^. We define their optimal alignment by solving

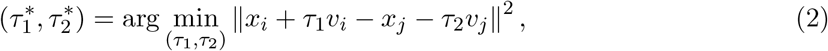

which finds the pair of displacements along the velocity directions that minimizes the distance between the two states after perturbation. The scalars *τ*_1_ and *τ*_2_ can be interpreted as local progression times along the flow required to best align the two cells.

This optimization admits a closed-form solution obtained by solving a 2 × 2 linear system for each pair of cells. Details of the derivation, numerical implementation, and stability considerations are provided in Supplementary Note 1.3.

To capture both spatial proximity and this notion of dynamical phase, FlowMap computes the *pairwise phase distance* between cells using both their gene expression profiles and local velocity vectors (Supplementary Note 1). This phase distance is then combined with the standard Euclidean distance to define a *velocity-aware distance*:

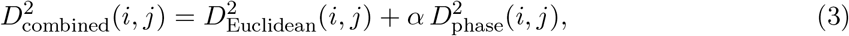

where *D*_Euclidean_(*i, j*) is the standard Euclidean distance between cells *i* and *j* in expression space, *D*_phase_(*i, j*) is their estimated phase distance, and *α* controls the relative weighting between spatial and dynamical separation.

The resulting metric is small for cells that are both close in expression space and synchronized in their progression along the flow, and large for cells that are separated along the trajectory. By preserving this combined distance in the embedding, FlowMap aligns the low-dimensional geometry with the underlying cellular dynamics.

Starting from the velocity-aware distance matrix, FlowMap constructs a neighborhood graph by connecting each cell to its *k* nearest neighbors (KNN) based on the combined metric. This graph serves as the input to manifold learning algorithms such as UMAP or t-SNE, enabling embeddings that jointly preserve spatial similarity and temporal progression derived from the vector field.

### Manifold reconstruction

A key limitation of common embedding methods such as t-SNE or UMAP is the absence of an explicit, differentiable mapping between the low-dimensional embedding and the original high-dimensional gene expression space. These methods typically rely on cell–cell transition probabilities or neighborhood graphs, which are effective for visualization but limit interpretability: without a mapping function, it is not possible to directly recover local tangent spaces, quantify feature contributions, or project inferred trajectories back into gene space.

FlowMap addresses this limitation by reconstructing the embedding as a smooth *differentiable manifold* using a spline-based function. This approach preserves the geometric structure of the original embedding while acting as an effective denoising step: by enforcing smoothness, the spline filters out high-frequency measurement noise and retains only the dominant, coherent patterns of variation that drive the cellular dynamics. In this way, the learned manifold provides a low-dimensional approximation of the high-dimensional data that captures the main dynamical trends, enabling both interpretable geometry and reduced noise sensitivity. By endowing the embedding with an explicit, parametric mapping, the reconstructed surface supports rigorous geometric analysis, enabling downstream computations such as curvature, geodesics, and local basis vectors.

Formally, let X ∈ ℝ^*N*×*p*^ denote the matrix of gene expression profiles for *N* cells and *p* genes, and Y ∈ ℝ^*N*×*d*^ their corresponding low-dimensional coordinates (*d* ≪ *p*, typically *d* = 2). We seek a smooth mapping *ψ*: ℝ^*d*^ → ℝ^*p*^ such that

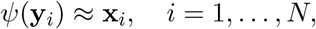

where x_*i*_ and y_*i*_ are the expression profile and embedding coordinates of cell *i*, respectively. We model *ψ* as a vector-valued spline in a reproducing kernel Hilbert space (RKHS) [44],

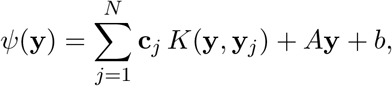

where *K*(·, ·) is a positive-definite kernel, {c_*j*_} are coefficient vectors, and *A* ∈ ℝ^*p*×*d*^, *b* ∈ ℝ^*p*^ define the affine component.

The coefficients are obtained by solving the regularized regression problem

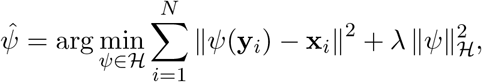

where *H* denotes the RKHS associated with *K*, and *λ >* 0 controls the smoothness–accuracy trade-off. Details of the linear system solution and numerical implementation are provided in Supplementary Note 2.1-2.2.

A common choice for *K* is the class of *polyharmonic spline* kernels [22, 45], defined as radial basis functions of the form

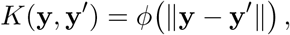

where

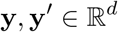

are embedding coordinates and *d* denotes the embedding dimension. The radial function

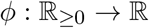

is given by

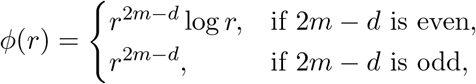

where *m* denotes the spline order and satisfies

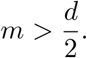

Higher values of *m* produce smoother interpolating functions by penalizing higher-order derivatives of the reconstructed manifold.

#### Thin-plate spline as a special case

When *d* = 2 and the kernel is chosen as the biharmonic radial basis function

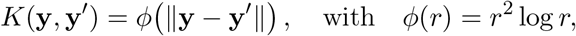

the above formulation reduces to the classical thin-plate spline (TPS), where the RKHS norm corresponds to a bending energy penalty.

#### Smoothing parameter selection

The parameter *λ* controls the effective complexity of the reconstructed surface by trading off fidelity to the data and smoothness of the mapping. Smaller values of *λ* allow the spline to closely interpolate the observed data, while larger values enforce a smoother, lower-variance surface. In practice, *λ* is selected using generalized cross-validation (GCV), which provides a data-driven estimate of the optimal bias–variance trade-off (Supplementary Note 2.3).

### Tangent space and velocity projection

Given the smooth mapping

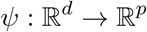

from embedding to gene expression space, the *tangent space* at an embedded point y is the *d*-dimensional subspace of ℝ^*p*^ spanned by the columns of the Jacobian

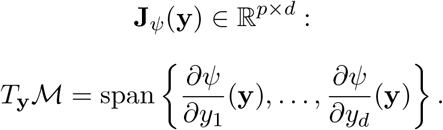

The Jacobian J_*ψ*_(y) provides a local linear approximation of the manifold, mapping embedding coordinates to gene expression space. A closed-form expression for J_*ψ*_(y) under the spline model is given in Supplementary Note 2.4.

Given a high-dimensional velocity vector

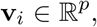

we estimate its low-dimensional representation by projecting it onto the local tangent space. This is formulated as the least-squares problem

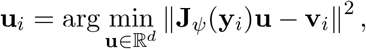

which seeks the embedding-space vector whose pushforward best matches the observed velocity.

The solution is given by

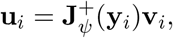

where 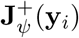 denotes the Moore–Penrose pseudoinverse. This yields a low-dimensional velocity field

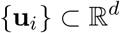

defined on the embedding.

To obtain a smooth and interpretable vector field, we fit a **second spline function**

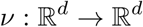

to the projected velocities {**u**_*i*_}. Unlike the manifold reconstruction *ψ*, which maps embedding coordinates to gene expression space, *ν* operates entirely within the embedding and models the velocity field directly in low-dimensional coordinates. The resulting function defines a continuous vector field in the embedding space that is consistent with the geometry induced by the reconstructed manifold.

### Embedding optimization algorithm

The tangent space formulation naturally suggests that the embedding can be refined to better align with the underlying dynamics. While standard embedding methods (e.g., UMAP, t-SNE) are constructed solely from positional relationships, they do not explicitly enforce consistency with RNA velocity. As a result, the embedding may distort dynamical structure even when local neighborhoods are preserved.

FlowMap addresses this by refining an initial embedding using both positional and velocity information. Let *ψ* denote the spline mapping from embedding coordinates to gene expression space, and *ν* denote the low-dimensional velocity field. We optimize the embedding coordinates {*y*_*i*_} by minimizing the joint objective

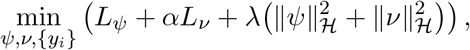

where

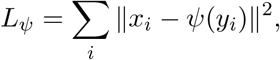

and

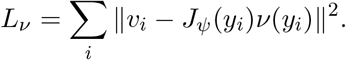

Here, *L*_*ψ*_ enforces reconstruction of the observed gene expression profiles, while *L*_*ν*_ enforces consistency between the observed high-dimensional velocities *v*_*i*_ and their projection through the Jacobian *J*_*ψ*_(*y*_*i*_). The smoothness terms

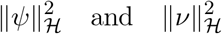

are RKHS norms associated with the spline model, controlling the regularity of both the manifold and the velocity field.

By jointly minimizing these terms, the embedding is iteratively adjusted to reduce mismatches between observed expression and inferred dynamics, yielding a representation that is both geometrically faithful and dynamically consistent.

#### Algorithm 1

Iterative refinement of embedding and velocity field

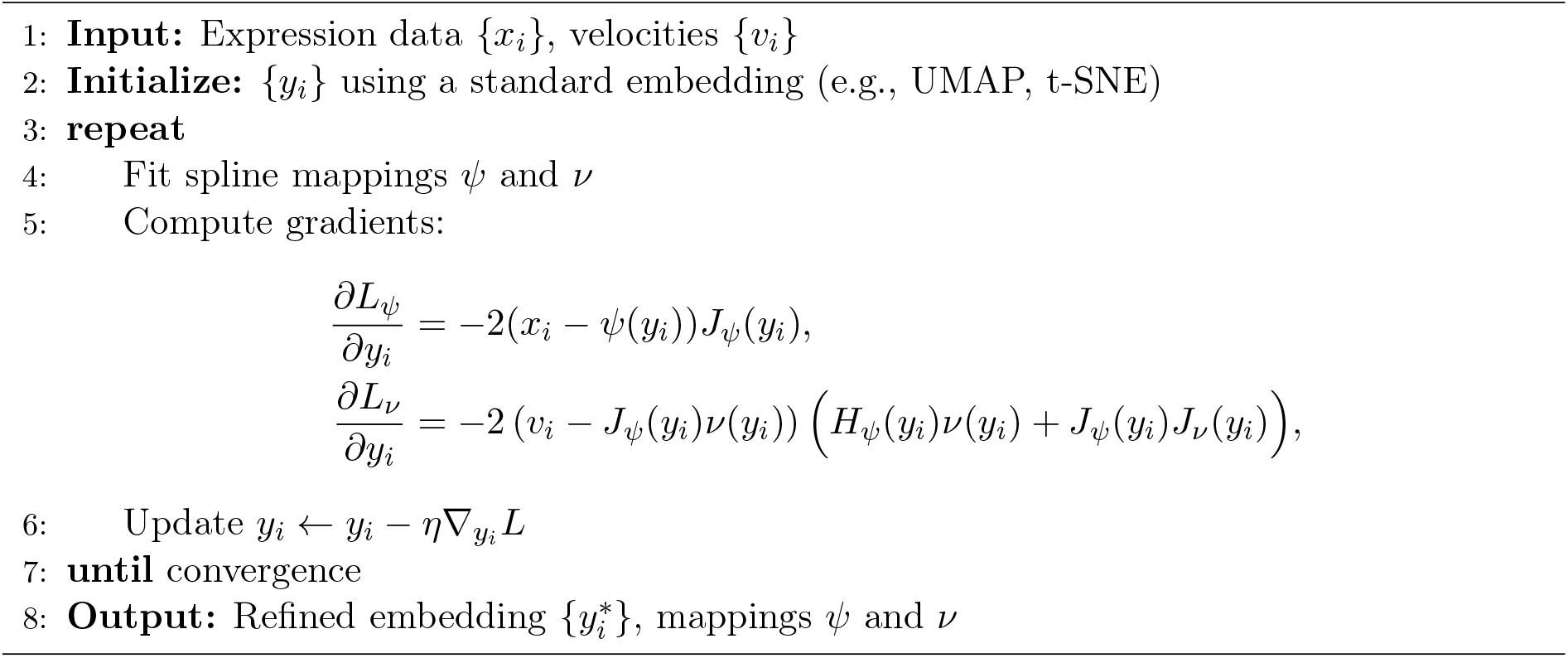

#### Algorithm 1: Iterative Embedding Refinement

This procedure refines the initial embedding by repositioning cells in regions where velocity information is inconsistent with the current geometry, improving alignment between cellular states and their inferred dynamics.

### Interpreting differentiable velocity fields with FlowMap

#### Gene-level consistency of the reconstructed manifold and velocity field

To assess how well the reconstructed manifold captures the underlying cellular dynamics, we evaluate consistency between observed and reconstructed quantities for both expression and RNA velocity. Although this analysis is typically performed at the gene level, the same framework can also be applied to principal components or other feature representations.

Let *i* index cells and *j* index features (e.g., genes or principal components). Given embedding coordinates *y*_*i*_, the reconstructed expression is

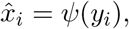

and the reconstructed velocity is

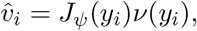

where *ψ* denotes the fitted manifold spline and *ν* the fitted velocity spline in embedding coordinates.

For each feature *j*, we quantify consistency using both the coefficient of determination (*R*^2^) and Pearson correlation (*r*) between observed and reconstructed values across cells.

For expression:

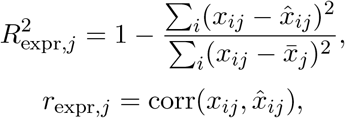

where

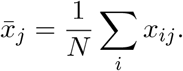

Because the manifold reconstruction is fit independently for each feature through a regression model with an affine component,

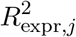

is typically non-negative and approaches 1 when the reconstructed manifold accurately captures the feature variation.

For velocity:

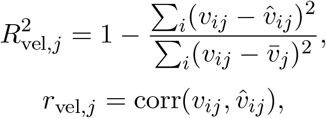

where

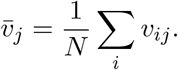

As described previously, the velocity reconstruction is obtained by first projecting observed velocities onto the local tangent space to obtain embedding-space velocities *u*_*i*_, fitting the smooth velocity spline *ν*, and then mapping the smoothed velocities back to expression space through the manifold Jacobian:

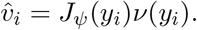

Because reconstructed velocities are constrained to lie in the tangent space of the manifold,

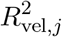

may become negative when the projected velocity explains less variance than the mean-centered baseline.

Genes or features with high consistency in both expression and velocity are interpreted as dynamically aligned with the reconstructed manifold, whereas features with low consistency are less compatible with the inferred vector field structure. Because velocity reconstruction depends on both the manifold geometry and the projected vector field, velocity consistency provides a stricter measure of dynamical coherence than expression reconstruction alone.

These quantities provide a feature-level diagnostic for identifying components that contribute most strongly to the learned dynamical geometry.

#### Gene-expression gradients and velocity alignment

Because FlowMap reconstructs a smooth mapping between the embedding space and the ambient gene-expression space, local gene-expression gradients can be computed directly from the manifold Jacobian.

Let

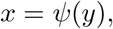

where

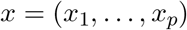

represents gene-expression coordinates and

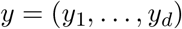

denotes the embedding coordinates.

The Jacobian of the manifold mapping is

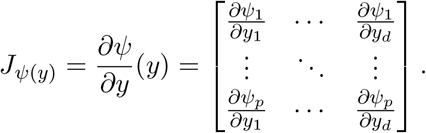

Each row of the Jacobian defines the local gradient of a gene-expression coordinate with respect to the embedding:

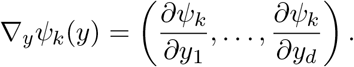

Because the embedding inherits a Riemannian metric from the reconstructed manifold, gene gradients are compared to the velocity field using the pullback metric

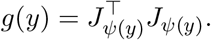

The corresponding Riemannian gradient is

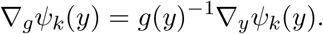

Given the fitted velocity spline *ν*(*y*), we quantify alignment between the local gene gradient and the inferred flow using the metric-aware cosine similarity

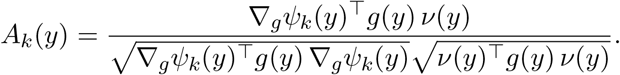

Positive alignment indicates that the gene increases along the direction of cellular progression, whereas negative alignment indicates decreasing expression along the flow. Values near zero correspond to genes whose local gradients are approximately orthogonal to the velocity field and therefore weakly associated with the local dynamical trajectory.

### Vector field geometry

#### RNA velocity as a flow on a Riemannian manifold

We model cellular dynamics as a continuous flow defined on a smooth, low-dimensional state manifold M. Let M be a *d*-dimensional manifold equipped with a Riemannian metric *g*, which defines inner products on the tangent space *T*_*x*_*M* at each point *x* ∈ *M*.

RNA velocity induces a vector field

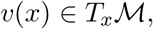

and the corresponding dynamics on the manifold satisfy

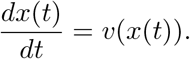

FlowMap represents the dynamics in embedding coordinates *y* ∈ ℝ^*d*^ using the fitted velocity spline

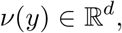

which defines the embedding-space dynamics

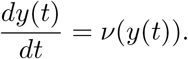

The manifold geometry is determined locally by the Jacobian of the reconstruction mapping *ψ*, which induces the pullback metric

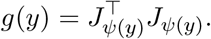

This metric describes how distances, angles, and local flow directions in the embedding correspond to the geometry of the reconstructed expression manifold. Consequently, even locally linear velocity fields in embedding coordinates may correspond to curved trajectories in expression space. Exact analysis of the dynamics therefore depends on both the velocity field and the local manifold geometry, making global solutions difficult to obtain in closed form.

To analyze the dynamics near fixed points, we therefore locally flatten the manifold geometry and approximate the flow using a locally Euclidean linear system.

#### Local flattening and linearization of the dynamics

To analyze the flow near a fixed point, we linearize the dynamics in locally Euclidean coordinates. The Riemannian metric *g*(*y*^∗^) induces a local inner product on 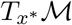. Let

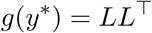

denote the Cholesky decomposition of the metric tensor. The transformation

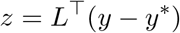

defines locally flattened coordinates in which the metric becomes the identity matrix. In these coordinates, the flow equation can be approximated by its first-order Taylor expansion,

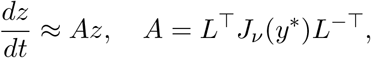

where *J*_*ν*_(*y*^∗^) is the Jacobian of the embedding velocity field evaluated at the fixed point. This procedure locally normalizes the metric structure, reducing the problem to a standard Euclidean linear system and yielding a stable representation of the local dynamics.

#### Classification of fixed points via linearized dynamics

The qualitative behavior of the flow near *x*^∗^ is determined by the eigenvalues of the linear operator *A*. In particular, the signs of the real parts characterize local stability, while complex eigenvalues indicate rotational or oscillatory behavior. The main classes of fixed points are summarized in Table 1.

**Table 1:**
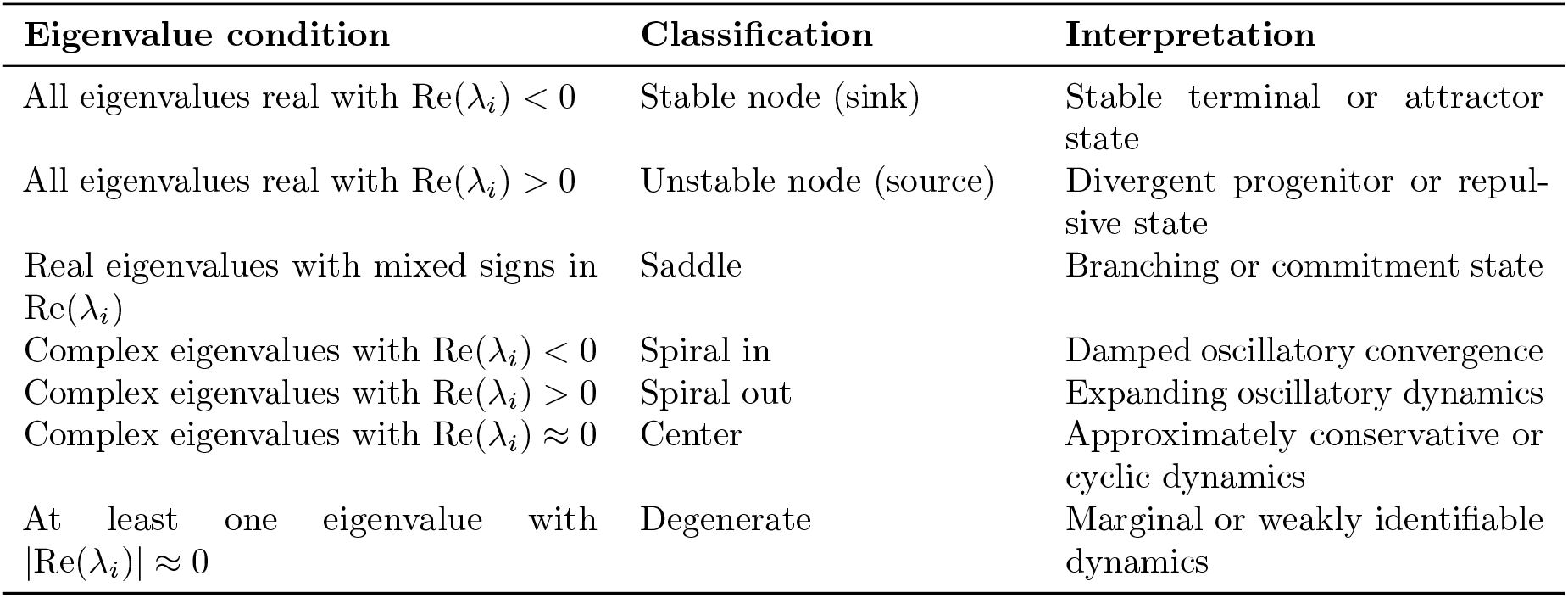
Classification of fixed points from the eigenvalues of the linearized operator *A*.

Because this classification is invariant under smooth local coordinate transformations, it provides an intrinsic and robust characterization of local cellular dynamics on the manifold.

#### Practical implementation via local Jacobian regression

In principle, both the Riemannian metric *g*(*x*) and the Jacobian of the coordinate vector field *J*_*u*_(*y*) can be computed in closed form from the fitted spline representations of the expression manifold and projected velocity field. These analytic derivatives provide exact local geometric quantities and are well defined everywhere on the reconstructed manifold. In practice, however, direct use of closed-form Jacobians leads to unstable estimates near fixed points. Because the vector field is inferred from noisy, sparsely sampled data, higher-order derivatives amplify local interpolation artifacts, resulting in irregular Jacobians that overfit small-scale fluctuations rather than capturing biologically meaningful dynamics.

To obtain robust estimates of local flow geometry, we therefore adopt a data-driven local linearization strategy. For each candidate fixed point *y*^∗^, we identify a neighborhood N(*y*^∗^) consisting of its *k* nearest neighbors in the embedding space. Within this neighborhood, the vector field is approximated by a first-order linear model,

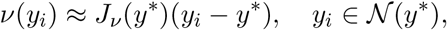

with the constraint that *u*(*y*^∗^) ≈ 0. We estimate the Jacobian *J*_*u*_(*y*^∗^) by solving a weighted least-squares regression problem,

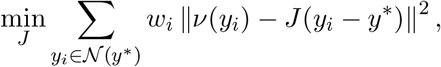

where the weights *w*_*i*_ decrease smoothly with distance from *y*^∗^. This local regression effectively smooths the Jacobian estimate, averages over measurement noise, and avoids over-interpreting high-frequency variation introduced by spline interpolation.

The resulting locally averaged Jacobian is then combined with the metric tensor at *y*^∗^ to perform the metric-aware linearization described above. This approach yields stable, interpretable estimates of fixed-point geometry while preserving the intrinsic structure of the underlying manifold and vector field.

### Least-action path formulation for trajectory inference

To infer differentiation trajectories connecting fixed points to terminal cell states, we adopt a least-action path (LAP) framework motivated by Dynamo [5]. Let

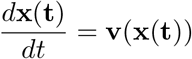

denote the ordinary differential equation (ODE) associated with the reconstructed RNA velocity vector field, where x(*t*) ∈ ℝ^*d*^ represents the cellular state at time *t*, and v(x) specifies the local direction and magnitude of state change. Solutions of this ODE generate flow curves (trajectories or paths) that describe continuous progression through the cellular state space.

In practice, observed cellular trajectories may deviate from ideal flow curves because of noise, stochasticity, sparse sampling, or imperfections in the estimated vector field. To identify trajectories that remain maximally consistent with the inferred dynamics, we define an action functional over candidate paths:

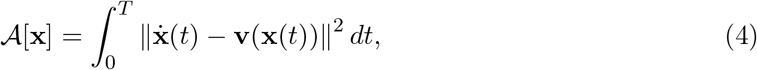

where 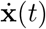 denotes the instantaneous tangent vector of the trajectory. This functional measures the discrepancy between the trajectory direction and the local velocity field along the path. Minimizing the action therefore favors trajectories that closely follow the vector field while remaining globally smooth and dynamically coherent.

Given a source state x_*s*_ and a target state x_*t*_, corresponding to fixed points or terminal cellular states, the least-action path is defined as the solution of the variational problem

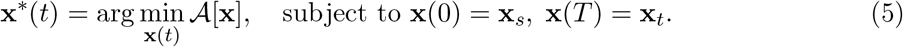

Numerically, the trajectory is discretized into a finite set of control points, and the action is minimized using iterative gradient-based optimization. The resulting path approximates the most dynamically consistent transition trajectory between the two cellular states under the reconstructed vector field.

In FlowMap, least-action trajectories are computed directly within the low-dimensional geometry-preserving embedding. The velocity field entering the action functional is the smooth tangent-space–constrained vector field reconstructed from the FlowMap manifold. Consequently, inferred trajectories remain consistent with both the manifold geometry and the denoised continuous dynamics represented by the embedding.

### Curvature analysis

#### Decomposition of the acceleration vector for RNA-velocity

Let

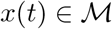

denote a trajectory on the cellular state manifold evolving according to the RNA velocity field

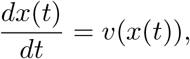

where

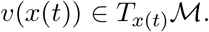

Differentiating along the trajectory gives the acceleration vector

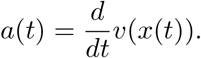

Let

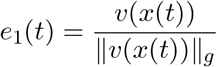

denote the unit flow direction, and let

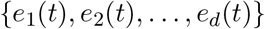

be an orthonormal basis for the tangent space

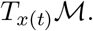

Because the manifold is embedded in the ambient gene-expression space ℝ^*p*^, the tangent space admits an orthogonal complement given by the normal space. Let

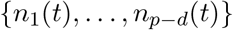

be an orthonormal basis for the normal space of the embedded manifold at *x*(*t*).

The acceleration vector admits the orthogonal decomposition [31]

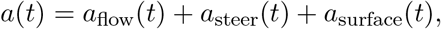

where

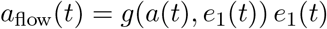

is the component parallel to the local flow direction,

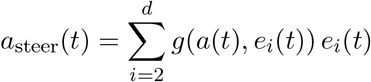

is the component tangent to the manifold but orthogonal to the flow direction, and

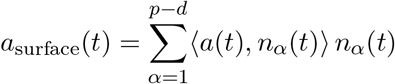

is the component orthogonal to the embedded manifold in the ambient expression space.

The flow component *a*_flow_ measures local acceleration or deceleration along the trajectory. The steering component *a*_steer_ captures directional turning of the flow within the manifold and reflects intrinsic curvature of the cellular dynamics. The surface component *a*_surface_ measures deviation of the dynamics away from the locally embedded manifold surface in the ambient expression space.

Because these components are mutually orthogonal,

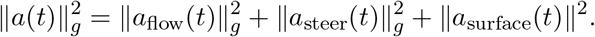

#### Interpretation of the acceleration components

Each term in the decomposition of the acceleration vector captures a distinct aspect of the local dynamics of an embedded flow curve.

The flow component *a*_flow_(*t*) lies along the instantaneous velocity direction and quantifies changes in the *speed* of progression along the trajectory. A nonzero *a*_flow_(*t*) indicates local acceleration or deceleration along the same path without altering the direction of motion. Consequently, this component reflects temporal rescaling of the dynamics rather than geometric bending of the trajectory in state space.

The steering component *a*_steer_(*t*) lies within the tangent space of the manifold but is orthogonal to the flow direction. It captures changes in direction that occur *along the manifold itself*, corresponding to steering or turning within the intrinsic geometry of the learned surface. When *a*_steer_(*t*) = 0, the trajectory follows a geodesic of the manifold, representing the straightest possible path constrained to the surface. Nonzero values of *a*_steer_(*t*) therefore reflect intrinsic curvature of the flow induced by the underlying biological dynamics.

The surface component *a*_surface_(*t*) is orthogonal to the manifold and measures how the trajectory bends relative to the ambient gene-expression space in which the manifold is embedded. This term captures extrinsic curvature arising from the geometry of the reconstructed manifold itself rather than from directional changes within the surface. Intuitively, *a*_surface_(*t*) reflects how the trajectory responds to the “terrain” of the manifold, analogous to motion over a curved landscape embedded in a higher-dimensional space.

In practice, low-dimensional RNA-velocity visualizations primarily reveal the tangent components of the dynamics, namely the flow and steering terms. The flow component describes local acceleration or deceleration along trajectories, while the steering component appears visually as turning or redirection of the velocity field within the embedding. By contrast, the surface component cannot generally be identified directly from the embedding visualization alone. Because the embedding represents a curved manifold projected into a low-dimensional coordinate system, deviations orthogonal to the manifold are not visually separable from the curvature of the surface itself. Recovering the surface component therefore requires an explicit differentiable reconstruction of the embedding manifold together with its tangent and normal spaces, which in FlowMap is provided by the spline reconstruction.

Together, these components disentangle speed modulation, intrinsic steering, and extrinsic bending of cellular trajectories, providing a geometrically interpretable decomposition of local dynamics in the FlowMap embedding.

#### Computing acceleration components

Let

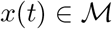

denote a trajectory on the cellular state manifold, and let

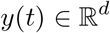

denote its embedding coordinates. FlowMap represents the manifold using a smooth mapping

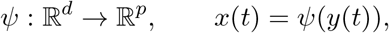

where *p* is the dimension of the ambient gene-expression space.

The dynamics on the manifold satisfy

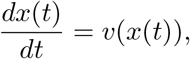

while the projected dynamics in embedding coordinates satisfy

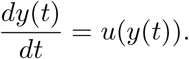

The velocity in expression space is therefore

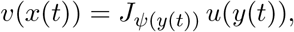

where

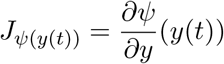

is the Jacobian of the manifold mapping evaluated at *y*(*t*).

The ambient acceleration vector is

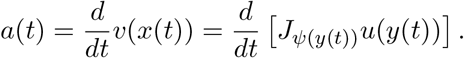

Equivalently, for each expression coordinate *k* = 1, …, *p*,

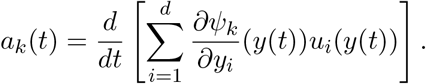

Applying the product rule gives

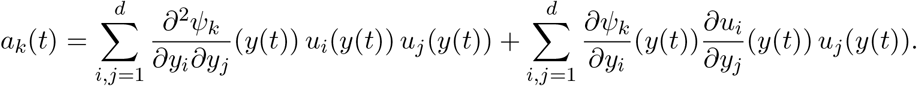

The first term captures acceleration induced by local curvature of the reconstructed expression manifold, while the second term captures local variation of the embedded velocity field.

To decompose the acceleration vector, we first project it onto the tangent space of the embedded manifold:

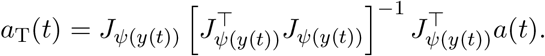

The surface component is defined as the residual orthogonal to the tangent space,

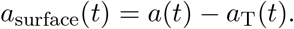

Let

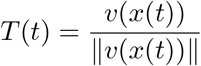

denote the unit flow direction in expression space. The flow component is

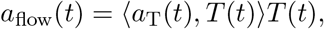

and the steering component is

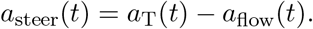

Thus, the acceleration decomposes as

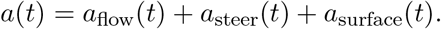

#### Curvature and acceleration

Curvature quantifies how rapidly a trajectory changes direction along the flow, after removing changes in speed. The acceleration of an embedded flow curve decomposes into (motivated in Supplementary Note 3.1)

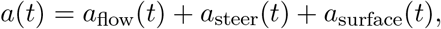

where *a*_flow_ is parallel to the velocity, *a*_steer_ is tangent to the manifold but orthogonal to the velocity, and *a*_surface_ is normal to the reconstructed manifold.

Let

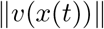

denote the magnitude of the velocity vector in expression space. Since curvature measures directional change rather than acceleration along the same direction, the flow component is not included in the curvature. The steering curvature is defined as

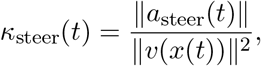

which measures turning of the trajectory within the manifold. The surface curvature is defined as

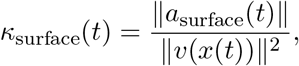

which measures bending of the trajectory induced by the embedding of the reconstructed manifold in ambient expression space.

Because *a*_steer_ and *a*_surface_ are orthogonal, the total curvature of the ambient trajectory is

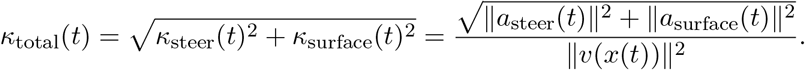

Thus, curvature separates intrinsic directional steering of the cellular flow from extrinsic bending of the reconstructed expression manifold. The flow component *a*_flow_ instead measures speed modulation along the same trajectory and is therefore reported separately from curvature. Additional geometric motivation and the relationship to intrinsic and extrinsic surface curvature are provided in Supplementary Notes 3.1–3.2.

Normalization by the squared velocity magnitude removes dependence on traversal speed: faster motion along the same geometric path produces larger acceleration but not larger curvature. Consequently, curvature reflects how sharply the trajectory bends in expression space rather than the absolute magnitude of transcriptional change.

### Data preprocessing and implementation

FlowMap is implemented in Python using standard scientific computing libraries, including NumPy, SciPy, scikit-learn, Scanpy, and scVelo. The core components of the method, including spline fitting, Jacobian computation, tangent-space projection, and velocity mapping, admit closed-form solutions and can therefore be computed efficiently without specialized hardware acceleration. All analyses in this study were performed on standard desktop and laptop hardware without GPU acceleration. Unless otherwise specified, FlowMap was applied to PCA representations computed from the processed expression matrices.

The cell cycle [25], pancreas [27], hematopoiesis [28], and B cell activation [33] datasets were processed using the standard scVelo stochastic workflow. Briefly, gene expression matrices were normalized and log transformed, highly variable genes were selected, and principal component analysis and neighborhood graph construction were performed using Scanpy. RNA velocity was then estimated using the stochastic model implemented in scVelo with default parameters.

For the Larry hematopoiesis dataset [28], fate probabilities were estimated using CoSpar [46] based on available clonotype lineage information. For the spatial transcriptomics dataset [36], velocity estimates were obtained using SIRV with default parameters. Complete analysis scripts and figure-generation workflows are provided in the public code repository.

### Benchmark

For simulated vector field benchmarks, we evaluated agreement between the ground-truth dynamics and the corresponding low-dimensional embeddings using standard metrics for neighborhood preservation and vector-field consistency. Embedding quality was assessed using trustworthiness [47] and neighborhood overlap measured using the Jaccard index [48]. Vector-field consistency was evaluated using velocity magnitude correlation, local cosine similarity between neighboring velocity vectors, and smoothness metrics based on velocity differences between neighboring cells. For benchmark comparisons, FlowMap, scVelo [4], Dynamo [5], VeloViz [24], and GraphVelo [14] were run using default parameters, with the simulated datasets providing only the expression matrix and velocity matrix as inputs.

For real RNA velocity datasets, we performed three categories of benchmark comparisons. In the first setting, both the RNA velocity estimates and cell embeddings were fixed, allowing comparison of vector-field reconstruction and projection independently of the embedding geometry. In the second setting, RNA velocity estimates were fixed while methods were allowed to construct both the cell embedding and vector-field representation, enabling evaluation of the joint embedding and velocity structure. For these two benchmark settings, RNA velocity was estimated using the stochastic mode implemented in scVelo. Cell cycle and hematopoiesis embeddings were generated using FlowMap, whereas pancreas and dentate gyrus embeddings were obtained from the standard scVelo workflow. Unless otherwise specified, FlowMap, scVelo, and Dynamo were run using default embedding parameters.

In the third benchmark setting, cell embeddings were fixed while different RNA velocity estimation methods were compared. Specifically, velocity estimates generated using the stochastic and dynamical modes of scVelo were compared against Dynamo-based velocity estimates.

## Data Availability

All datasets used in this study are publicly available and were previously published in the corresponding original studies. We used either publicly available processed datasets or standard preprocessing workflows implemented in Scanpy and scVelo to ensure consistency across benchmarks.

| Dataset | Reference | Processed data source | Velocity estimation |
| --- | --- | --- | --- |
| Cell cycle | [25] | Dynamo | scVelo |
| Pancreas | [27] | scVelo | scVelo |
| Larry hematopoiesis | [28] | PyroVelocity | scVelo |
| B cell activation | [33] | In-house Scanpy and scVelo | scVelo |
| Mouse organogenesis | [36] | SIRV | SIRV |

## Code Availability

The FlowMap software package is available at: https://github.com/jingyuanhu01/flowmap. The documentation and tutorial are available at: https://flowmap.readthedocs.io/en/latest/index.html. The manuscript source files and analysis scripts used to generate the figures and results are available at: https://github.com/jingyuanhu01/flowmap-manuscript.

## Acknowledgements

This work was supported by the National Human Genome Research Institute (NHGRI) grant U01HG012041 (to H.S. and J.D.) and by the National Science Foundation (NSF) grant DMS-2515684 (to R.M.).

## Supplementary Tables

**Table S1:** Comparison of capabilities across RNA velocity and dynamical modeling frameworks. FlowMap focuses on geometry-aware embedding construction and downstream dynamical analysis, while Dynamo, scVelo, and Velocyto emphasize velocity and kinetic parameter estimation. ^∗^Dynamo models a differentiable vector field in gene or PCA space, but does not define a differentiable embedding manifold.

| Capability | FlowMap | Dynamo | scVelo | Velocyto |
| --- | --- | --- | --- | --- |
| **Velocity Modeling** |  |  |  |  |
| Velocity estimation | × | ✓ | ✓ | ✓ |
| Kinetic parameter estimation | × | ✓ | ✓ | × |
| **Embedding Construction** |  |  |  |  |
| Differentiable manifold | ✓ | ✓* | × | × |
| Embedding reconstruction & denoising | ✓ | × | × | × |
| Embedding quality evaluation | ✓ | × | × | × |
| **Dynamical Analysis** |  |  |  |  |
| Gene expression–visualization linkage | ✓ | × | × | × |
| Marker identification from embedding | ✓ | × | × | × |
| Pseudotime | ✓ | ✓ | ✓ | × |
| Least Action Paths (LAP) | ✓ | ✓ | × | × |
| Fixed-point identification | ✓ | ✓ | × | × |
| **Gene Regulatory Inference** |  |  |  |  |
| Regulation analysis | × | ✓ | × | × |
| Perturbation simulation | × | ✓ | × | × |

## Supplementary Figures

**Table S2:**
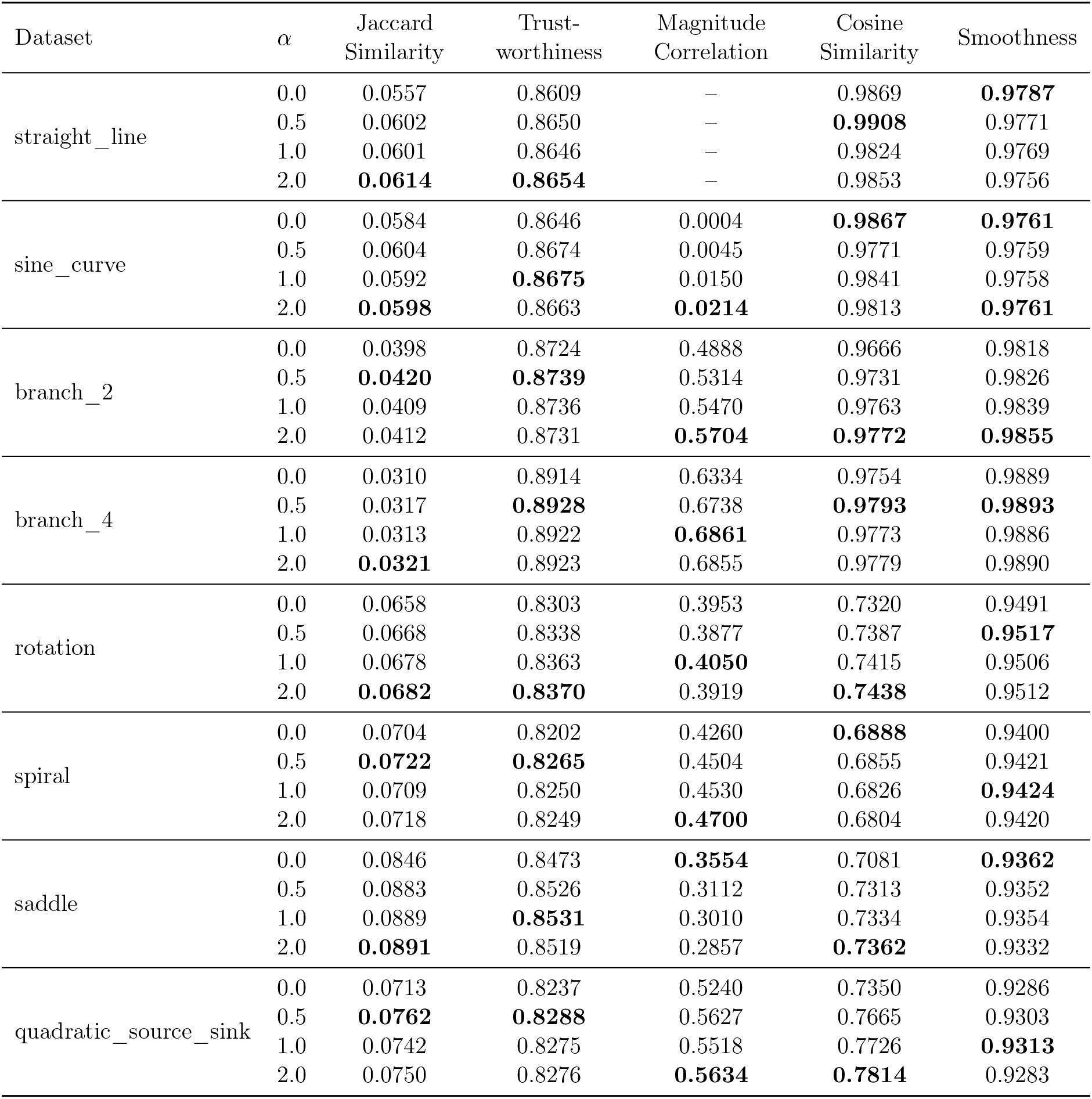
Effect of phase-distance weighting on FlowMap embeddings. Simulation benchmark across synthetic dynamical systems using different phase-distance weighting strengths *α*. Bold indicates the best metric value across *α* for each dataset.

**Table S3:**
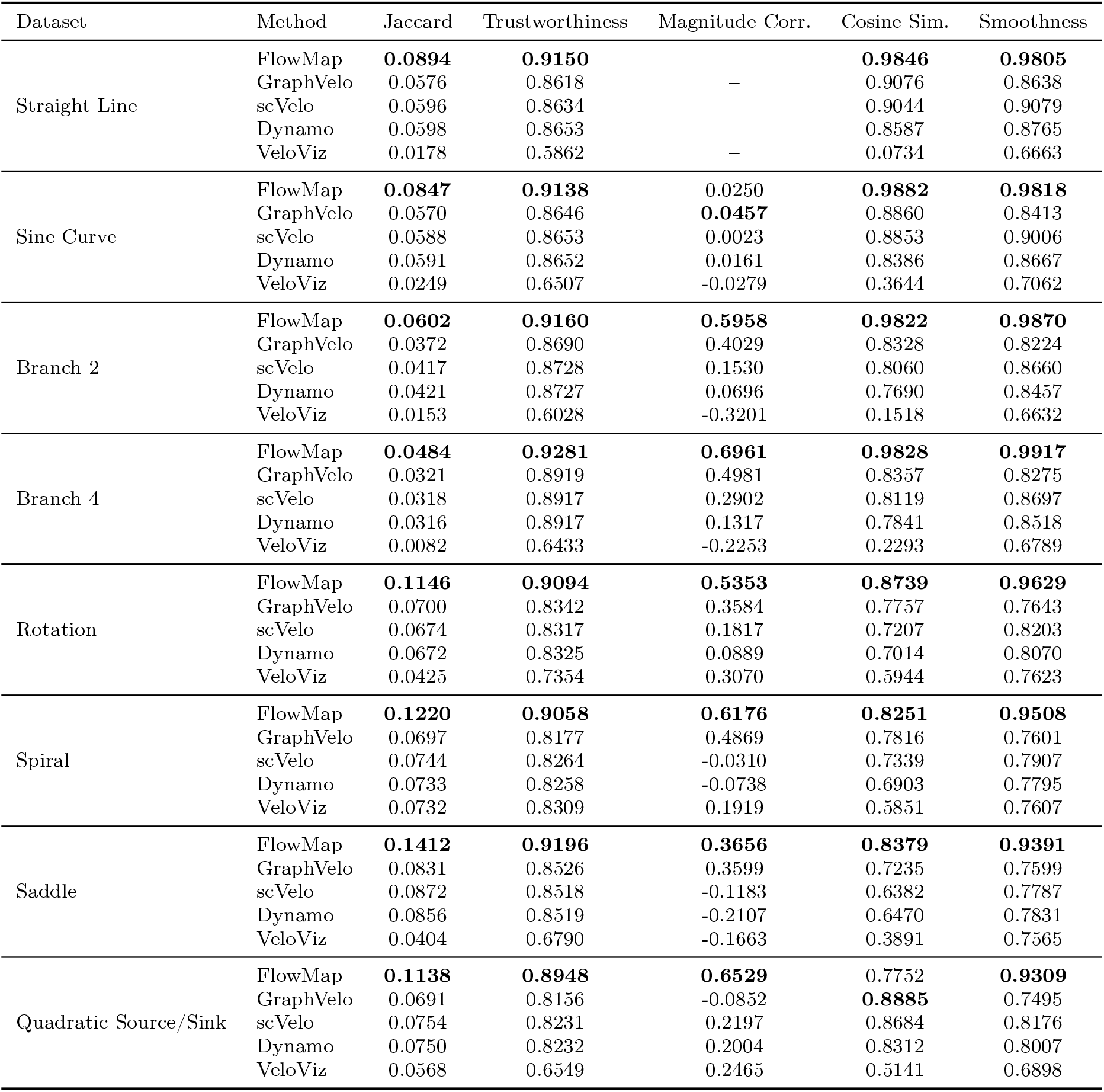
Benchmarking results on simulated dynamical systems. Bold indicates the best-performing method for each metric within each dataset.

**Figure S1:**
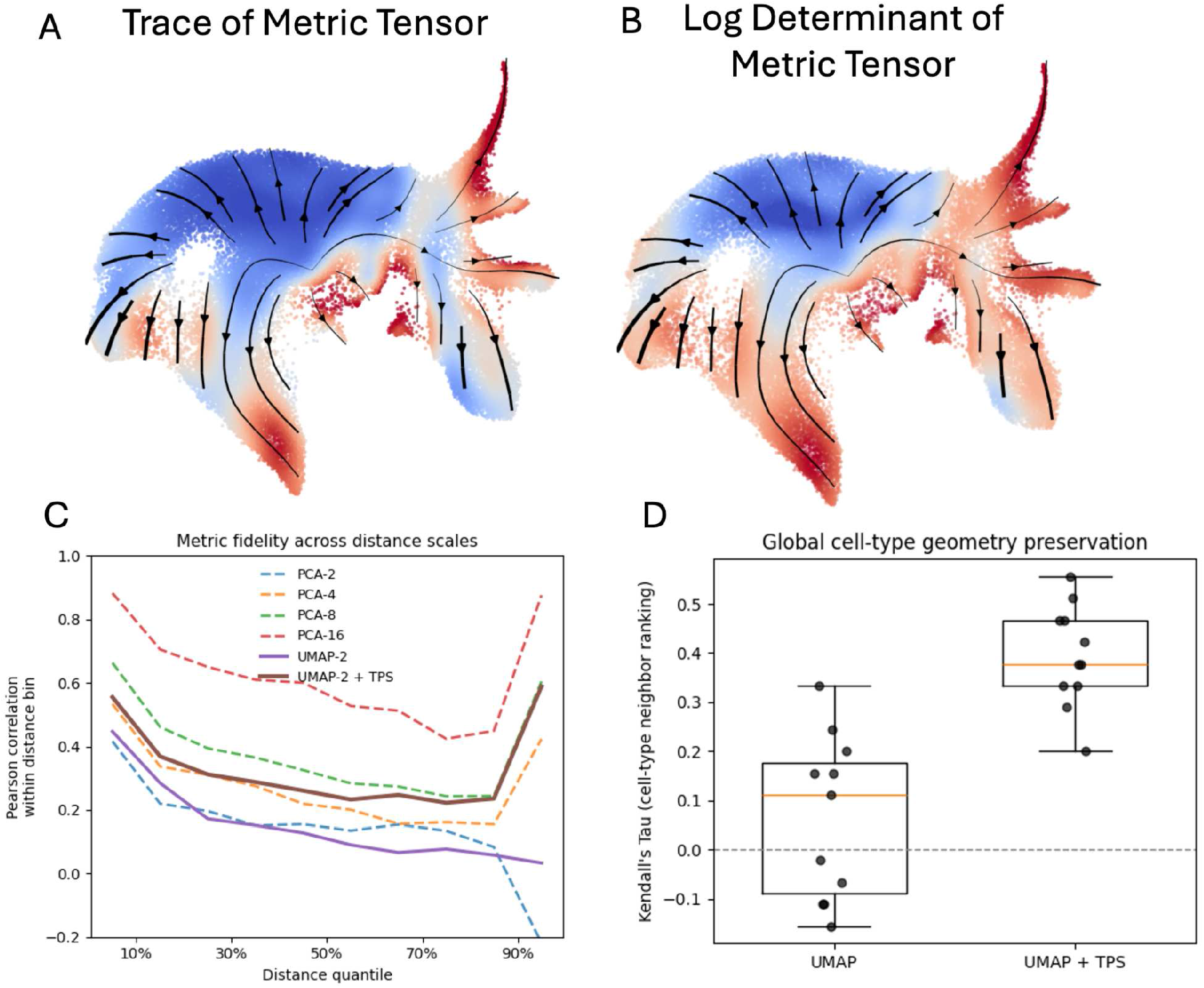
Geometry-aware correction of 2D embeddings using TPS. (A,B) Scalar summaries of the induced metric tensor on the embedding, visualized as the trace (A) and log-determinant (B). These quantities highlight spatially heterogeneous stretching and compression across the embedding, indicating that geometric distortion is not uniform. (C) Metric fidelity across distance scales. For each cell, pairwise distances to all other cells were computed in the ambient space and in each embedding. Distances were binned by ambient-space distance quantiles, and Pearson correlation between ambient and embedded distances was computed within each bin. UMAP exhibits rapid degradation of distance fidelity beyond short ranges, whereas UMAP+TPS improves correlations across intermediate and long distances, approaching the performance of low-dimensional PCA (PCA-4 to PCA-8). (D) Global cell-type geometry preservation measured by Kendall’s Tau correlation of cell-type neighbor rankings, following Chari and Pachter. TPS correction substantially improves preservation of global cell-type relationships relative to UMAP alone.

**Figure S2:**
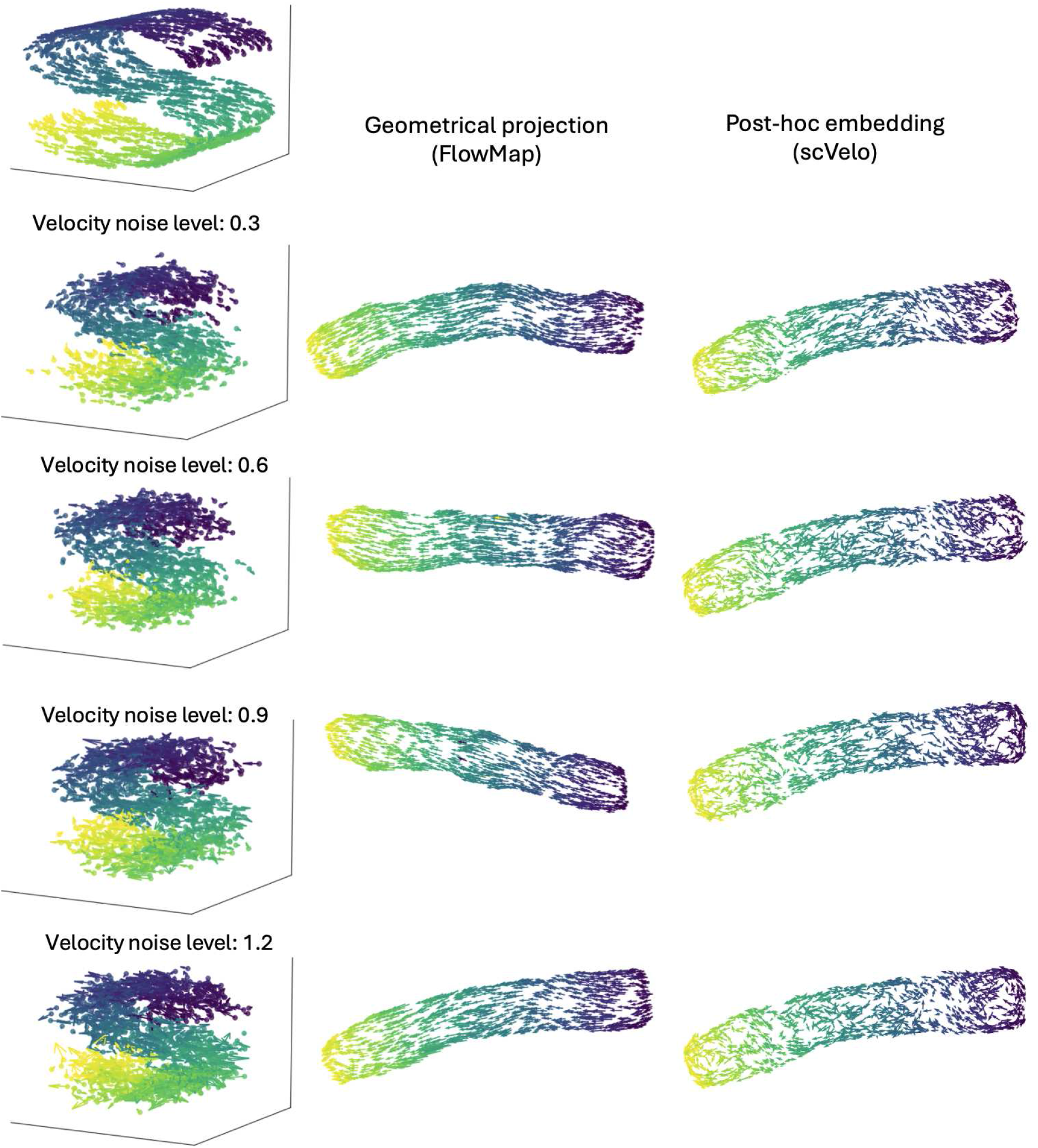
Geometric projection is necessary for stable velocity visualization under noise. *Left:* Ground-truth three-dimensional S-curve with increasing levels of additive velocity noise (top to bottom), colored by latent phase. *Middle (FlowMap):* Velocity vectors projected onto the learned manifold using a geometric (tangent-space) constraint, yielding coherent and stable flows even at high noise levels. *Right (scVelo):* Post-hoc velocity embedding obtained without enforcing geometric consistency, resulting in distorted and unstable vector fields as noise increases.

**Figure S3:**
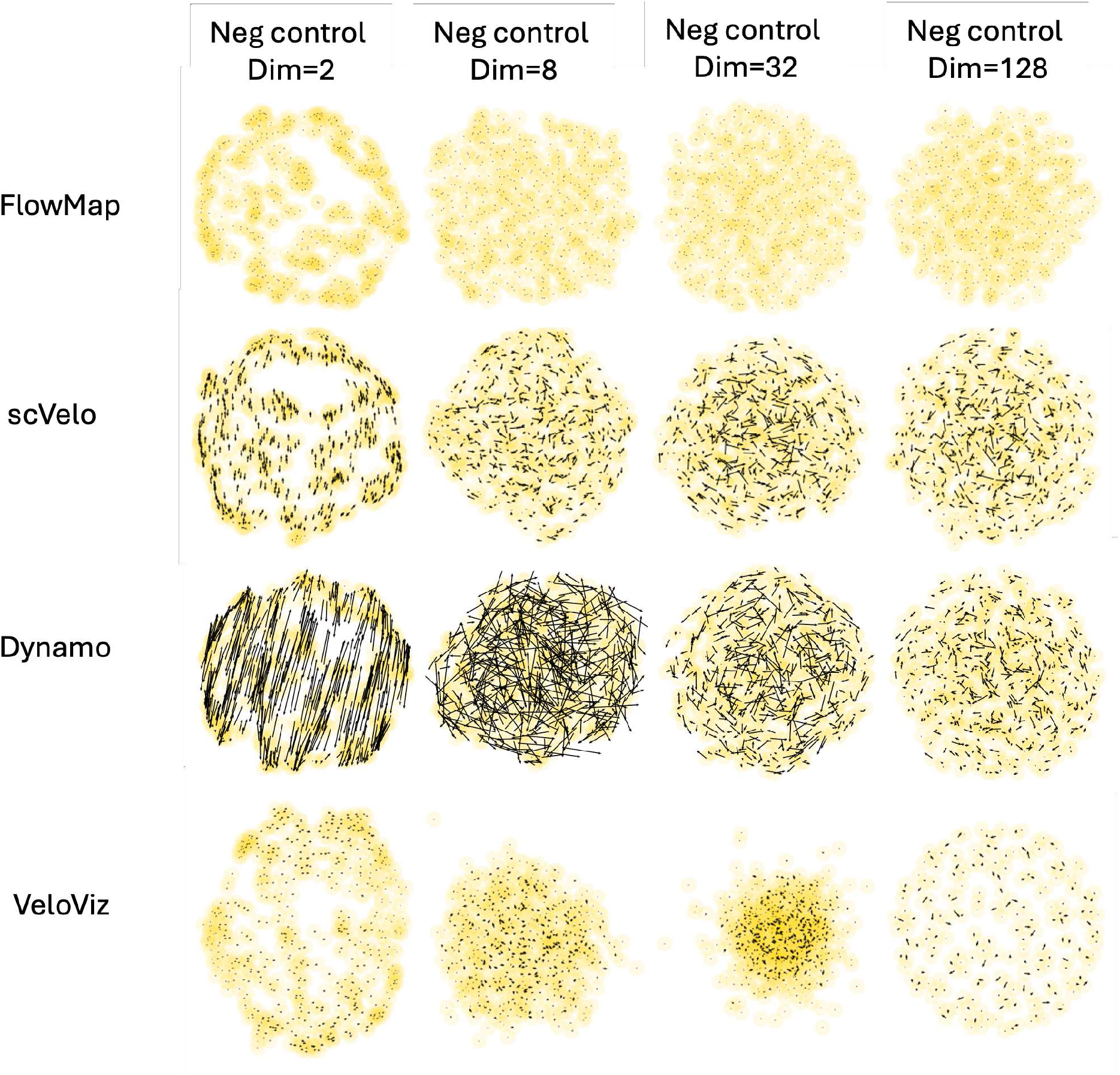
Negative control for embedding-induced velocity artifacts. RNA velocity vectors were generated as independent Gaussian noise in the original expression space at varying input dimensionalities (columns: Dim = 2, 8, 32, 128). Cells were embedded into two dimensions and velocities were visualized using four embedding strategies (rows). Despite the absence of any underlying dynamical structure, neighborhood-based embedding approaches produce visually coherent flow patterns that depend on the embedding geometry and smoothing procedure. In contrast, geometry-constrained projection suppresses these spurious patterns, yielding near-zero embedded velocities consistent with the absence of true dynamics. This illustrates that apparent low-dimensional flow structure can arise purely from embedding and interpolation effects, and that enforcing geometric consistency mitigates such artifacts.

**Figure S4:**
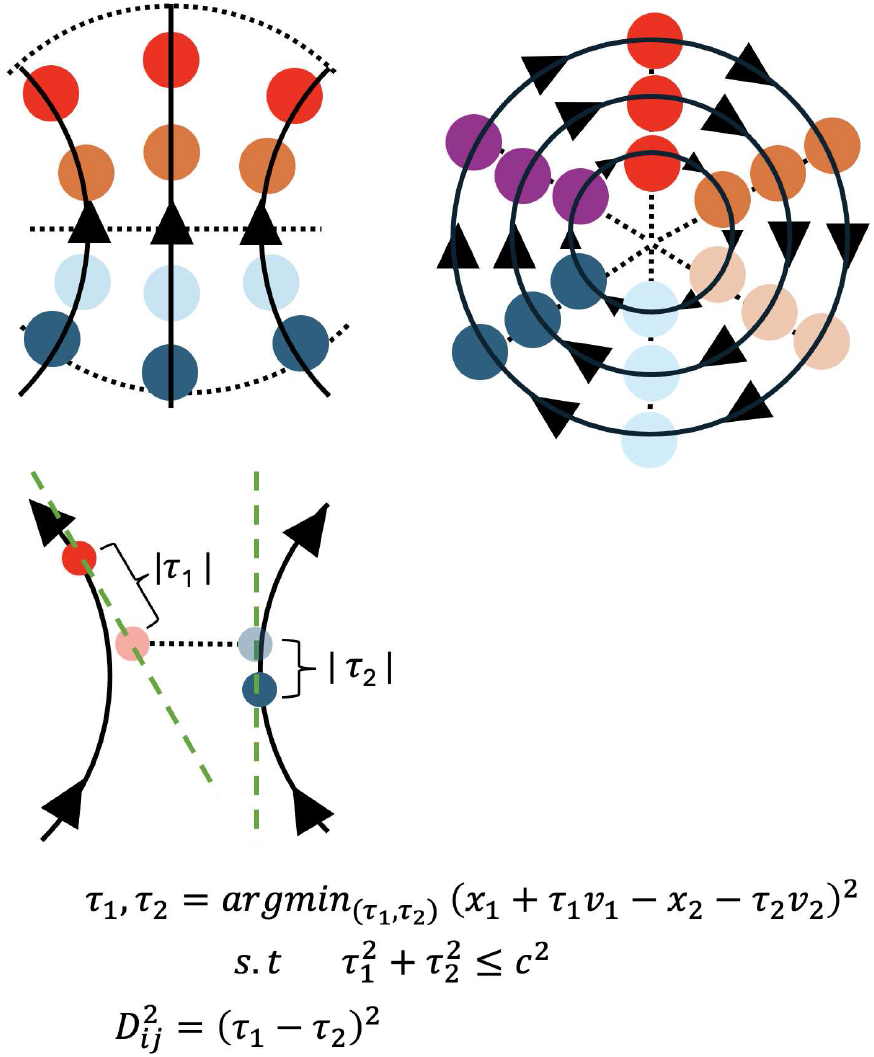
Conceptual illustration of phase along RNA velocity flow. Three examples of RNA velocity fields with labeled phase. Cells with the same color are at the same phase and lie on curves perpendicular to the local velocity field, representing equivalent positions along a continuous trajectory. Although such cells may differ in expression, they are synchronized in their progression along the flow. For nearby cells, phase is estimated by allowing each cell to move along its velocity direction and solving for the relative offsets (*τ*_1_, *τ*_2_) that best align their expression states. The resulting phase distance 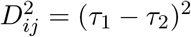 captures separation along the flow direction rather than transcriptional similarity alone.

**Figure S5:**
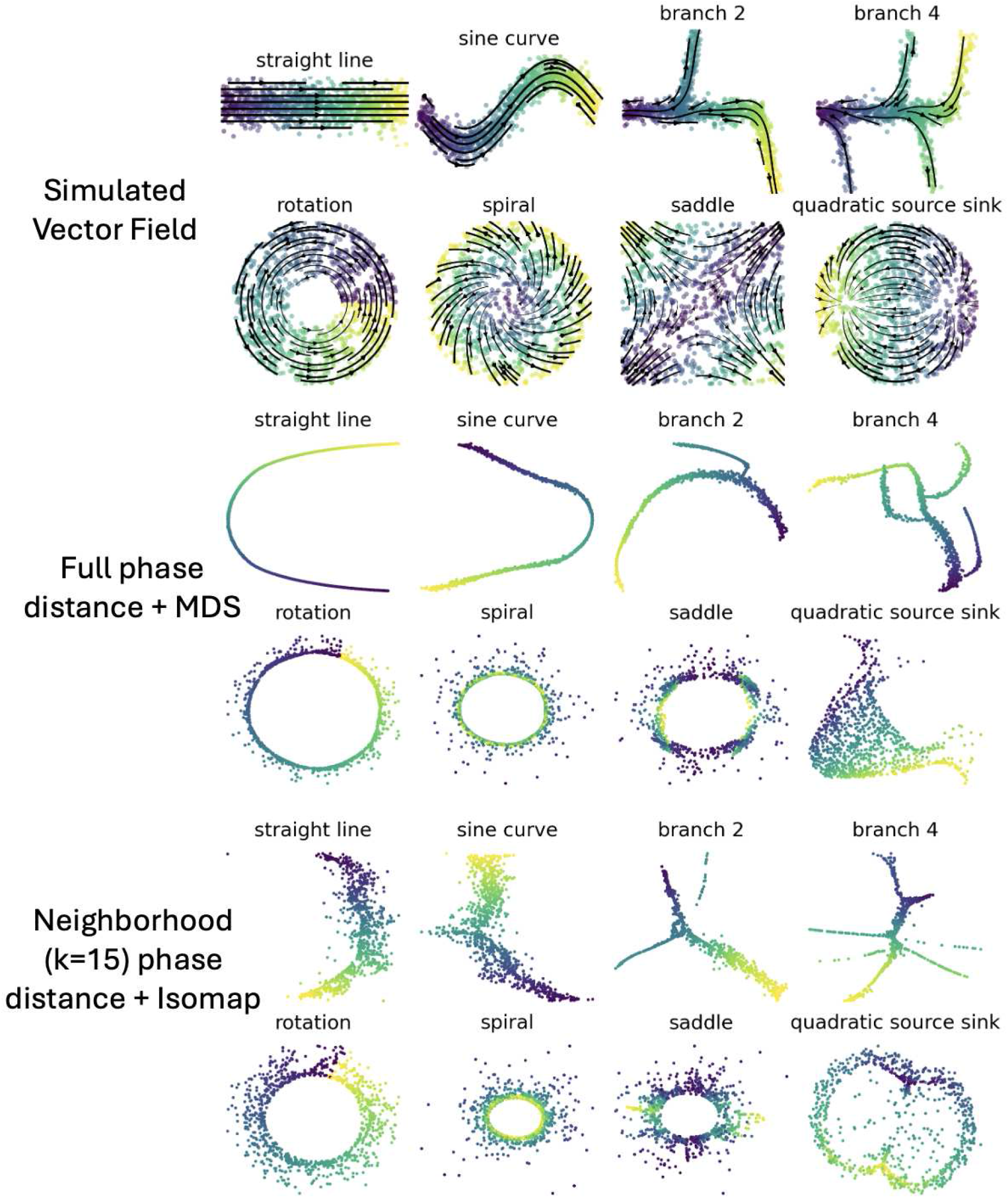
Phase distance captures cellular position along flow curves in simulated vector fields. **Top:** Simulated vector fields used for benchmarking, including linear, nonlinear, branching, rotational, spiral, saddle, and source–sink dynamics. Points are colored by true temporal position along the flow. **Middle:** Embeddings obtained using the full pairwise phase distance matrix followed by multidimensional scaling (MDS). Phase distance alone is sufficient to recover coherent low-dimensional representations that reflect progression along flow curves, straightening trajectories and separating branches according to their dynamical ordering. **Bottom:** Embeddings constructed using neighborhood-based phase distance (*k* = 15) and an Isomap-style procedure, where global distances are computed as shortest paths on the phase-distance graph.

**Figure S6:**
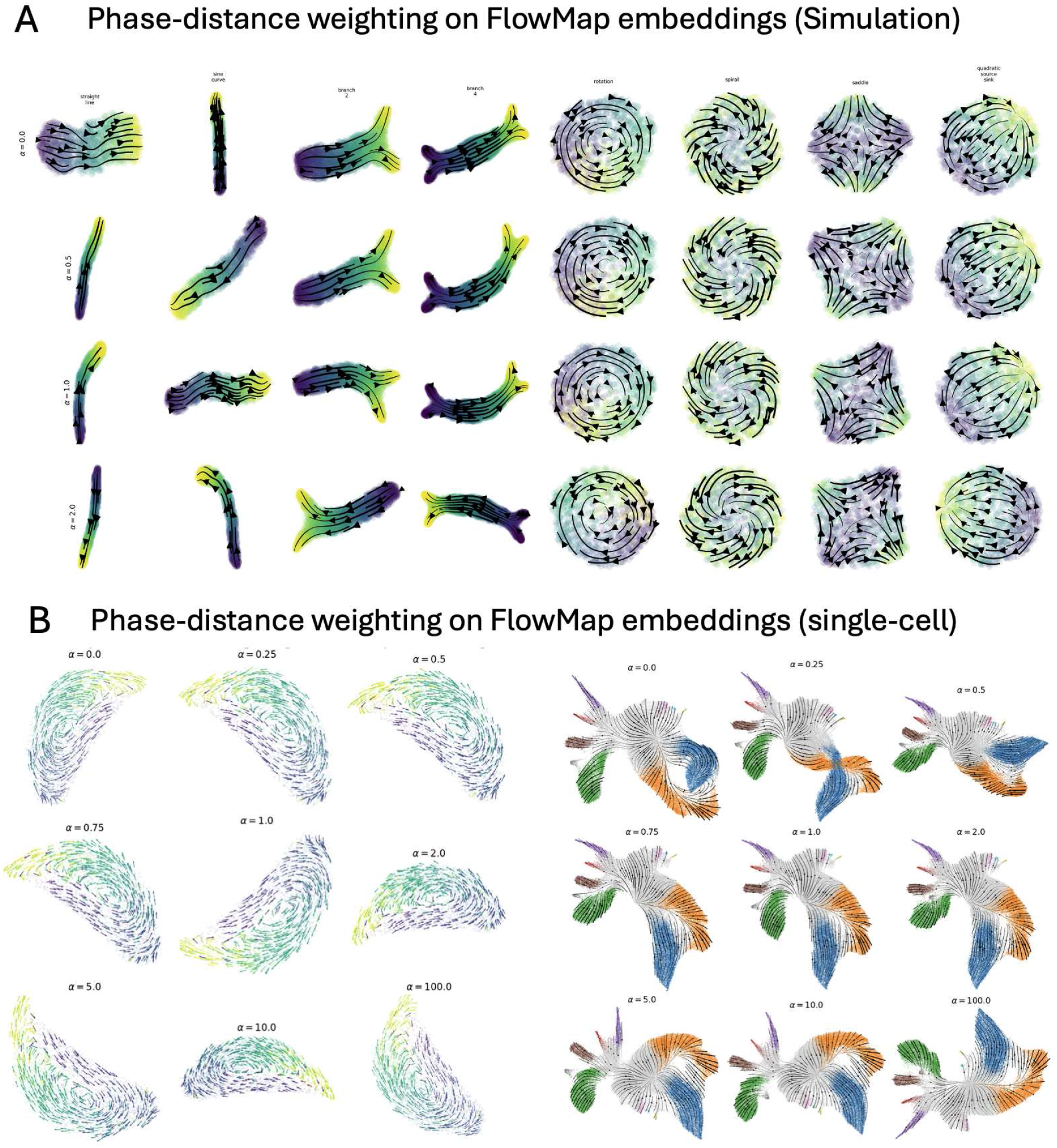
Phase-distance weighting on FlowMap embeddings. (A) Simulated vector field benchmark across eight synthetic dynamical systems. Rows correspond to increasing phase-distance weighting strength *α*, where the embedding distance is defined as Euclidean distance plus an additional phase-distance term. (B) Phase-distance weighting on real single-cell datasets. Left: cell-cycle dynamics. Right: hematopoietic differentiation in the LARRY dataset.

**Figure S7:**
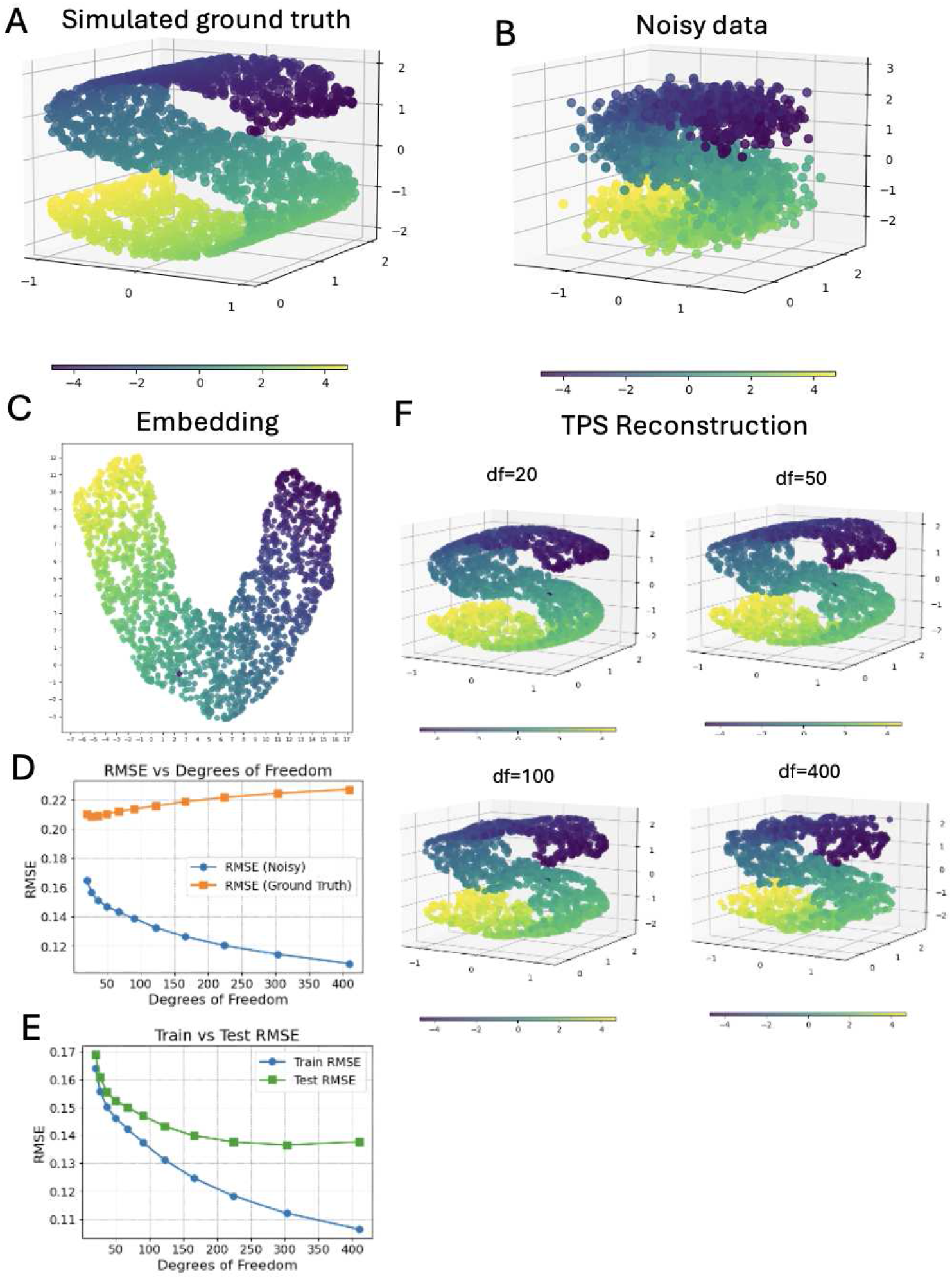
Thin-plate spline (TPS) surface reconstruction with varying degrees of freedom. (A) Simulated ground-truth manifold. (B) Noisy observations sampled from the same manifold. (C) Low-dimensional embedding constructed from the noisy data. (D) Reconstruction RMSE as a function of TPS degrees of freedom, evaluated against noisy data and ground truth. (E) Train and test RMSE as a function of degrees of freedom. (F) TPS reconstructions of the manifold for different degrees of freedom.

**Figure S8:**
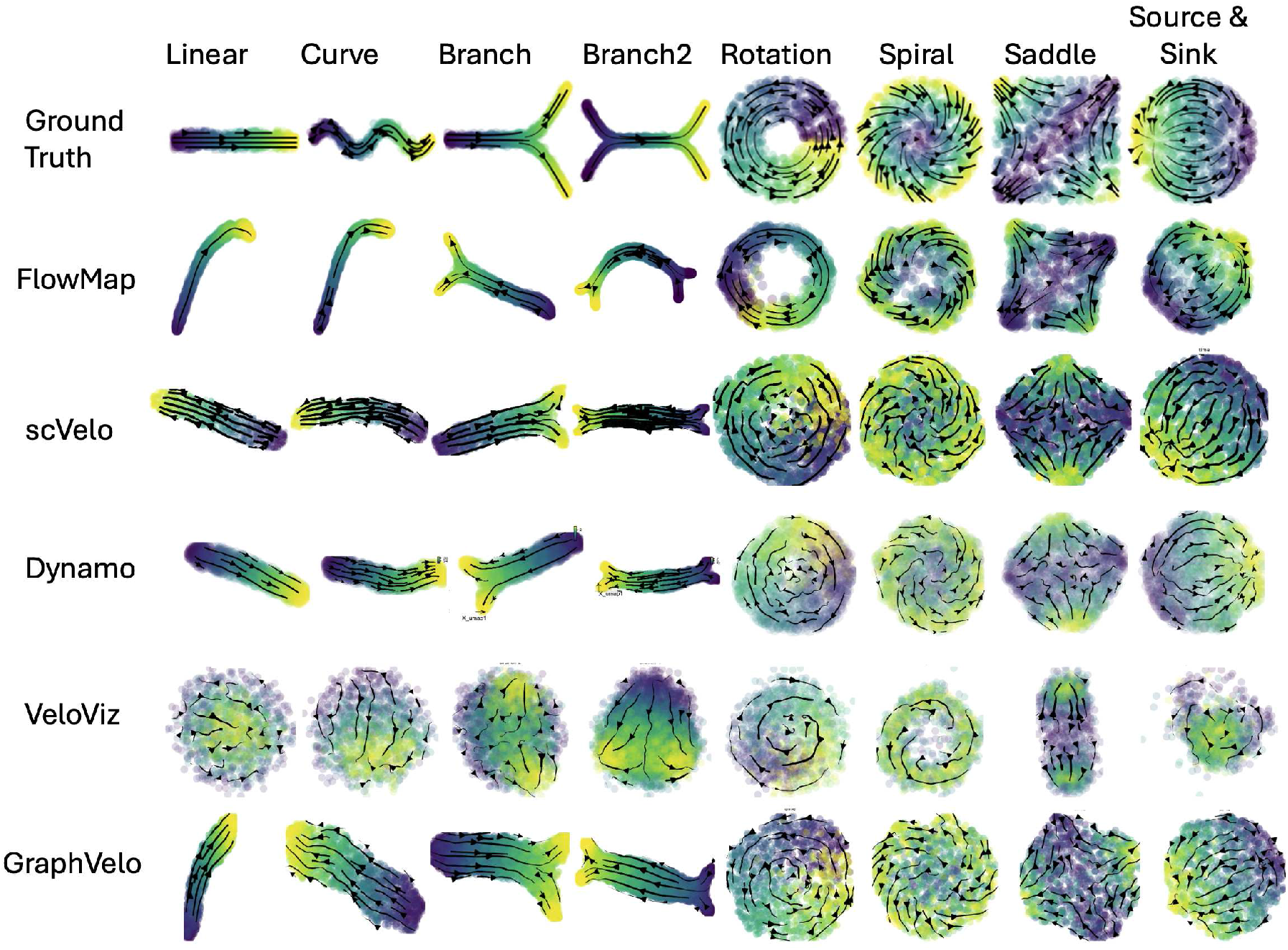
Simulation benchmark of FlowMap across diverse dynamical systems. Synthetic vector fields with linear, branching, and cyclic dynamics were used to evaluate embedding and velocity recovery. Cells are colored by ground-truth trajectory position, with arrows indicating velocity.

**Figure S9:**
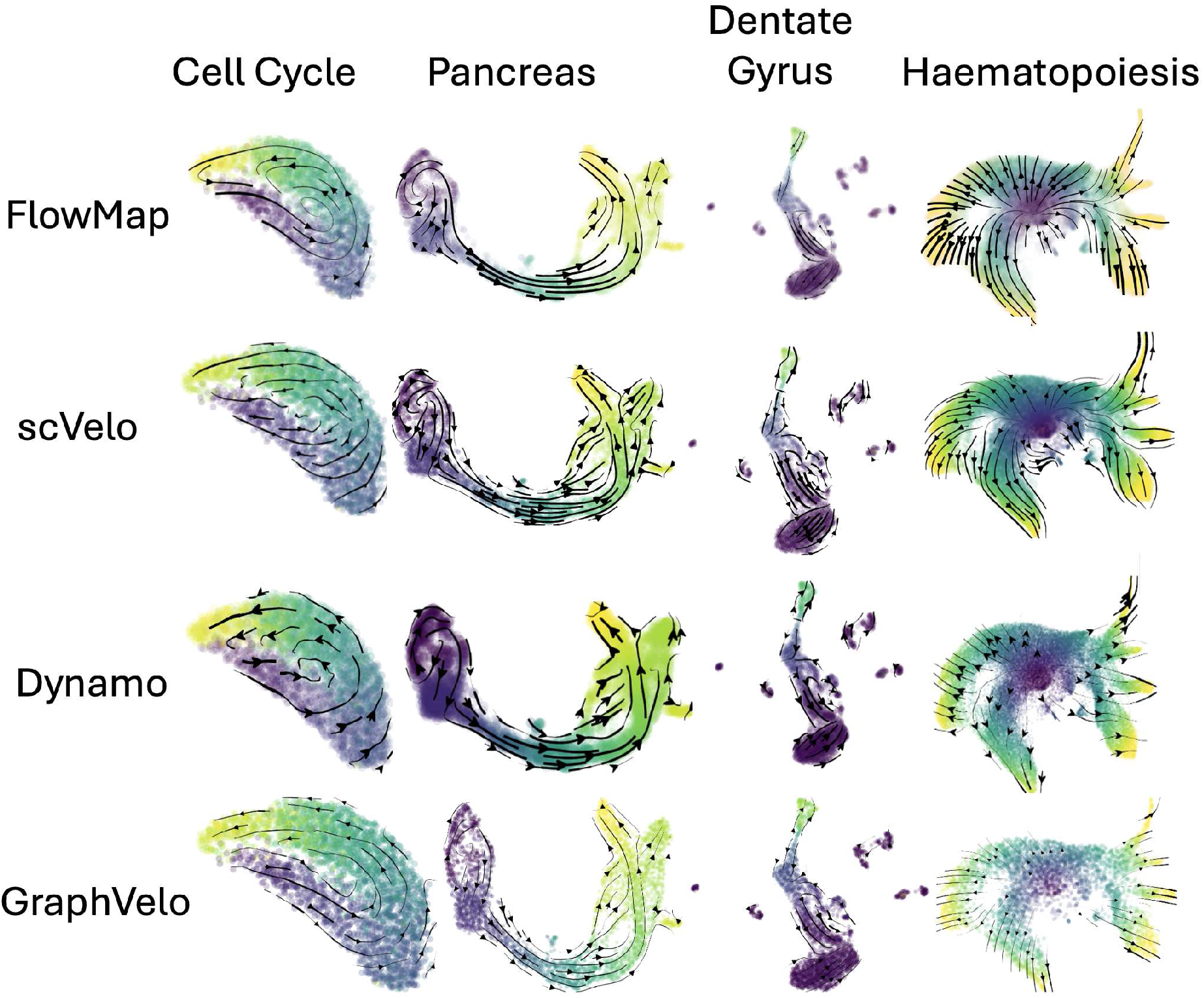
Benchmark of velocity field reconstruction on real single-cell datasets. Low-dimensional embeddings with overlaid RNA velocity fields for representative datasets exhibiting diverse dynamical structures, including continuous, curved, and branching trajectories. Each row corresponds to a different method or condition, illustrating variations in the coherence and structure of inferred velocity fields across embeddings.

**Figure S10:**
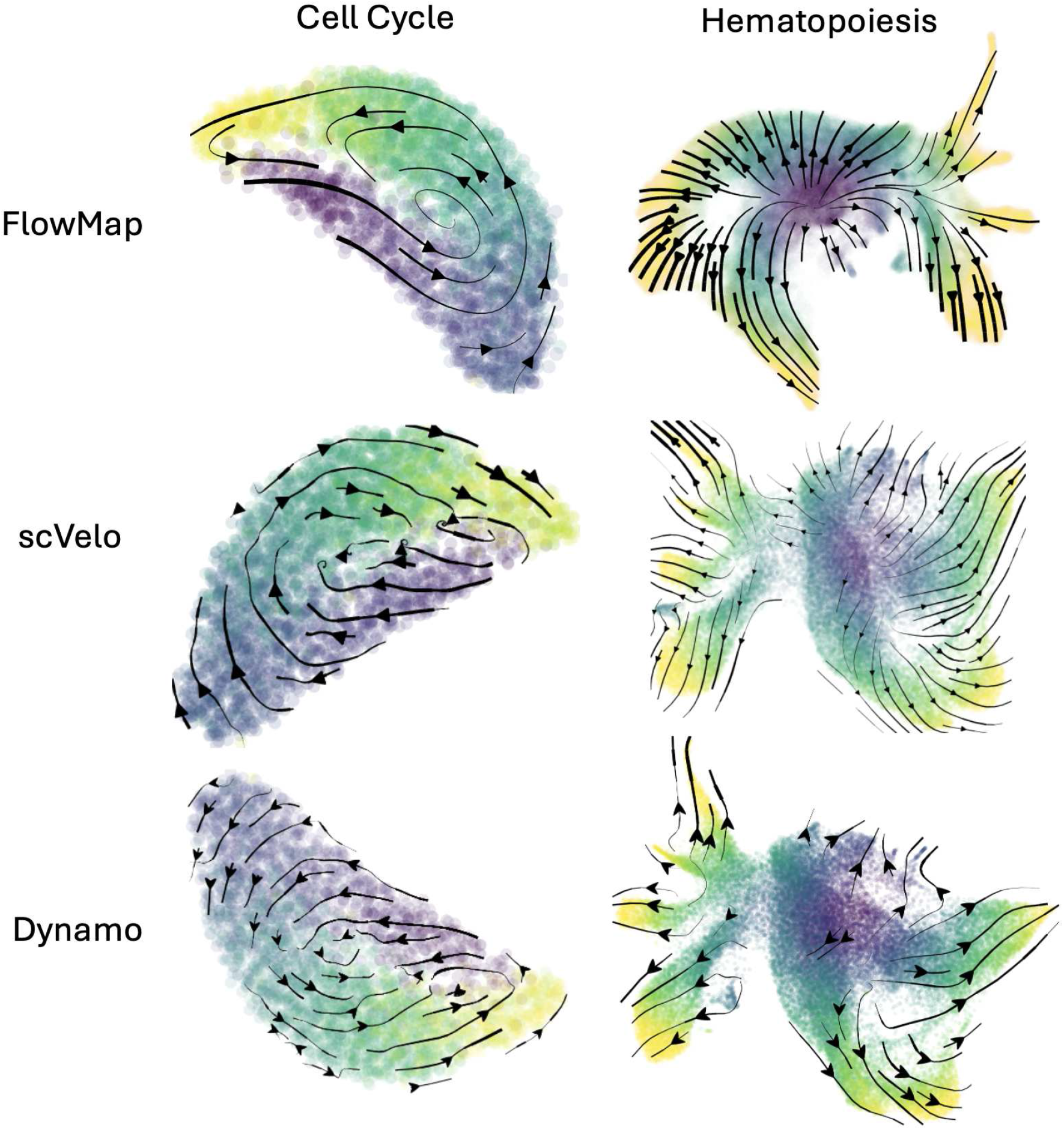
Benchmark of complete embedding and velocity analysis pipelines. Comparison of embeddings between FlowMap, scVelo and Dynamo on the cell-cycle and hematopoiesis datasets. The embedding is made with the default UMAP parameters.

**Figure S11:**
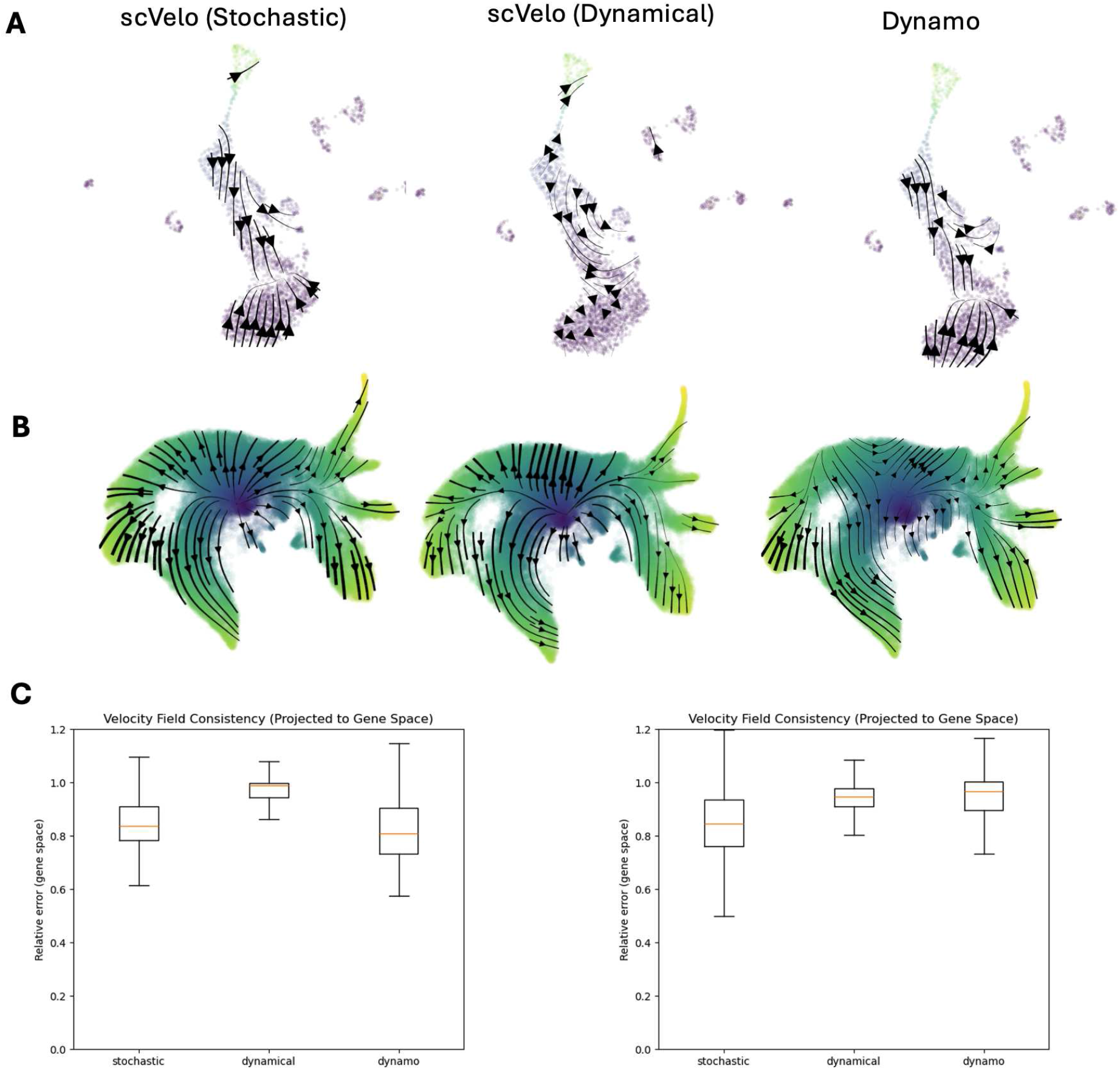
Comparison of velocity estimation methods. (A) Velocity vectors inferred by scVelo (stochastic), scVelo (dynamical), and Dynamo on the Dentate Gyrus dataset, visualized on a common embedding estimated using scVelo. (B) Velocity vectors inferred by scVelo (stochastic), scVelo (dynamical), and Dynamo on the Larry dataset, visualized on a common embedding estimated using FlowMap. (C) Quantitative evaluation of velocity field consistency after projecting the learned low-dimensional vector fields back to gene space. Box plots show cell-level relative errors between predicted and observed velocities for the Dentate Gyrus dataset (left) and Larry dataset (right).

**Figure S12:**
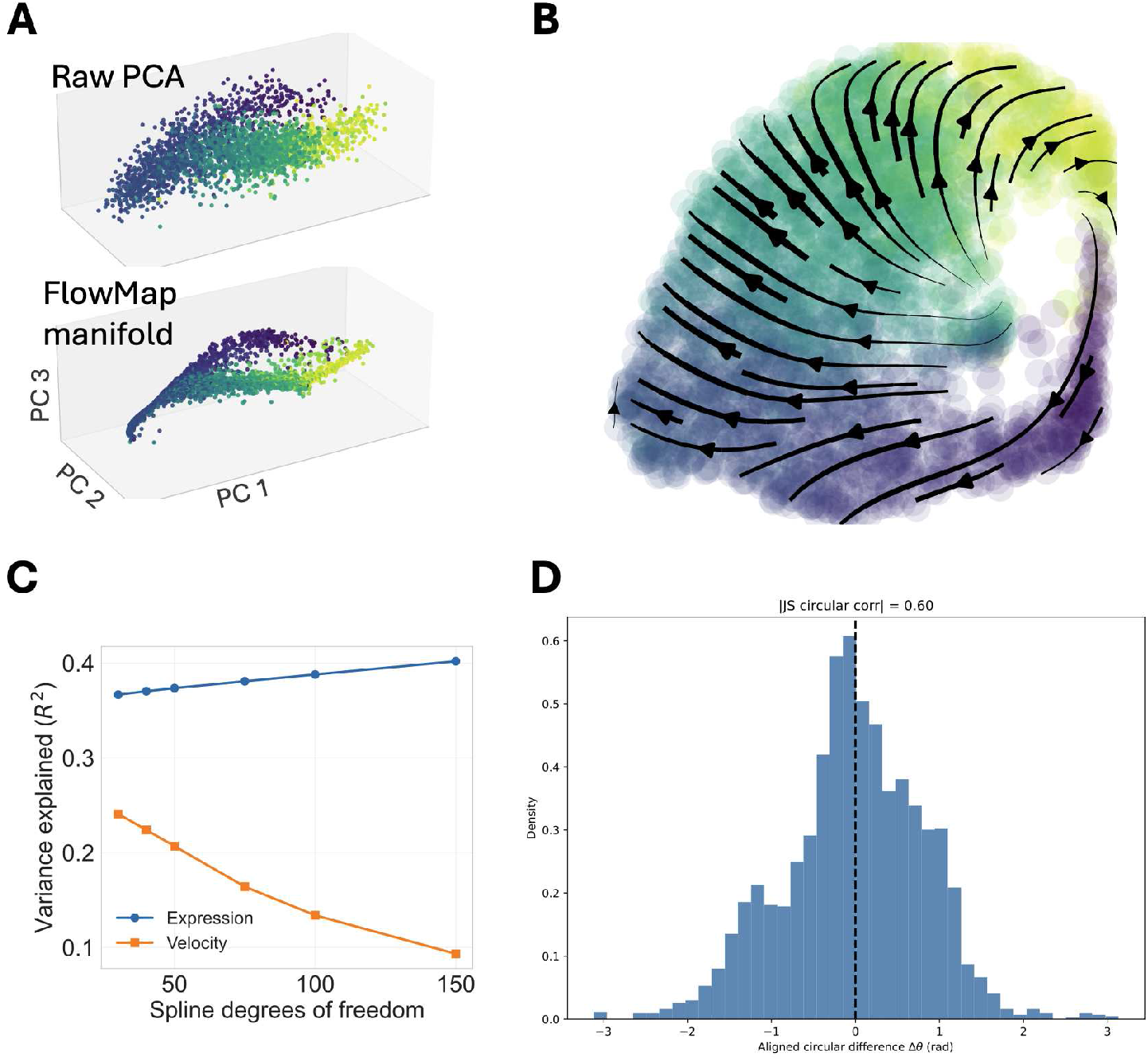
Diagnostics of FlowMap embedding using all highly variable genes. (A) FlowMap embedding and inferred RNA velocity field constructed using all 816 highly variable genes in the RPE1–FUCCI scEU-seq dataset. (B) Reconstruction performance as a function of spline complexity, showing variance explained (*R*^2^) for gene expression and RNA velocity across different degrees of freedom. (C) Evaluation of embedding geometry against ground-truth cell-cycle position using circular differences. Circular differences between embedding-derived angles and ground-truth phase are computed after correcting for a global rotational offset, and alignment is summarized using the Jammalamadaka–Sengupta circular–circular correlation coefficient.

**Figure S13:**
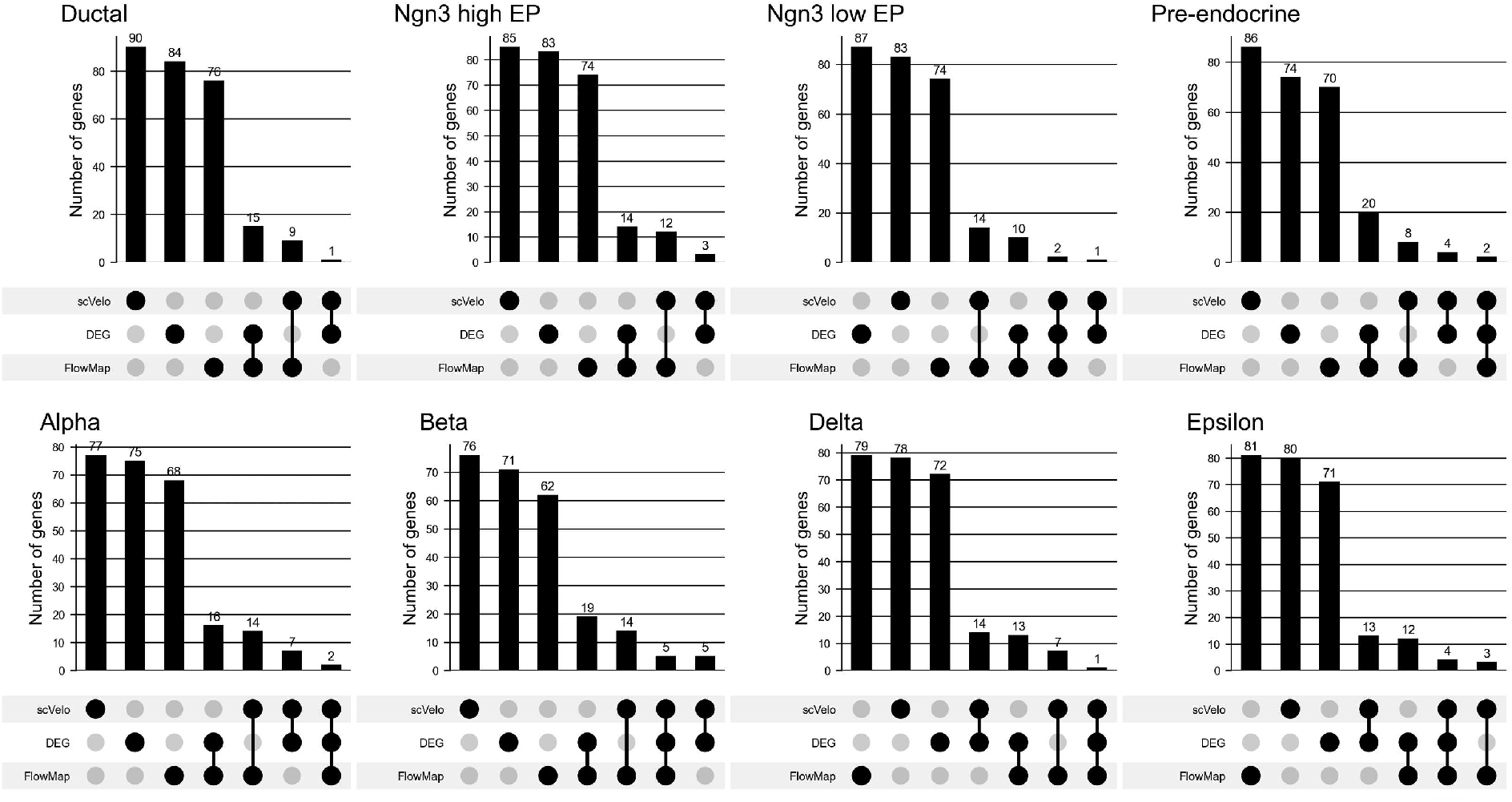
Overlap of prioritized genes across methods in pancreas endocrinogenesis. UpSet plots showing the overlap among the top 100 genes prioritized by FlowMap, differentially expressed genes (DEGs), and scVelo differential kinetics for each pancreatic cell state. Gene sets were computed independently within each cell type.

**Figure S14:**
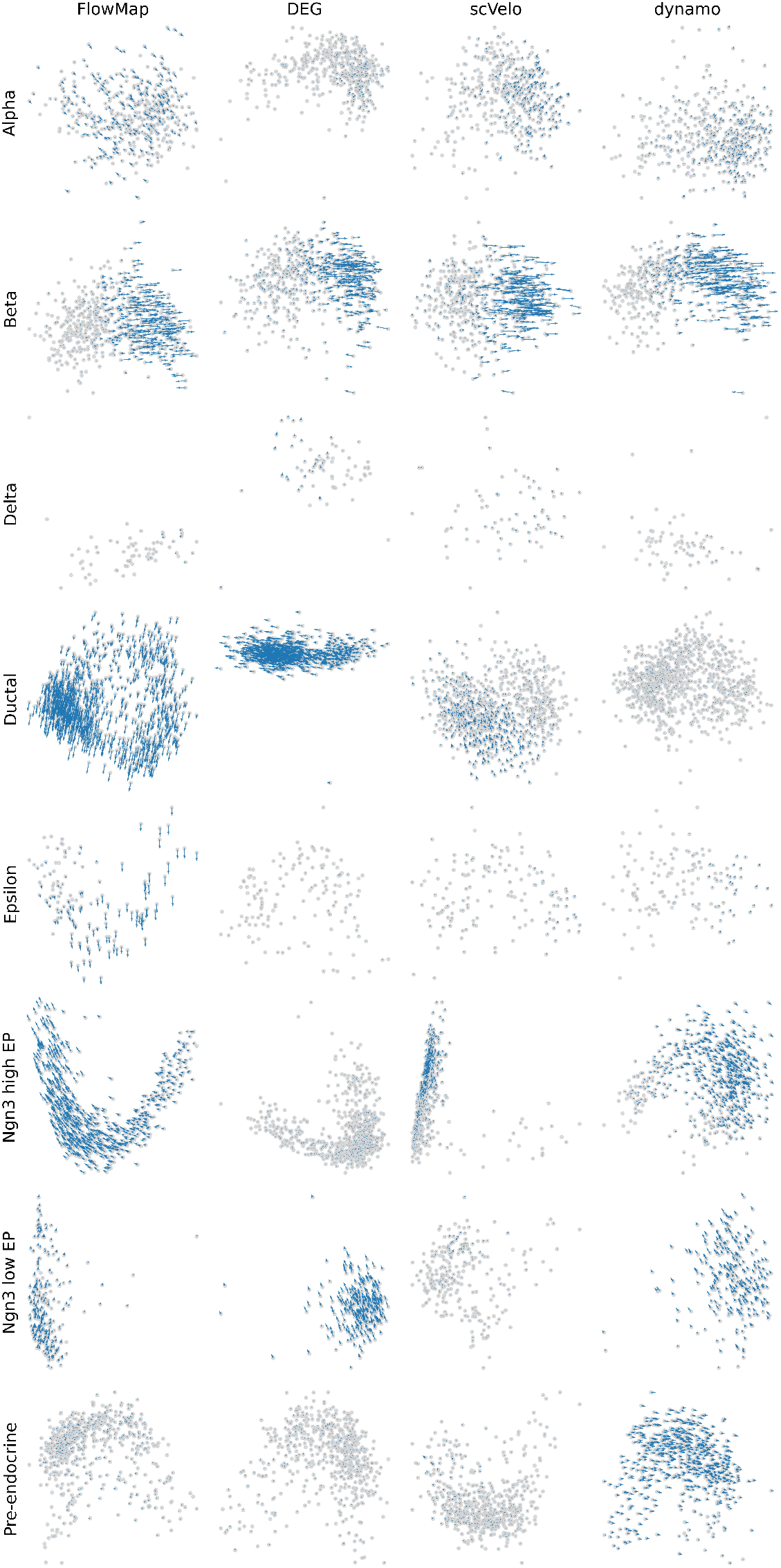
Local velocity structure induced by method-specific prioritized genes. Principal component projections of pancreatic cell states using the top 100 genes prioritized by each method within each cell type. For each panel, PCA was computed using only the selected genes from the corresponding method and cell type, and RNA velocity vectors were projected into the resulting two-dimensional PC space.

**Figure S15:**
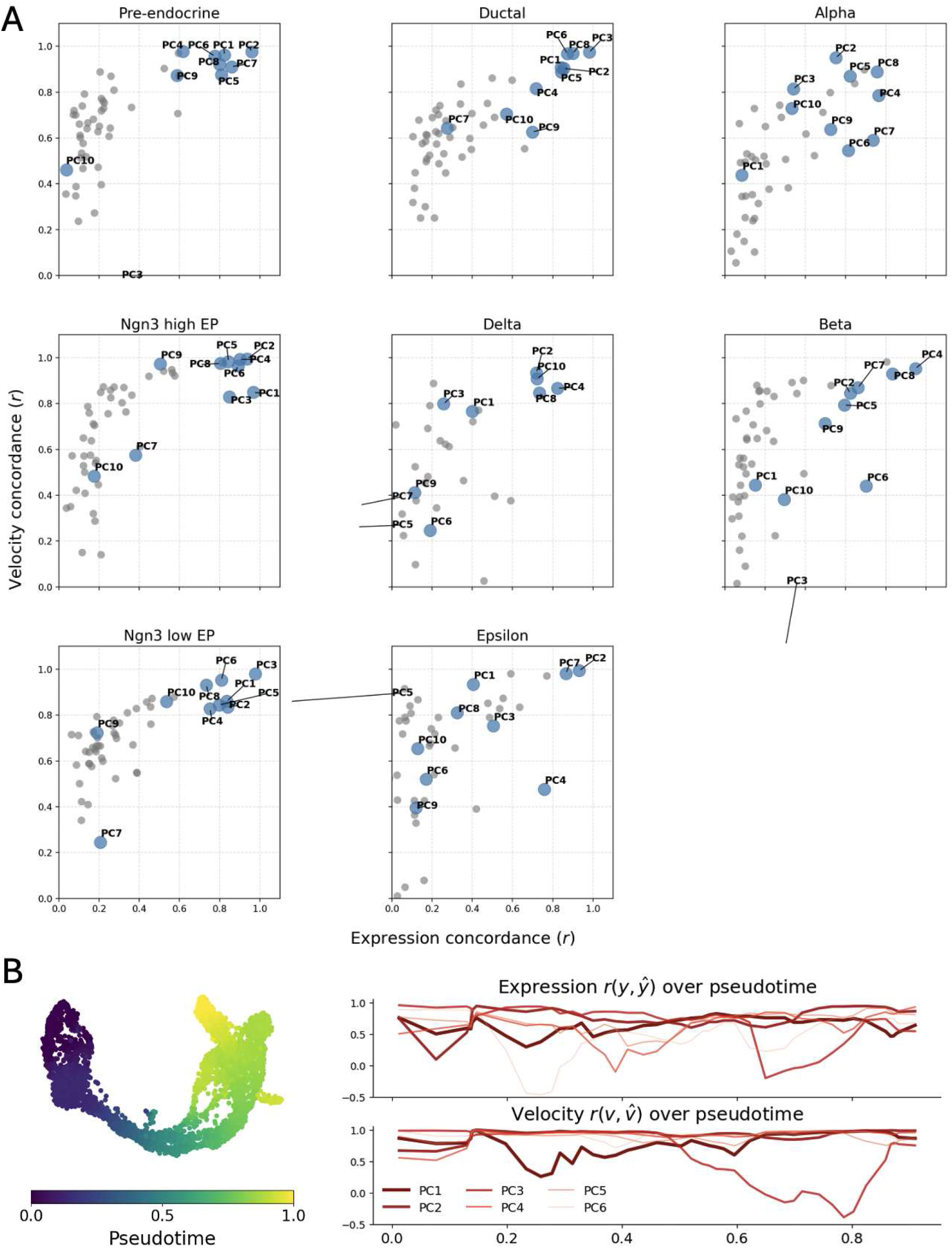
PC-level reconstruction concordance across pancreatic cell types. (A) For each cell type, expression and RNA velocity reconstruction consistencies are shown for PCs 1–50, with each point representing one principal component. PCs 1–10 are highlighted and labeled. (B) Expression and RNA velocity reconstruction consistency across pseudotime for PCs 1–6.

**Figure S16:**
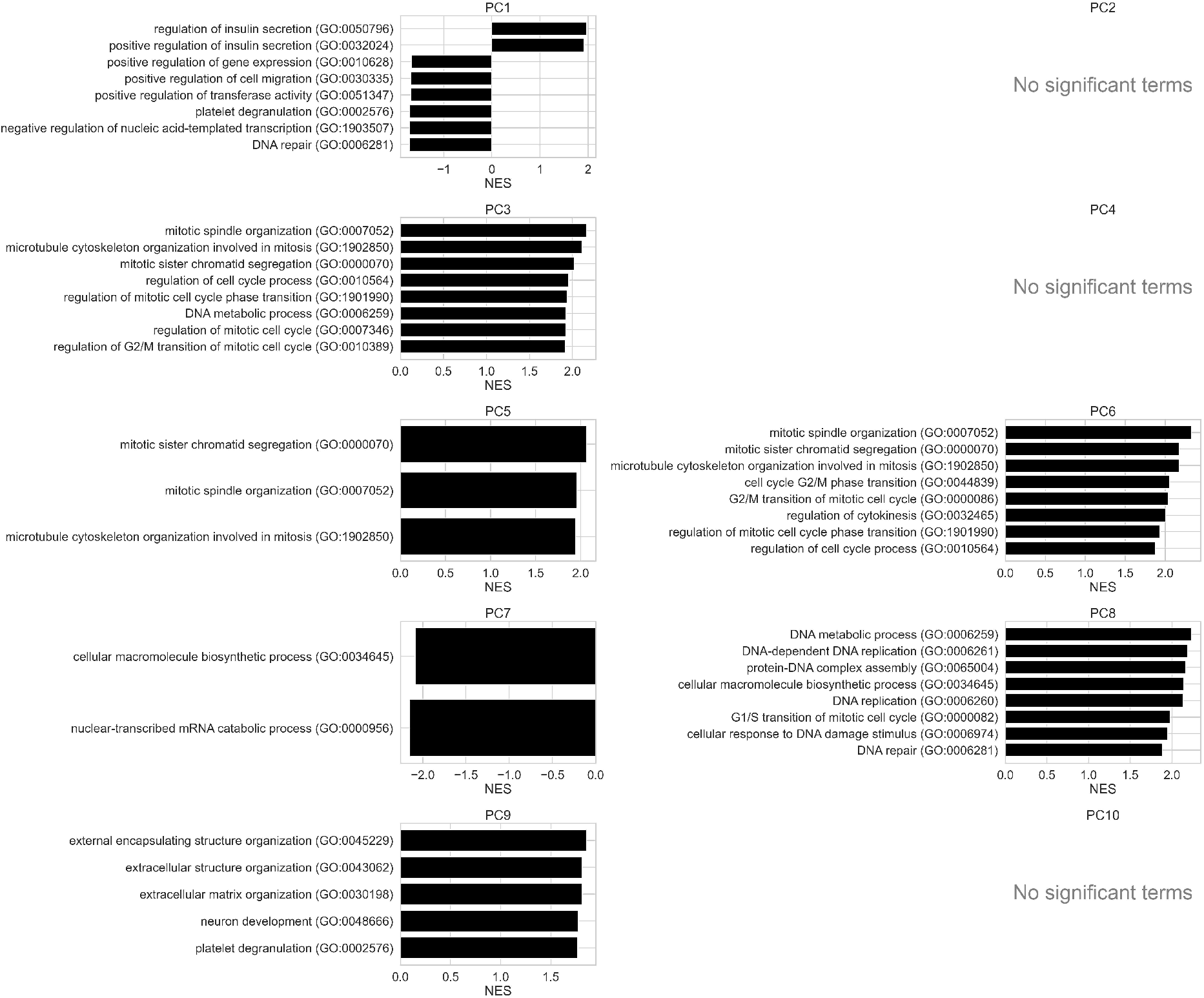
Gene set enrichment analysis of principal components in pancreatic differentiation. Preranked GSEA was performed on gene loadings for PCs 1–10 using GO Biological Process gene sets. Each panel shows the top enriched terms for a given PC when significant (FDR *<* 0.05); PCs without significant enrichment are indicated accordingly.

**Figure S17:**
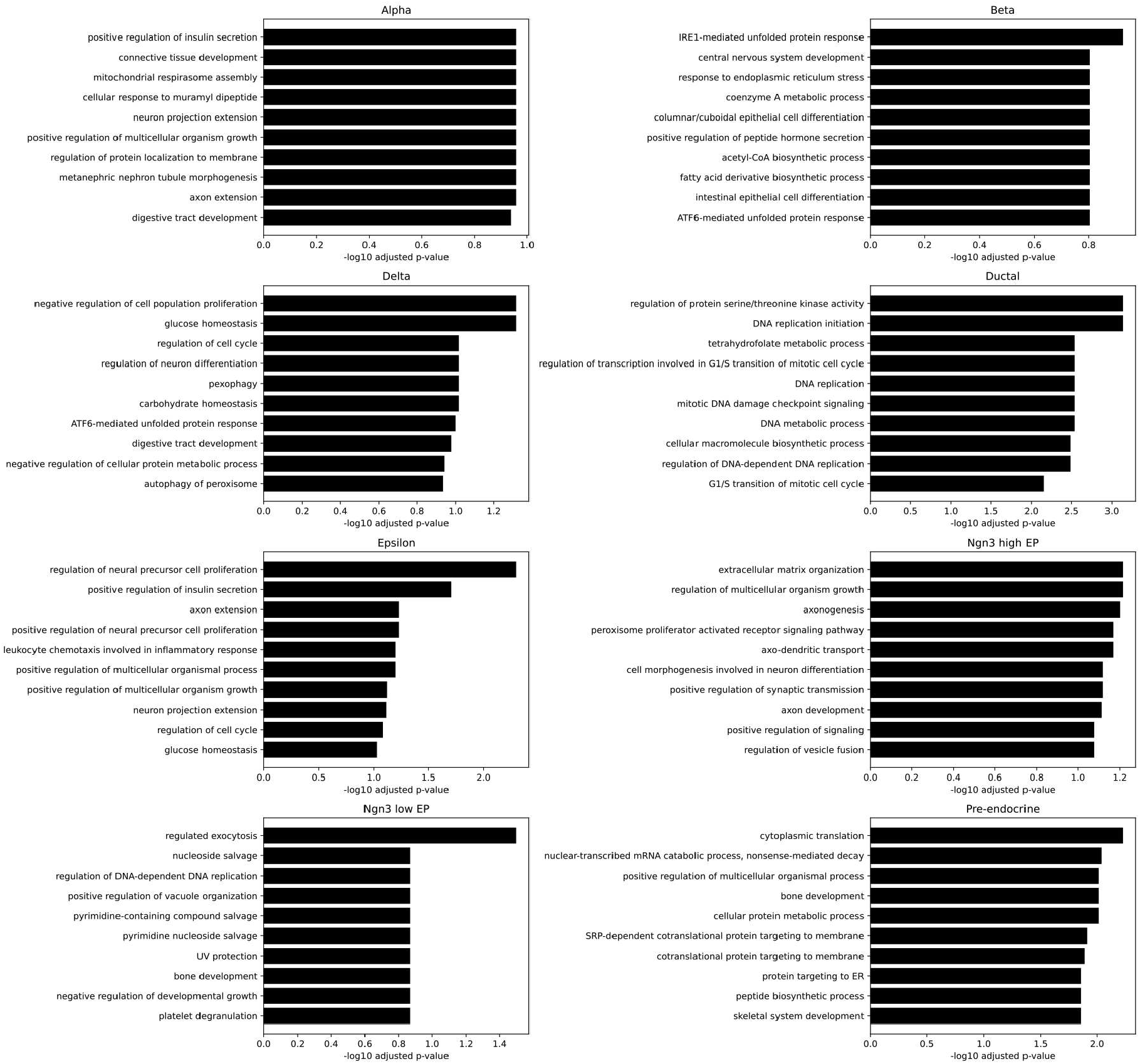
GO enrichment of FlowMap-prioritized pancreatic genes. Top GO biological process terms enriched among the top 100 FlowMap-prioritized genes for each pancreatic cell type.

**Figure S18:**
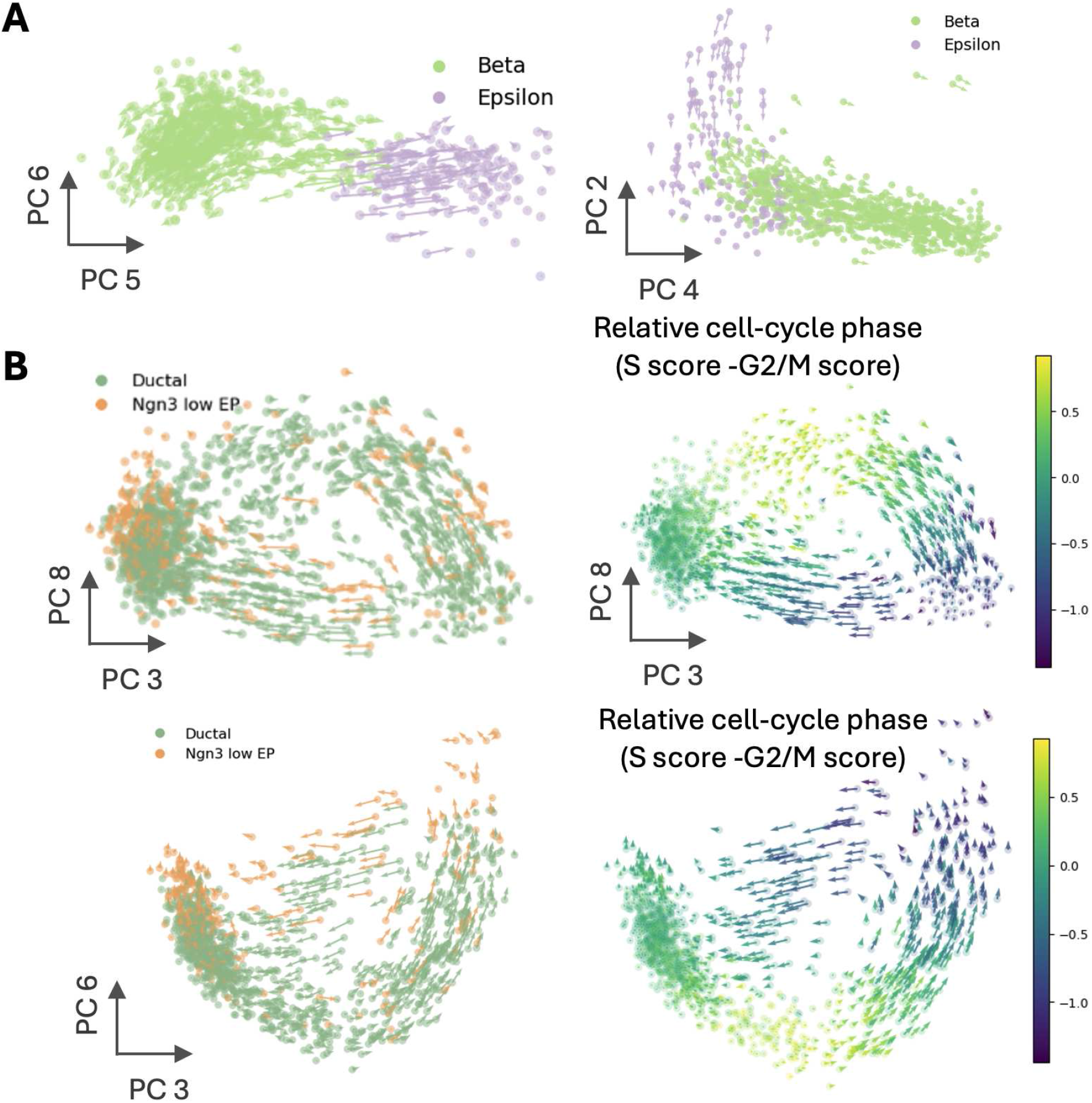
Additional vector field projections in pancreatic differentiation. Velocity fields projected onto additional principal component subspaces for pancreatic cell populations, highlighting cell-type–specific geometric structure across multiple latent dimensions.

**Figure S19:**
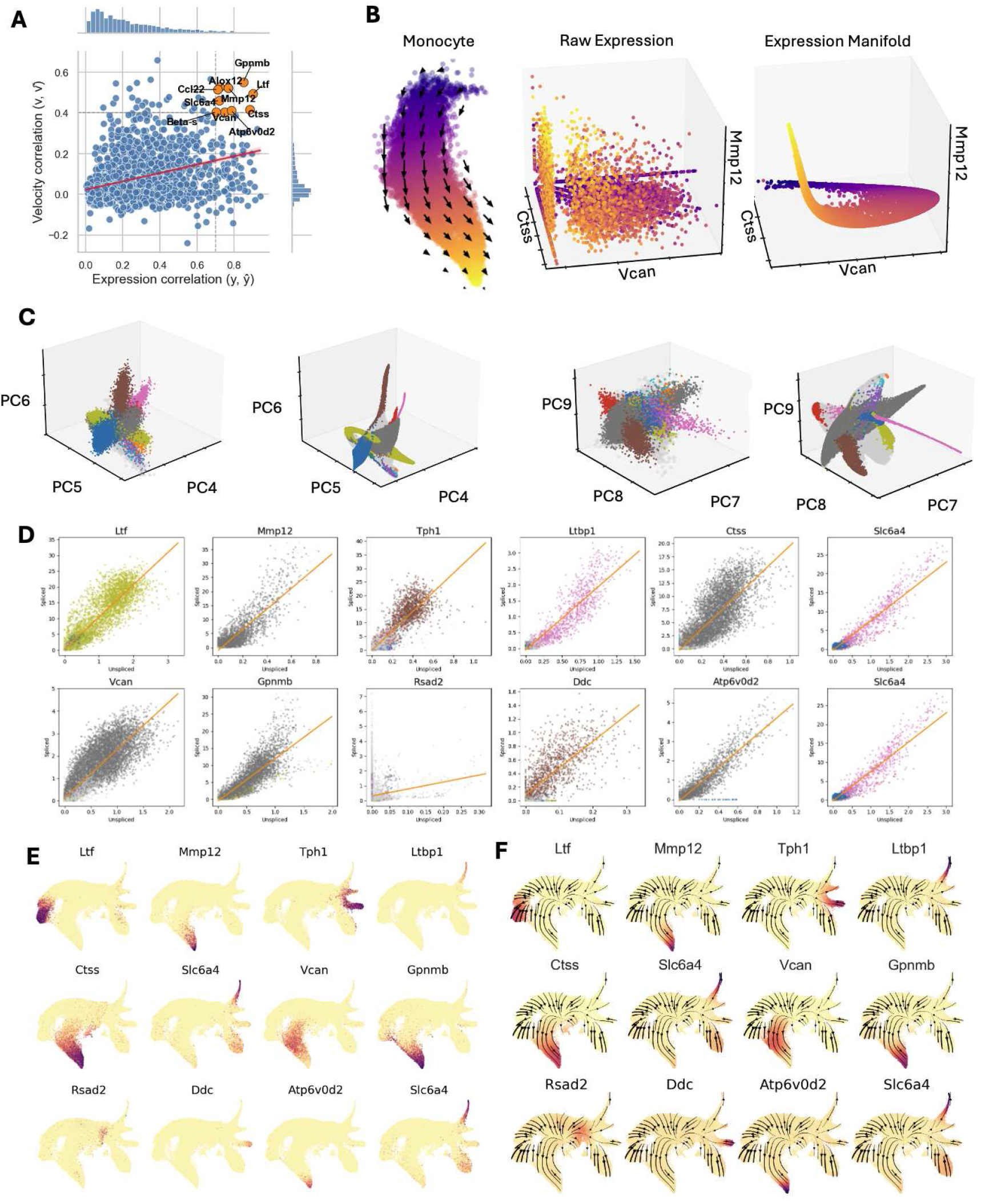
Gene-level velocity correlations and expression structure on the embedding. (**A**) Scatter plot showing gene-wise correlation between expression similarity and velocity similarity across cells, with selected genes highlighted. (**B**) Monocyte embedding with velocity vectors, alongside raw gene expression and the corresponding expression manifold for representative genes. (**C**) Principal component representations of gene expression, shown in raw PC space and after mapping to the learned manifold. (**D**) Phase portraits for selected genes, showing spliced versus unspliced transcript counts across cells with linear reference fits. (**E**) Raw gene expression projected onto the embedding for multiple genes. (**F**) Smoothed gene expression projected onto the embedding and visualized together with the learned velocity field, where the expression values can be viewed as a scalar field defined over the embedding.

**Figure S20:**
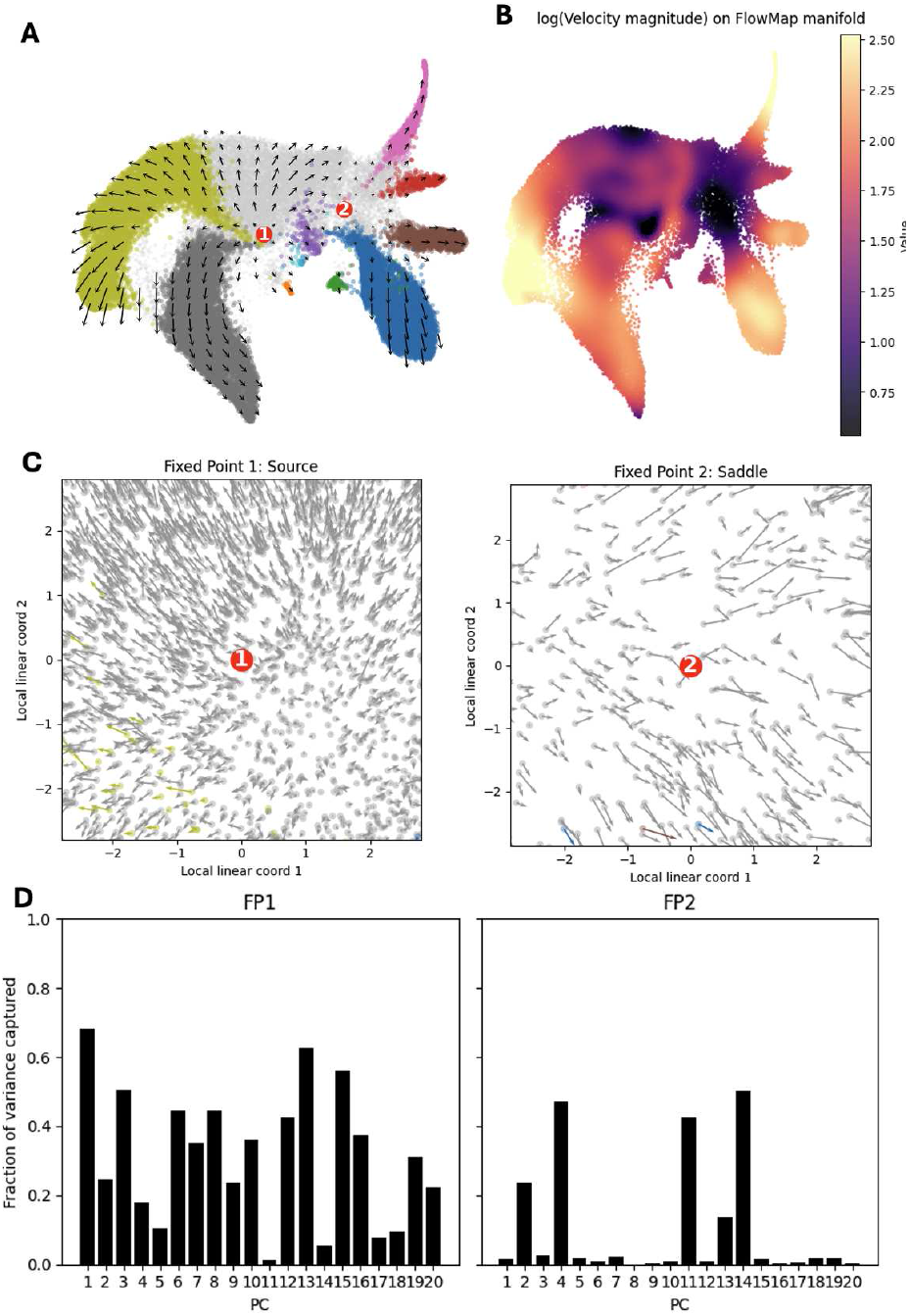
Fixed point structure and local linear analysis on the FlowMap embedding. (**A**) FlowMap embedding with the learned velocity field overlaid, highlighting two identified fixed points. (**B**) Visualization of the log-transformed velocity magnitude projected onto the FlowMap embedding. (**C**) Local linearized views of the velocity field around each fixed point, shown in locally flattened coordinates. (**D**) Alignment between principal components and the local linear subspace at each fixed point, quantified by the fraction of variance captured for the top principal components.

**Figure S21:**
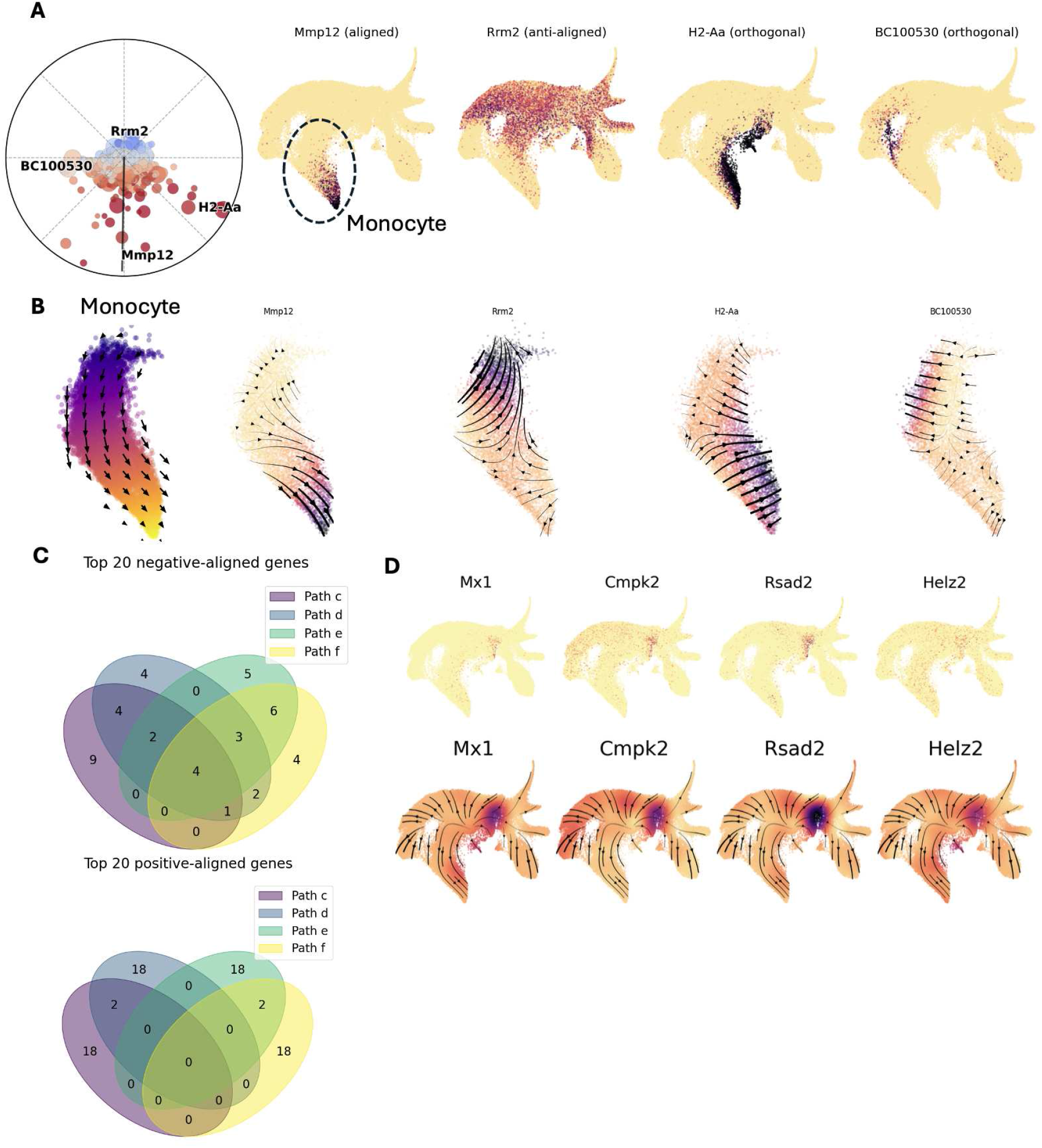
Gradient alignment and gene expression patterns in the monocyte trajectory. **(A)** Polar plot showing the alignment between gene-expression gradients and the inferred velocity field within the monocyte trajectory. Each point represents a gene, positioned by the angle between its gradient and the local velocity direction. One representative gene from each alignment category is highlighted, with corresponding smoothed expression patterns shown on the right. **(B)** Gradient vector fields of the same four genes restricted to monocyte cells. Streamlines indicate the direction of increasing gene expression. **(C)** Venn diagrams showing overlap among the top 20 negatively aligned genes (top) and positively aligned genes (bottom) across four inferred differentiation paths. **(D)** Smoothed expression of negatively aligned genes shared across all paths, visualized on the embedding.

**Figure S22:**
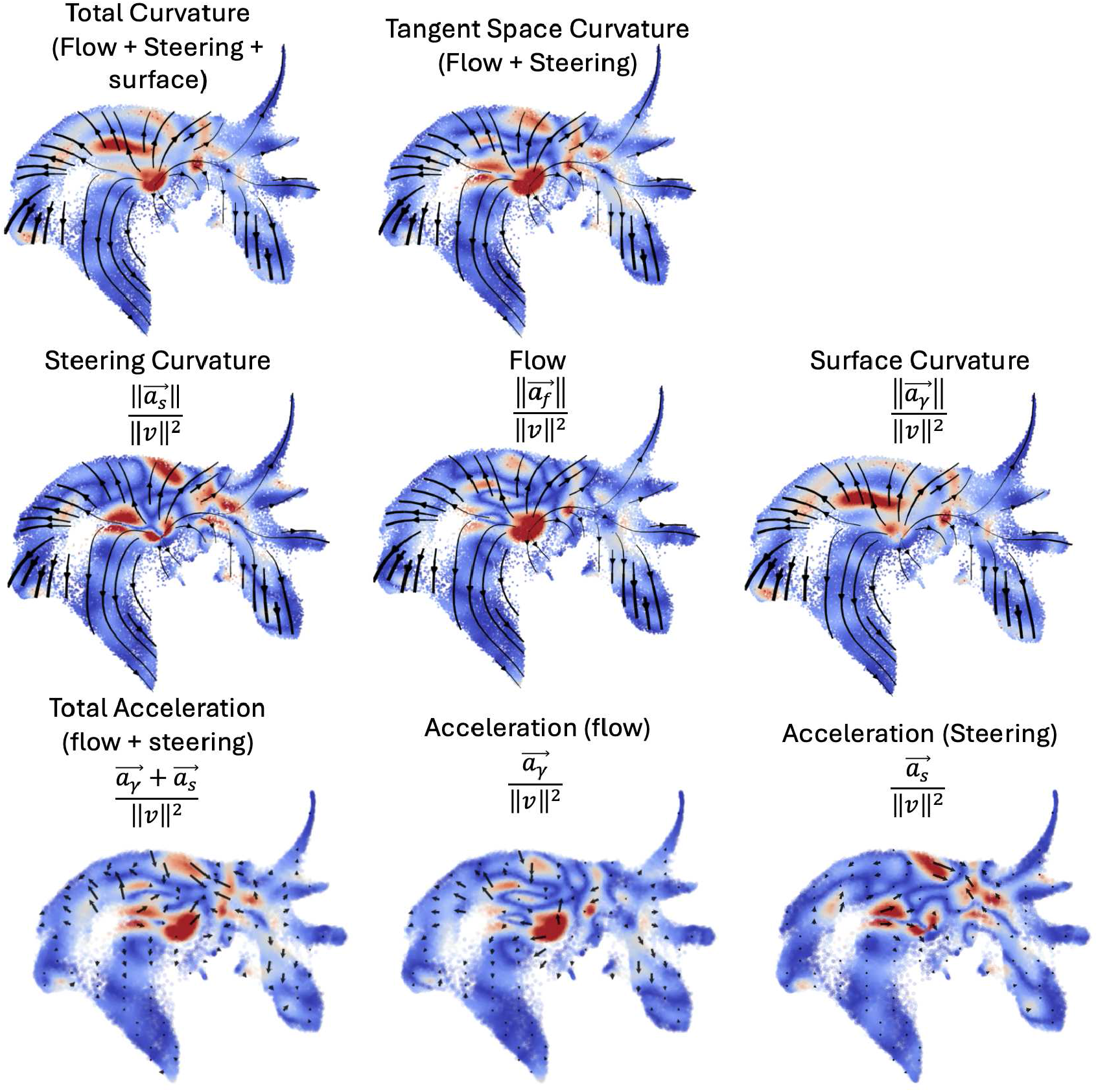
Additional visualization of curvature and acceleration components on the FlowMap embedding. Panels show the decomposition of RNA velocity acceleration into flow, steering, and surface components, together with the corresponding curvature quantities across the embedded developmental manifold. Streamlines indicate the inferred velocity field in the first two rows, while colors represent the magnitude of the corresponding acceleration or curvature term. In the third row, acceleration components are visualized directly as quiver vectors.

**Figure S23:**
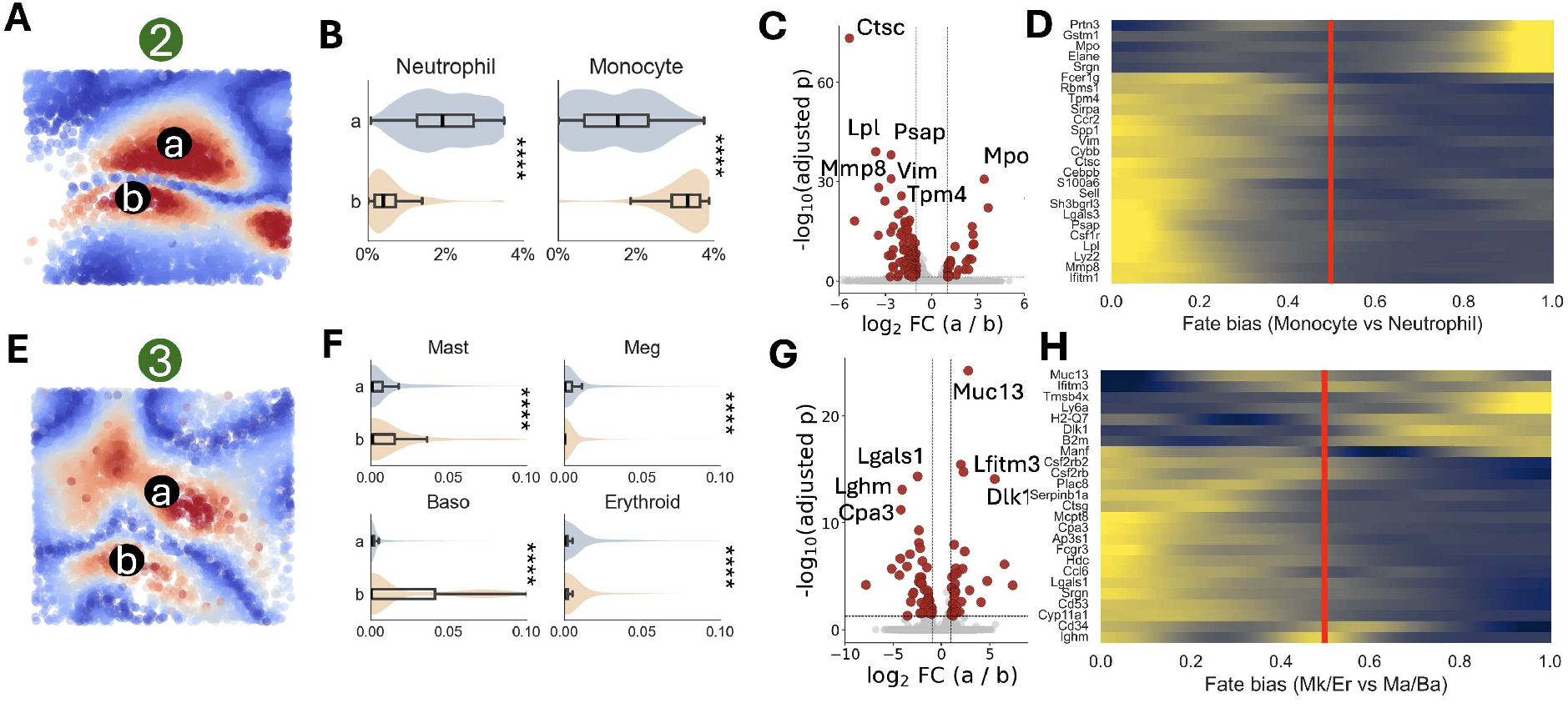
Additional steering-curvature regions associated with distinct fate biases and transcriptional programs. (A–D) Analysis of steering-curvature region 2. **(A)** FlowMap embedding showing the selected region and two subregions (a and b). **(B)** Comparison of inferred fate probabilities between subregions. **(C)** Differential expression analysis between subregions a and b. **(D)** Expression of differentially expressed genes ordered by relative fate bias within the selected region. **(E–H)** Same analysis as in (A–D) for steering-curvature region 3.

**Figure S24:**
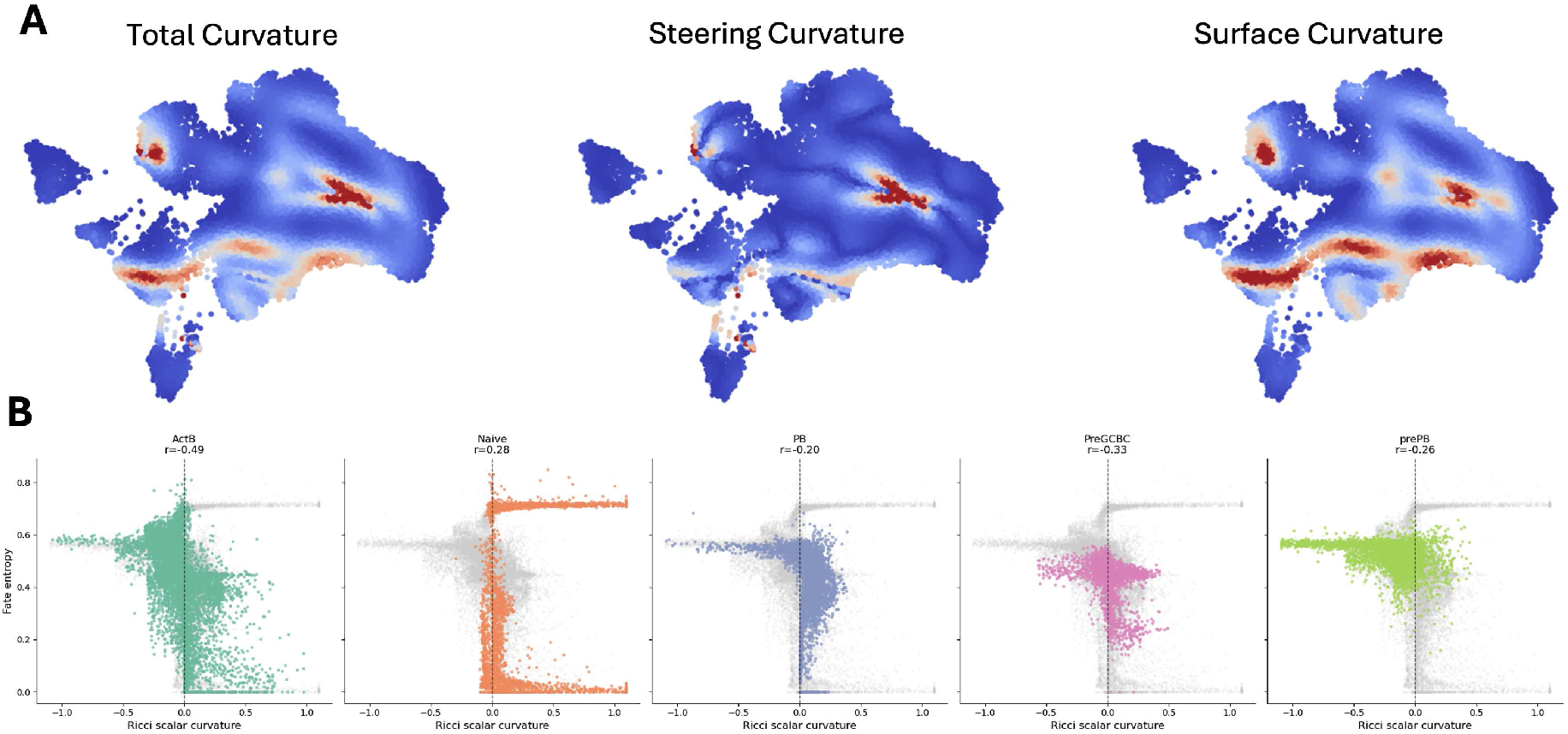
Curvature structure, gene expression patterns, and fate entropy across B-cell differentiation states. **(A)** Total curvature decomposed into steering and surface curvature components on the FlowMap embedding. **(B)** Scatter plots of Ricci scalar curvature versus CellRank fate entropy for individual B-cell populations. Pearson correlation coefficients are indicated above each panel.

**Figure S25:**
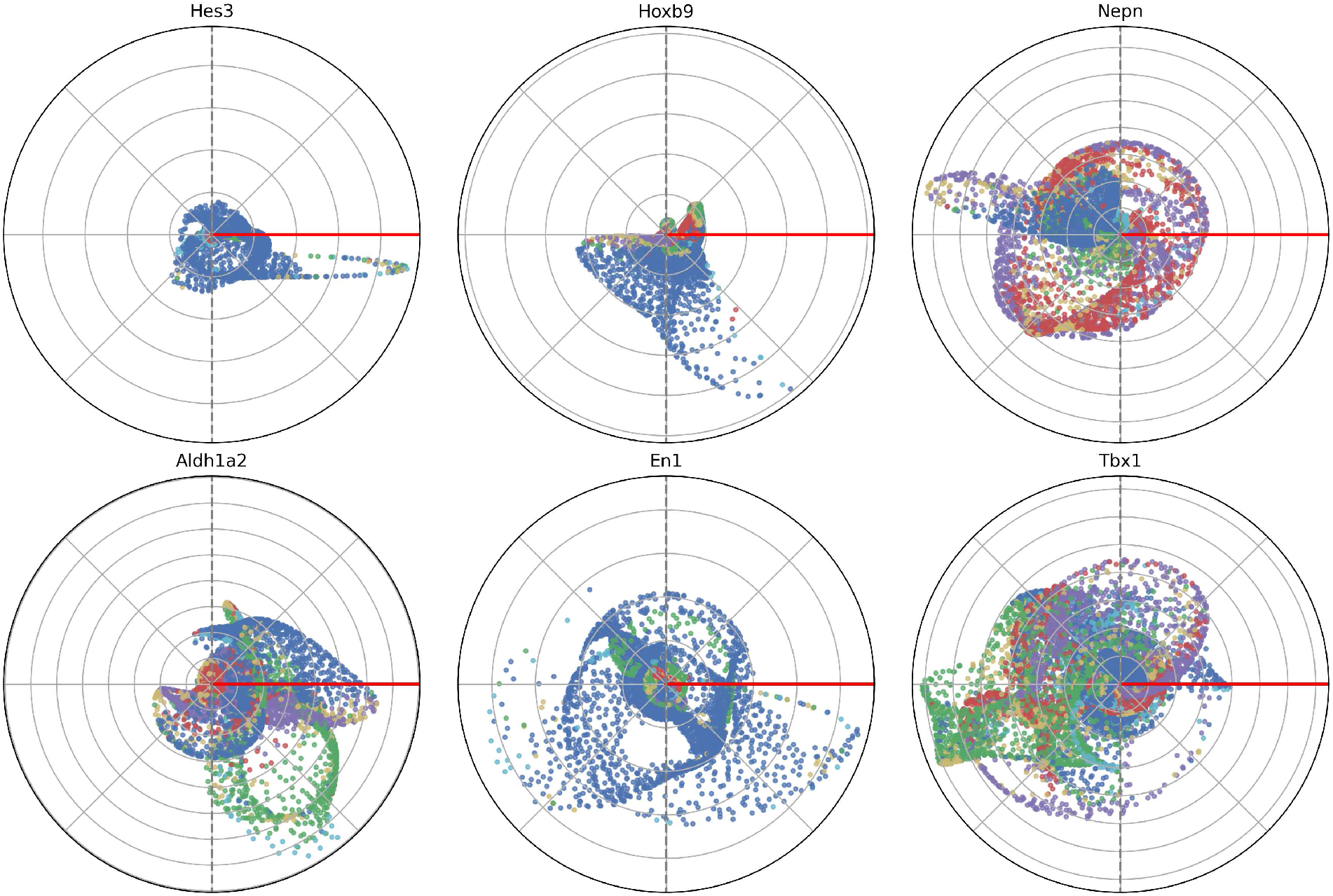
Gene-specific alignment of expression gradients with RNA velocity for mouse organogenesis data. Points show per-cell gradient direction (angle) and strength (radius) relative to RNA velocity, colored by cell type.

## References

[1] Cole Trapnell, Davide Cacchiarelli, Jonna Grimsby, Prapti Pokharel, Shuqiang Li, Michael Morse, Niall J Lennon, Kenneth J Livak, Tarjei S Mikkelsen, and John L Rinn. The dynamics and regulators of cell fate decisions are revealed by pseudotemporal ordering of single cells. Nature biotechnology, 32(4):381–386, 2014.

[2] Allon Wagner, Aviv Regev, and Nir Yosef. Revealing the vectors of cellular identity with single-cell genomics. Nature biotechnology, 34(11):1145–1160, 2016.

[3] Gioele La Manno, Ruslan Soldatov, Amit Zeisel, Emelie Braun, Hannah Hochgerner, Viktor Petukhov, Katja Lidschreiber, Maria E Kastriti, Peter Lönnerberg, Alessandro Furlan, et al. Rna velocity of single cells. Nature, 560(7719):494–498, 2018.

[4] Volker Bergen, Marius Lange, Stefan Peidli, F Alexander Wolf, and Fabian J Theis. Generalizing rna velocity to transient cell states through dynamical modeling. Nature biotechnology, 38(12):1408–1414, 2020.

[5] Xiaojie Qiu, Yan Zhang, Jorge D Martin-Rufino, Chen Weng, Shayan Hosseinzadeh, Dian Yang, Angela N Pogson, Marco Y Hein, Kyung Hoi Joseph Min, Li Wang, et al. Mapping transcriptomic vector fields of single cells. Cell, 185(4):690–711, 2022.

[6] Mingze Gao, Chen Qiao, and Yuanhua Huang. Unitvelo: temporally unified rna velocity reinforces single-cell trajectory inference. Nature Communications, 13(1):6586, 2022.

[7] Marius Lange, Volker Bergen, Michal Klein, Manu Setty, Bernhard Reuter, Mostafa Bakhti, Heiko Lickert, Meshal Ansari, Janine Schniering, Herbert B Schiller, et al. Cellrank for directed single-cell fate mapping. Nature methods, 19(2):159–170, 2022.

[8] Kevin R Moon, David Van Dijk, Zheng Wang, Scott Gigante, Daniel B Burkhardt, William S Chen, Kristina Yim, Antonia van den Elzen, Matthew J Hirn, Ronald R Coifman, et al. Visualizing structure and transitions in high-dimensional biological data. Nature biotechnology, 37(12):1482–1492, 2019.

[9] Dmitry Kobak and Philipp Berens. The art of using t-sne for single-cell transcriptomics. Nature communications, 10(1):5416, 2019.

[10] Caleb Weinreb, Samuel Wolock, Betsabeh K Tusi, Merav Socolovsky, and Allon M Klein. Fundamental limits on dynamic inference from single-cell snapshots. Proceedings of the National Academy of Sciences, 115(10):E2467–E2476, 2018.

[11] Meritxell Sáez, Robert Blassberg, Elena Camacho-Aguilar, Eric D Siggia, David A Rand, and James Briscoe. Statistically derived geometrical landscapes capture principles of decision-making dynamics during cell fate transitions. Cell systems, 13(1):12–28, 2022.

[12] Adam Gayoso, Philipp Weiler, Mohammad Lotfollahi, Dominik Klein, Justin Hong, Aaron Streets, Fabian J Theis, and Nir Yosef. Deep generative modeling of transcriptional dynamics for rna velocity analysis in single cells. Nature methods, 21(1):50–59, 2024.

[13] F Alexander Wolf, Fiona K Hamey, Mireya Plass, Jordi Solana, Joakim S Dahlin, Berthold Göttgens, Nikolaus Rajewsky, Lukas Simon, and Fabian J Theis. Paga: graph abstraction reconciles clustering with trajectory inference through a topology preserving map of single cells. Genome biology, 20(1):59, 2019.

[14] Yuhao Chen, Yan Zhang, Jiaqi Gan, Ke Ni, Ming Chen, Ivet Bahar, and Jianhua Xing. Graphvelo allows for accurate inference of multimodal velocities and molecular mechanisms for single cells. Nature Communications, 16(1):7831, 2025.

[15] Haotian Cui, Hassaan Maan, Maria C Vladoiu, Jiao Zhang, Michael D Taylor, and Bo Wang. Deepvelo: deep learning extends rna velocity to multi-lineage systems with cell-specific kinetics. Genome biology, 25(1):27, 2024.

[16] Chen Qiao and Yuanhua Huang. Representation learning of rna velocity reveals robust cell transitions. Proceedings of the National Academy of Sciences, 118(49):e2105859118, 2021.

[17] Joshua B Tenenbaum, Vin de Silva, and John C Langford. A global geometric framework for nonlinear dimensionality reduction. science, 290(5500):2319–2323, 2000.

[18] Sam T Roweis and Lawrence K Saul. Nonlinear dimensionality reduction by locally linear embedding. science, 290(5500):2323–2326, 2000.

[19] Laurens van der Maaten and Geoffrey Hinton. Visualizing data using t-sne. Journal of machine learning research, 9(Nov):2579–2605, 2008.

[20] Leland McInnes, John Healy, and James Melville. Umap: Uniform manifold approximation and projection for dimension reduction. arXiv preprint arXiv:1802.03426, 2018.

[21] Tara Chari and Lior Pachter. The specious art of single-cell genomics. PLOS Computational Biology, 19(8):e1011288, 2023.

[22] Jean Duchon. Splines minimizing rotation-invariant semi-norms in sobolev spaces. In Constructive theory of functions of several variables: proceedings of a conference held at oberwolfach April 25–May 1, 1976, pages 85–100. Springer, 2006.

[23] Trevor Hastie and Werner Stuetzle. Principal curves. Journal of the American statistical association, 84(406):502–516, 1989.

[24] Lyla Atta, Arpan Sahoo, and Jean Fan. Veloviz: Rna velocity-informed embeddings for visualizing cellular trajectories. Bioinformatics, 38(2):391–396, 2022.

[25] Nico Battich, Joep Beumer, Buys De Barbanson, Lenno Krenning, Chloé S Baron, Marvin E Tanenbaum, Hans Clevers, and Alexander Van Oudenaarden. Sequencing metabolically labeled transcripts in single cells reveals mrna turnover strategies. Science, 367(6482):1151–1156, 2020.

[26] Sonia Nestorowa, Fiona K Hamey, Blanca Pijuan Sala, Evangelia Diamanti, Mairi Shepherd, Elisa Laurenti, Nicola K Wilson, David G Kent, and Berthold Göttgens. A single-cell resolution map of mouse hematopoietic stem and progenitor cell differentiation. Blood, The Journal of the American Society of Hematology, 128(8):e20–e31, 2016.

[27] Aimée Bastidas-Ponce, Sophie Tritschler, Leander Dony, Katharina Scheibner, Marta Tarquis-Medina, Ciro Salinno, Silvia Schirge, Ingo Burtscher, Anika Böttcher, Fabian J Theis, et al. Comprehensive single cell mrna profiling reveals a detailed roadmap for pancreatic endocrinogenesis. Development, 146(12):dev173849, 2019.

[28] Caleb Weinreb, Alejo Rodriguez-Fraticelli, Fernando D Camargo, and Allon M Klein. Lineage tracing on transcriptional landscapes links state to fate during differentiation. Science, 367(6479):eaaw3381, 2020.

[29] Duluxan Sritharan, Shu Wang, and Sahand Hormoz. Computing the riemannian curvature of image patch and single-cell rna sequencing data manifolds using extrinsic differential geometry. Proceedings of the national academy of sciences, 118(29):e2100473118, 2021.

[30] Tram Huynh and Zixuan Cang. Topological and geometric analysis of cell states in single-cell transcriptomic data. Briefings in Bioinformatics, 25(3):bbae176, 2024.

[31] Manfredo P Do Carmo. Differential geometry of curves and surfaces: revised and updated second edition. Courier Dover Publications, 2016.

[32] Anthony Baptista, Ben D MacArthur, and Christopher RS Banerji. Charting cellular differentiation trajectories with ricci flow. Nature Communications, 15(1):2258, 2024.

[33] Nicholas A Pease, Jingyu Fan, Swapnil Keshari, Jered Stratton, Peter Gerges, Betsy Ann Varghese, Narayanan VP Nampoothiri, Christopher S McGinnis, Wenxi Zhang, Steven B Gierlack, et al. Cell cycle-coupled transcriptional network orchestrates human b cell fate bifurcation. bioRxiv, 2025.

[34] Lewis Wolpert. Positional information and patterning revisited. Journal of Theoretical Biology, 269(1):359–365, 2011.

[35] An Wang, Donald Geman, Uthsav Chitra, and Laurent Younes. Mapping spatial gradients in spatial transcriptomics data with score matching. bioRxiv, pages 2025–11, 2025.

[36] Tim Lohoff, Shila Ghazanfar, A Missarova, Noushin Koulena, Nico Pierson, Jonathan A Griffiths, Evan S Bardot, C-HL Eng, Richard CV Tyser, Ricard Argelaguet, et al. Integration of spatial and single-cell transcriptomic data elucidates mouse organogenesis. Nature biotechnology, 40(1):74–85, 2022.

[37] Tamim Abdelaal, Laurens M Grossouw, R Jeroen Pasterkamp, Boudewijn PF Lelieveldt, Marcel JT Reinders, and Ahmed Mahfouz. Sirv: spatial inference of rna velocity at the single-cell resolution. NAR genomics and bioinformatics, 6(3):lqae100, 2024.

[38] Muriel Rhinn and Pascal Dollé. Retinoic acid signalling during development. Development, 139(5):843–858, 2012.

[39] Qi Qiu, Peng Hu, Xiaojie Qiu, Kiya W Govek, Pablo G Cámara, and Hao Wu. Massively parallel and time-resolved rna sequencing in single cells with scnt-seq. Nature methods, 17(10):991–1001, 2020.

[40] Chen Li, Maria C Virgilio, Kathleen L Collins, and Joshua D Welch. Multi-omic single-cell velocity models epigenome–transcriptome interactions and improves cell fate prediction. Nature biotechnology, 41(3):387–398, 2023.

[41] Kenji Kamimoto, Christian M. Hoffmann, and Samantha A. Morris. Dissecting cell identity via network inference and in silico gene perturbation. Nature, 614(7949):742–751, 2023.

[42] Jiachen Li, Xiaoyong Pan, Hong-Bin Shen, and Yuan Ye. Tfvelo: gene regulation inspired rna velocity estimation. Nature Communications, 15(1):1740, 2024.

[43] Weixu Wang, Zhiyuan Hu, Philipp Weiler, Sarah Mayes, Marius Lange, Daniel M. Fountain, Julianna O. Haug, Jingye Wang, Zhengyuan Xue, Tatjana Sauka-Spengler, and Fabian J. Theis. Regvelo: Gene-regulatory-informed dynamics of single cells. Cell, 189:1–28, 2026.

[44] George S Kimeldorf and Grace Wahba. A correspondence between bayesian estimation on stochastic processes and smoothing by splines. The Annals of Mathematical Statistics, 41(2):495–502, 1970.

[45] Grace Wahba. Spline models for observational data. SIAM, 1990.

[46] Shou-Wen Wang, Michael J Herriges, Kilian Hurley, Darrell N Kotton, and Allon M Klein. Cospar identifies early cell fate biases from single-cell transcriptomic and lineage information. Nature Biotechnology, 40(7):1066–1074, 2022.

[47] Jarkko Venna and Samuel Kaski. Neighborhood preservation in nonlinear projection methods: An experimental study. In International conference on artificial neural networks, pages 485–491. Springer, 2001.

[48] Neo Christopher Chung, BłaŽej Miasojedow, Michał Startek, and Anna Gambin. Jaccard/tanimoto similarity test and estimation methods for biological presence-absence data. BMC bioinformatics, 20(Suppl 15):644, 2019.

