## Supplemental Note 1 for "FlowMap: Geometry–Dynamics Consistent Embedding of RNA Velocity for Interpretable Cellular Trajectories"

#### 1 Velocity-aware phase distance metric

##### 1.1 Definition of phase

Given a vector field, cells sharing the same *phase* are defined as those lying on the same *level set* of an associated scalar function. These level sets are locally orthogonal to the direction of the vector field and represent points that are synchronized along the flow.

Formally, consider a vector field $V(x)$ defined on a smooth manifold, governing the dynamics of particles via

$$\frac{dx}{d\tau} = V(x),$$

where $\tau$ denotes a *time/phase parameter* along the flow.

In the special case of a potential vector field in Euclidean space, the vector field is the gradient of a scalar potential function $\Phi$, i.e., $V(x) = \nabla\Phi(x)$. In this setting, the phase of a point $x$ is naturally defined as $\Phi(x)$, and points of equal phase lie on the level sets

$$\{x : \Phi(x) = c, \text{ for constant } c\}.$$

More generally, for an arbitrary vector field $V(x)$, the phase difference between two points $x_1$ and $x_2$ along a flow curve can be expressed as

$$\tau_2 - \tau_1 = \int_{x_1}^{x_2} \frac{V(x)^\top}{\|V(x)\|^2}\, d\ell,$$

where $d\ell$ denotes the differential arc-length along a smooth path connecting $x_1$ to $x_2$. This expression defines a local notion of phase difference that is consistent with the flow direction, and remains valid for vector fields defined on smooth manifolds.

##### 1.2 Interpretation of phase distance

The phase distance between two nearby points can be interpreted as a local estimate of their relative progression along the vector field. In practice, this quantity is inferred from the displacement between the points projected onto their respective velocity directions.

To build intuition, consider the symmetric first-order estimator

$$\hat{t}_{\text{sym}} = \frac{1}{2}\left[\frac{(x_2 - x_1)^\top V(x_1)}{\|V(x_1)\|^2} + \frac{(x_2 - x_1)^\top V(x_2)}{\|V(x_2)\|^2}\right],$$

which provides a local approximation of the phase difference between two points $x_1$ and $x_2$.

The first term corresponds to a forward estimate of the phase difference obtained by projecting the displacement $x_2 - x_1$ onto the velocity at $x_1$, i.e., solving the linearized relation

$$x_2 \approx x_1 + t_1 V(x_1),$$

while the second term corresponds to a backward estimate obtained by projecting onto $V(x_2)$, i.e.,

$$x_1 \approx x_2 + t_2 V(x_2).$$

The symmetric estimator $\hat{t}_{\text{sym}}$ is thus the average of these forward and backward first-order approximations, reducing directional bias and improving robustness.

From a continuous perspective, this estimator can be interpreted as a discrete approximation of the phase difference integral

$$\tau_2 - \tau_1 = \int_{x_1}^{x_2} \frac{V(x)^\top}{\|V(x)\|^2} \, d\ell,$$

where $\hat{t}_{\text{sym}}$ corresponds to a trapezoidal-rule approximation using local evaluations at the endpoints.

The full regression-based estimator introduced in the main text generalizes this first-order approximation by jointly using both positions and velocities to infer a consistent pair of local phase parameters. In this sense, $\hat{t}_{\text{sym}}$ provides an interpretable linearization, while the full estimator yields a more accurate and stable estimate of the phase distance in practice.

##### 1.3 Computing phase distance via local alignment

To compute the phase distance between two neighboring cells $i$ and $j$, we introduce a local alignment framework based on RNA velocity. Let the observed gene expression profiles be $x_i, x_j \in \mathbb{R}^p$ with corresponding velocity vectors $v_i, v_j \in \mathbb{R}^p$, where $j \in \mathcal{N}(i)$ denotes that cell $j$ is a neighbor of cell $i$.

We define the alignment distance after allowing displacements along the velocity directions as

$$\left\|(x_i + \tau_{ij} v_i) - (x_j + \tau'_{ij} v_j)\right\|,$$

where $\tau_{ij}$ and $\tau'_{ij}$ are local phase parameters.

The *phase distance* is then defined as the difference in optimal phase parameters:

$$D_{ij} = \left|\tau^*_{ij} - \tau'^*_{ij}\right|, \quad (\tau^*_{ij}, \tau'^*_{ij}) = \arg\min_{\tau_{ij}, \tau'_{ij}} \left\|(x_i + \tau_{ij} v_i) - (x_j + \tau'_{ij} v_j)\right\|^2.$$

Intuitively, this formulation measures how much each cell must progress along its local velocity direction in order to best align with its neighbor. The resulting phase distance therefore captures relative progression along the flow, rather than orthogonal variation in expression space.

The minimization problem admits a closed-form solution by solving a $2 \times 2$ linear system for each neighboring pair:

$$\min_{\boldsymbol{\tau}_{ij}} \left\|(x_i - x_j) + v_i \tau_{ij} - v_j \tau'_{ij}\right\|_2^2, \quad \boldsymbol{\tau}_{ij} = \begin{bmatrix} \tau_{ij} \\ \tau'_{ij} \end{bmatrix}.$$

Equivalently, this can be written in normal equation form as

$$\mathbf{Q}_{ij} \boldsymbol{\tau}_{ij} = \mathbf{b}_{ij},$$

where

$$\mathbf{Q}_{ij} = \begin{bmatrix} v_i^\top v_i & -v_i^\top v_j \\ -v_i^\top v_j & v_j^\top v_j \end{bmatrix}, \quad \mathbf{b}_{ij} = \begin{bmatrix} v_i^\top (x_j - x_i) \\ v_j^\top (x_i - x_j) \end{bmatrix}.$$

This formulation allows efficient computation of $\tau^*_{ij}$ and $\tau'^*_{ij}$, and thus the phase distance $D_{ij}$, for all neighboring pairs of cells.

#### 2 Spline-based manifold reconstruction

We construct a smooth mapping from the low-dimensional embedding space to the high-dimensional expression space. Let $\mathbf{y} \in \mathbb{R}^d$ denote embedding coordinates and $\mathbf{x} \in \mathbb{R}^p$ denote the corresponding expression profiles (in practice, principal component coordinates). The goal is to estimate a differentiable map

$$\psi : \mathbb{R}^d \to \mathbb{R}^p, \quad \psi(\mathbf{y}_i) \approx \mathbf{x}_i,$$

that reconstructs the high-dimensional structure from the embedding.

##### 2.1 Spline formulation

We model $\psi$ as a vector-valued spline function of the form

$$\psi(\mathbf{y}) = \sum_{i=1}^{k} \mathbf{c}_i \, K(\mathbf{y}, \mathbf{y}_i) + \mathbf{A}\mathbf{y} + \mathbf{b}, \tag{2.1}$$

where $K : \mathbb{R}^d \times \mathbb{R}^d \to \mathbb{R}$ is a positive-definite kernel, $\mathbf{c}_i \in \mathbb{R}^p$ are coefficient vectors, $\mathbf{A} \in \mathbb{R}^{p \times d}$, and $\mathbf{b} \in \mathbb{R}^p$ define the affine component.

A common choice is a radial basis function kernel

$$K(\mathbf{y}, \mathbf{y}') = \phi\big(\|\mathbf{y} - \mathbf{y}'\|\big), \tag{2.2}$$

where $\phi$ is a polyharmonic spline function.

The coefficients are obtained by solving the regularized least-squares problem

$$\min_{\psi} \sum_{i=1}^{k} \|\psi(\mathbf{y}_i) - \mathbf{x}_i\|^2 + \lambda \, \|\psi\|_{\mathcal{H}}^2, \tag{2.3}$$

where $\|\psi\|_{\mathcal{H}}$ denotes the reproducing kernel Hilbert space (RKHS) norm associated with $K$, and $\lambda \geq 0$ controls the smoothness of the mapping.

##### 2.2 Linear system formulation

Let $K \in \mathbb{R}^{k \times k}$ denote the kernel matrix with entries

$$K_{ij} = K(\mathbf{y}_i, \mathbf{y}_j), \quad P = [\mathbf{1}, \mathbf{Y}] \in \mathbb{R}^{k \times (d+1)}.$$

The spline coefficients are obtained by solving the block linear system

$$\begin{bmatrix} K + \lambda I_k & P \\ P^\top & 0 \end{bmatrix} \begin{bmatrix} C \\ V \end{bmatrix} = \begin{bmatrix} X \\ 0 \end{bmatrix}, \tag{2.4}$$

where $C \in \mathbb{R}^{k \times p}$ contains the kernel coefficients, $V \in \mathbb{R}^{(d+1) \times p}$ contains the affine parameters, and $X \in \mathbb{R}^{k \times p}$ contains the target expression values.

For overdetermined settings, the system can equivalently be written as a ridge regression problem

$$\Theta = (\tilde{K}^\top \tilde{K} + \lambda I)^{-1} \tilde{K}^\top X, \quad \tilde{K} = [K, P], \tag{2.5}$$

where $\Theta$ stacks the kernel and affine coefficients.

#### 2.3 Smoothing parameter selection

The parameter $\lambda$ controls the effective degrees of freedom (DoF) of the fitted surface. Given the hat matrix

$$H(\lambda) = \tilde{K}(\tilde{K}^\top \tilde{K} + \lambda I_m)^{-1}\tilde{K}^\top, \tag{2.6}$$

the DoF is defined as $\mathrm{Tr}(H(\lambda))$.

Using the singular values $\{\sigma_i\}$ of $\tilde{K}$, this can be expressed as

$$\mathrm{Tr}(H(\lambda)) = \sum_i \frac{\sigma_i^2}{\sigma_i^2 + \lambda}. \tag{2.7}$$

We choose $\lambda$ by minimizing the generalized cross-validation (GCV) score

$$\mathrm{GCV}(\lambda) = \frac{\|X - \hat{X}\|_F^2}{(N - \mathrm{Tr}(H(\lambda)))^2}, \tag{2.8}$$

where $\hat{X} = H(\lambda)X$ denotes the fitted values. This follows classical spline smoothing formulations [1]. In practice, we select $\lambda$ at the knee of the GCV curve to balance fidelity and smoothness.

This spline reconstruction provides a differentiable surface linking embedding and gene space, forming the foundation for tangent space computation, velocity projection, and downstream geometric analyses in FlowMap.

#### 2.4 Tangent Space and Analytic Derivatives of the Manifold

We derive analytic expressions for the Jacobian and Hessian of the spline-based manifold mapping. These quantities define the local tangent space and higher-order geometric structure used throughout FlowMap.

**General spline formulation.** Let $\psi : \mathbb{R}^d \to \mathbb{R}^p$ be a vector-valued spline of the form

$$\psi(\mathbf{y}) = \sum_{i=1}^{k} \mathbf{c}_i \, \phi(r_i) + A\mathbf{y} + b, \quad \text{where} \quad r_i = \|\mathbf{y} - \mathbf{y}_i\|.$$

**Jacobian.** By the chain rule, the gradient of each kernel term is

$$\nabla_{\mathbf{y}} \phi(r_i) = \phi'(r_i) \, \frac{\mathbf{y} - \mathbf{y}_i}{r_i}.$$

Thus, the Jacobian $\mathbf{J}_\psi(\mathbf{y}) \in \mathbb{R}^{p \times d}$ is

$$\mathbf{J}_\psi(\mathbf{y}) = \sum_{i=1}^{k} \mathbf{c}_i \, (\nabla_{\mathbf{y}} \phi(r_i))^\top + A.$$

Equivalently, for each component $\alpha = 1, \ldots, p$,

$$\frac{\partial \psi_\alpha}{\partial y_j}(\mathbf{y}) = \sum_{i=1}^{k} c_{i,\alpha} \, \phi'(r_i) \, \frac{y_j - y_{i,j}}{r_i} + A_{\alpha j}.$$

**Hessian.** Differentiating again, the Hessian of each coordinate is obtained via

$$\nabla^2_{\mathbf{y}} \phi(r_i) = \phi''(r_i) \frac{(\mathbf{y} - \mathbf{y}_i)(\mathbf{y} - \mathbf{y}_i)^\top}{r_i^2} + \phi'(r_i) \left( \frac{I}{r_i} - \frac{(\mathbf{y} - \mathbf{y}_i)(\mathbf{y} - \mathbf{y}_i)^\top}{r_i^3} \right).$$

Thus,

$$\frac{\partial^2 \psi_\alpha}{\partial y_p \partial y_q}(\mathbf{y}) = \sum_{i=1}^{k} c_{i,\alpha} \left[ \nabla^2_{\mathbf{y}} \phi(r_i) \right]_{pq}.$$

Collecting all coordinates yields a tensor

$$H(\mathbf{y}) \in \mathbb{R}^{p \times d \times d}.$$

**Thin-plate spline as a special case.** For the thin-plate spline kernel in $d = 2$,

$$\phi(r) = r^2 \log r,$$

we have

$$\phi'(r) = 2r \log r + r, \quad \phi''(r) = 2 \log r + 3.$$

Substituting into the general expressions yields

$$\nabla_{\mathbf{y}} \phi(r_i) = (2 \log r_i + 1)(\mathbf{y} - \mathbf{y}_i),$$

and

$$\frac{\partial^2 \psi_\alpha}{\partial y_p \partial y_q}(\mathbf{y}) = \sum_{i=1}^{k} c_{i,\alpha} \left[ \frac{2(y_p - y_{i,p})(y_q - y_{i,q})}{r_i^2} + (2 \log r_i + 1)\, \delta_{pq} \right],$$

recovering the classical TPS expressions used in implementation.

**Derived geometric quantities.** The analytic Jacobian and Hessian enable computation of standard differential-geometric quantities associated with the reconstructed manifold. The metric tensor (first fundamental form) is given by

$$g_{pq}(x) = \sum_{\alpha=1}^{D} \frac{\partial f_\alpha}{\partial x_p} \frac{\partial f_\alpha}{\partial x_q} = [J_f(x)^\top J_f(x)]_{pq}.$$

The Christoffel symbols (second kind) are defined as

$$\Gamma^k_{pq}(x) = g^{kl}(x) \, \langle H_{pq}(x), \, \partial_l f(x) \rangle, \quad \text{where} \quad H_{pq}(x) = \frac{\partial^2 f}{\partial x_p \partial x_q}(x).$$

From these, the Riemann curvature tensor, Ricci curvature, and scalar curvature follow:

$$R_{pqrs}(x) = \sum_A \left( h^A_{pr} h^A_{qs} - h^A_{ps} h^A_{qr} \right), \quad \mathrm{Ric}_{qs}(x) = g^{pr}(x) R_{pqrs}(x), \quad R(x) = g^{qs}(x) \mathrm{Ric}_{qs}(x),$$

where $h^A_{pq} = \langle H_{pq}(x), n^A(x) \rangle$ denotes the second fundamental form in each normal direction.

These expressions establish the connection between the spline-based manifold reconstruction and its intrinsic geometric structure. While not directly used in the main algorithm, they provide a foundation for geometric analysis of the learned embedding.

### 3 Curvature of embedded flow curves

#### 3.1 Acceleration decomposition and curvature of embedded flow curves

**Case 1: Flat space, constant speed, curved flow.** Consider a smooth trajectory

$$\gamma(t) \in \mathbb{R}^d$$

with velocity

$$\dot{\gamma}(t) = \frac{d\gamma(t)}{dt}$$

and acceleration

$$\ddot{\gamma}(t) = \frac{d^2\gamma(t)}{dt^2}.$$

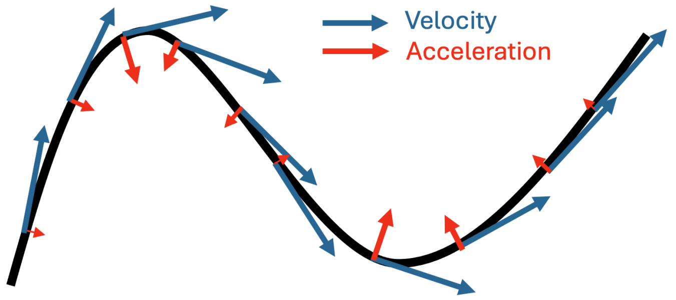


Figure 1: *Acceleration and curvature for constant-speed motion in flat space. Under constant-speed motion, the acceleration vector is perpendicular to the velocity vector and points toward the local center of curvature. Regions with larger curvature exhibit larger acceleration magnitude, reflecting more rapid turning of the trajectory.*

Assume the trajectory evolves with constant speed,

$$\|\dot{\gamma}(t)\| = c,$$

while the direction of motion changes over time. Define the unit tangent vector

$$T(t) = \frac{\dot{\gamma}(t)}{\|\dot{\gamma}(t)\|}.$$

The curvature of the trajectory is defined as

$$\kappa(t) = \left\| \frac{dT}{ds} \right\|,$$

where $s$ denotes arc length along the curve. Curvature therefore measures how rapidly the direction of the trajectory changes per unit distance traveled.

In flat Euclidean space with constant speed, the acceleration vector is entirely orthogonal to the direction of motion and satisfies

$$\ddot{\gamma}(t) = c^2 \frac{dT}{ds}.$$

Consequently,

$$\|\ddot{\gamma}(t)\| = c^2\kappa(t).$$

Thus, under constant-speed motion, acceleration arises purely from curvature of the trajectory: the trajectory bends even though its speed remains unchanged.

**Case 2: Flat space, varying speed, curved flow.** We now consider a trajectory

$$\gamma(t) \in \mathbb{R}^d$$

whose speed and direction both vary over time. Define the unit tangent vector

$$T(t) = \frac{\dot{\gamma}(t)}{\|\dot{\gamma}(t)\|}.$$

The acceleration then decomposes as

$$\underbrace{\ddot{\gamma}(t)}_{\text{total acceleration}} = \underbrace{\frac{d}{dt}\|\dot{\gamma}(t)\|\,T(t)}_{\text{flow acceleration}} + \underbrace{\|\dot{\gamma}(t)\|^2\frac{dT}{ds}}_{\text{steering acceleration}}.$$

The first term is parallel to the velocity vector and represents acceleration or deceleration along the trajectory. The second term is orthogonal to the velocity vector and captures turning of the flow through the curvature

$$\kappa(t) = \left\|\frac{dT}{ds}\right\|.$$

This decomposition closely parallels the flow/steering decomposition introduced in the main text. The tangential term corresponds to the flow acceleration $a_{\text{flow}}$, while the orthogonal term corresponds to the steering component $a_{\text{steer}}$.

Importantly, curvature is a normalized geometric quantity that measures directional change independently of speed. The steering acceleration scales as

$$\|a_{\text{steer}}(t)\| = \|\dot{\gamma}(t)\|^2\kappa(t),$$

showing that the same geometric curvature produces larger steering acceleration when the trajectory moves more rapidly through the state space. Consequently, curvature reflects the intrinsic geometry of the trajectory itself, whereas the steering acceleration additionally depends on how quickly the system traverses that trajectory.

**Case 3: Curved space, constant speed, geodesic flow.** We now consider trajectories constrained to a curved manifold [2]

$$\mathcal{M} \subset \mathbb{R}^p.$$

Let

$$\psi : \mathbb{R}^d \to \mathbb{R}^p$$

denote a smooth parameterization of the manifold, and let

$$y(t) = (y_1(t), \dots, y_d(t))$$

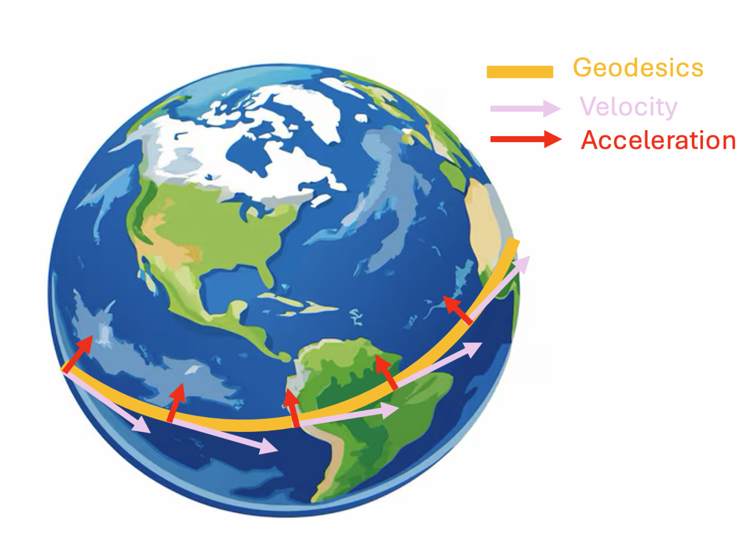


Figure 2: *Geodesic motion on a curved manifold under constant-speed flow. Although the trajectory is intrinsically straight on the manifold* ($a_{\mathrm{steer}} = 0$)*, the acceleration remains nonzero because the manifold itself is curved in the ambient space. For a sphere, the acceleration vectors have approximately constant magnitude and point normal to the surface, reflecting extrinsic curvature of the manifold rather than intrinsic turning of the flow.*

be the intrinsic coordinates of the trajectory. The ambient trajectory is

$$\gamma(t) = \psi(y(t)).$$

Using the coordinate basis induced by the embedding,

$$\partial_i \psi(y) = \frac{\partial \psi}{\partial y_i}(y), \qquad i = 1, \dots, d,$$

the velocity becomes

$$\dot{\gamma}(t) = \sum_{i=1}^{d} \dot{y}_i(t)\, \partial_i \psi(y(t)).$$

The acceleration decomposes into tangent and surface components:

$$\underbrace{\ddot{\gamma}(t)}_{\text{total acceleration}} = \underbrace{\sum_{i=1}^{d} \ddot{y}_i(t)\, \partial_i \psi(y(t))}_{a_{\mathrm{tangent}}(t)} + \underbrace{\sum_{i,j=1}^{d} \dot{y}_i(t) \dot{y}_j(t)\, \partial^2_{ij} \psi(y(t))}_{a_{\mathrm{surface}}(t)}.$$

Here

$$\partial^2_{ij} \psi(y) = \frac{\partial^2 \psi}{\partial y_i \partial y_j}(y)$$

denotes the Hessian of the manifold parameterization. Equivalently, the surface term can be written using the Hessian operator:

$$H_{\psi(y(t))}[\dot{y}(t), \dot{y}(t)] = \sum_{i,j=1}^{d} \dot{y}_i(t) \dot{y}_j(t)\, \partial^2_{ij} \psi(y(t)).$$

The tangential component $a_{\text{tangent}}(t)$ captures intrinsic acceleration along the manifold and can be further decomposed into

$$a_{\text{tangent}}(t) = a_{\text{flow}}(t) + a_{\text{steer}}(t),$$

where $a_{\text{flow}}(t)$ measures acceleration along the direction of motion and $a_{\text{steer}}(t)$ measures turning within the manifold.

For constant-speed geodesic flow,

$$a_{\text{flow}}(t) = 0, \qquad a_{\text{steer}}(t) = 0,$$

so that

$$a_{\text{tangent}}(t) = 0.$$

Consequently,

$$\ddot{\gamma}(t) = a_{\text{surface}}(t),$$

meaning that all remaining acceleration arises purely from bending of the manifold within the ambient gene-expression space.

The curvature of the ambient trajectory is obtained by normalizing the acceleration by speed:

$$\kappa(t) = \frac{\|\ddot{\gamma}(t)\|}{\|\dot{\gamma}(t)\|^2} = \frac{\|a_{\text{surface}}(t)\|}{c^2}, \qquad \|\dot{\gamma}(t)\| = c.$$

Thus, for constant-speed geodesic motion on a curved manifold, curvature arises entirely from the extrinsic geometry of the embedded surface rather than from steering of the flow within the manifold.

**Case 4: Curved space, general flow.** In the general setting, the trajectory

$$\gamma(t) = \psi(y(t))$$

may both change speed and turn within a curved manifold. The acceleration decomposes into tangent and surface components:

$$\underbrace{\ddot{\gamma}(t)}_{\text{total acceleration}} = \underbrace{\sum_{i=1}^{d} \ddot{y}_i(t)\,\partial_i\psi(y(t))}_{a_{\text{tangent}}(t)} + \underbrace{\sum_{i,j=1}^{d} \dot{y}_i(t)\dot{y}_j(t)\,\partial_{ij}^2\psi(y(t))}_{a_{\text{surface}}(t)}.$$

Let

$$e_1(t) = \frac{\dot{\gamma}(t)}{\|\dot{\gamma}(t)\|}$$

denote the unit flow direction, and let

$$\{e_1(t), e_2(t), \ldots, e_d(t)\}$$

be an orthonormal basis of the tangent space $T_{\gamma(t)}\mathcal{M}$. Then the tangential acceleration can be expanded as

$$a_{\text{tangent}}(t) = \underbrace{\langle a_{\text{tangent}}(t), e_1(t)\rangle e_1(t)}_{a_{\text{flow}}(t)} + \underbrace{\sum_{i=2}^{d}\langle a_{\text{tangent}}(t), e_i(t)\rangle e_i(t)}_{a_{\text{steer}}(t)}.$$

Therefore, the full acceleration decomposes as

$$\ddot{\gamma}(t) = a_{\text{flow}}(t) + a_{\text{steer}}(t) + a_{\text{surface}}(t).$$

Here,

$$a_{\text{flow}}(t)$$

measures acceleration or deceleration along the direction of motion,

$$a_{\text{steer}}(t)$$

measures directional turning within the manifold, and

$$a_{\text{surface}}(t)$$

measures bending induced by the embedding of the manifold in ambient gene-expression space.

The curvature of the ambient trajectory is

$$\kappa(t) = \frac{\|\ddot{\gamma}_{\perp}(t)\|}{\|\dot{\gamma}(t)\|^2},$$

where

$$\ddot{\gamma}_{\perp}(t) = a_{\text{steer}}(t) + a_{\text{surface}}(t)$$

denotes the component of the acceleration orthogonal to the flow direction.

Normalization by $\|\dot{\gamma}(t)\|^2$ ensures invariance to time reparameterization of the trajectory, so that curvature depends only on the geometry of the path rather than the speed at which it is traversed.

Because $a_{\text{steer}}(t)$ and $a_{\text{surface}}(t)$ are orthogonal,

$$\kappa(t)^2 = \frac{\|a_{\text{steer}}(t)\|^2 + \|a_{\text{surface}}(t)\|^2}{\|\dot{\gamma}(t)\|^4}.$$

We can equivalently define component-wise curvature contributions:

$$\kappa_{\text{steer}}(t) = \frac{\|a_{\text{steer}}(t)\|}{\|\dot{\gamma}(t)\|^2}, \qquad \kappa_{\text{surface}}(t) = \frac{\|a_{\text{surface}}(t)\|}{\|\dot{\gamma}(t)\|^2}.$$

Then

$$\kappa(t)^2 = \kappa_{\text{steer}}(t)^2 + \kappa_{\text{surface}}(t)^2.$$

The steering curvature measures directional turning within the manifold, whereas the surface curvature measures bending induced by the embedding of the reconstructed manifold in ambient gene-expression space.

The flow component $a_{\text{flow}}(t)$ instead quantifies acceleration or deceleration along the trajectory and therefore affects traversal speed but not geometric curvature.

In practice, RNA-velocity visualizations primarily expose the flow and steering components, whereas the surface component becomes identifiable only through an explicit differentiable reconstruction of the embedding manifold.

##### 3.2 Relationship to intrinsic and extrinsic curvature

The decomposition developed above is closely related to the distinction between intrinsic and extrinsic geometry in differential geometry. Consider the constant-speed geodesic case discussed in Case 3, where

$$\gamma(t) = \psi(y(t))$$

is a trajectory on a smooth embedded manifold

$$\mathcal{M} \subset \mathbb{R}^p$$

satisfying

$$a_{\text{flow}}(t) = 0, \qquad a_{\text{steer}}(t) = 0.$$

In this setting, the acceleration reduces to

$$\ddot{\gamma}(t) = a_{\text{surface}}(t) = \sum_{i,j=1}^{d} \dot{y}_i(t)\dot{y}_j(t)\, \partial_{ij}^2 \psi(y(t)).$$

The second-derivative term

$$\partial_{ij}^2 \psi(y) = \frac{\partial^2 \psi}{\partial y_i \partial y_j}(y)$$

describes how the tangent space changes as one moves along the manifold. Collectively, these second derivatives form the Hessian of the embedding map,

$$H_{\psi(y)}[\dot{y}, \dot{y}],$$

which measures the local bending of the embedded surface in the ambient space.

From the viewpoint of intrinsic geometry, a geodesic trajectory is locally straight: it has no steering acceleration within the manifold itself. However, when the manifold is embedded in a higher-dimensional ambient space, the trajectory may still appear curved due to bending of the surface. The resulting acceleration is therefore entirely extrinsic.

The link to Riemann curvature can be written through the Gauss equation [2]. Let

$$B_{ij}(y) = \Pi_\perp \left[\partial_{ij}^2 \psi(y)\right]$$

denote the normal component of the Hessian of the embedding map, where $\Pi_\perp$ is projection onto the normal space of the surface. Then the Riemann curvature tensor of the embedded surface satisfies

$$R_{ijkl}(y) = \langle B_{ik}(y), B_{jl}(y) \rangle - \langle B_{il}(y), B_{jk}(y) \rangle \,.$$

Thus, the intrinsic curvature of the learned surface is determined by how the Hessian of $\psi$ bends the tangent space into the ambient gene-expression space.

This distinction parallels the classical separation between intrinsic and extrinsic curvature in Riemannian geometry. Intrinsic curvature describes geometric properties that can be measured entirely within the manifold itself, independent of how the manifold is embedded. Extrinsic curvature instead depends on how the manifold bends inside the ambient space. In FlowMap, the surface acceleration

$$a_{\text{surface}}(t)$$

captures this extrinsic bending directly through the Hessian of the reconstructed embedding map.

Consequently, even when a cellular trajectory follows an intrinsic geodesic of the learned state-space manifold, the trajectory may still exhibit substantial curvature in gene-expression space due to the geometry of the embedding itself.

### References


[1] Grace Wahba. *Spline models for observational data*. SIAM, 1990.

[2] Manfredo P Do Carmo. *Differential geometry of curves and surfaces: revised and updated second edition*. Courier Dover Publications, 2016.
